# Global patterns of genetic admixture reveal effects of language contact

**DOI:** 10.1101/2024.12.19.629340

**Authors:** Anna Graff, Damián E. Blasi, Erik J. Ringen, Vladimir Bajić, Daphné Bavelier, Kentaro K. Shimizu, Brigitte Pakendorf, Chiara Barbieri, Balthasar Bickel

**Affiliations:** Department of Comparative Language Science, University of Zurich; Zurich, 8050, Switzerland; Department of Evolutionary Biology and Environmental Studies, University of Zurich; Zurich, 8057, Switzerland; Center for the Interdisciplinary Study of Language Evolution (ISLE), University of Zurich; Zurich, 8057, Switzerland; Catalan Institute for Research and Advanced Studies (ICREA); Barcelona, 08010, Spain; Center for Brain and Cognition, Pompeu Fabra University; Barcelona, 08010, Spain; Human Biology and Primate Evolution, Freie Universität Berlin; Berlin, 14195, Germany; Faculté de Psychologie et des Sciences de l’Education, Université de Genève; Geneva, 1202, Switzerland; Kihara Institute for Biological Research, Yokohama City University; Yokohama, 244-0813, Japan; Laboratoire Dynamique Du Language, CNRS & Université Lumière Lyon 2; Lyon, 69007, France; Dipartimento di Scienze della vita e dell’ambiente, Università degli Studi di Cagliari; Cagliari, 09124, Italy

## Abstract

When speakers of different languages are in contact, they often borrow features like sounds, words, or syntactic patterns from one language to the other, but the lack of historical data has hampered estimation of this effect at a global scale. We break out of this impasse by using genetic admixture as a proxy for population contact. We find that language pairs whose speaker populations underwent genetic admixture or that are located in the same geo-historical area share more features than others, suggesting borrowing. The effect varies strongly across features, partly following expectations from differences in lifelong learnability, partly responding to differences in social imbalances during contact. Additionally, we find that for some features, admixture decreases sharing. This likely reflects signals of divergence (schismogenesis) under contact.

---

Unlike their genes, humans transfer cultural traits not only through vertical inheritance but also through horizontal borrowing, also referred to as copying, spread, diffusion, or convergence in linguistics (*1–5*). When populations are in contact with each other, they often adopt or borrow cultural elements such as technologies, beliefs, practices, and names for these, be it by choice or imposition (*6*). For example, mobile phones and words for them have rapidly spread across the world in the past thirty years or so (*7*). Such instances of horizontal transfer are made possible by the remarkable learning abilities that characterize humans over the entire lifespan. In the case of language, these abilities can result in various extents of bilingualism and multilingualism, patterns which provide particularly fertile grounds for borrowing (*8*).

However, the extent of such borrowing remains heavily debated, with far-reaching consequences on the validity of the tree model for linguistic evolution (*9–16*). On the one hand, core vocabulary is remarkably persistent in vertical transmission and tends to resist borrowing. Many words are inherited with only slight modification all the way from the root of a language phylogeny to large swathes of its descendants (e.g., the words for *mother*, *madre*, *Mutter*, and *mère* in the Indo-European daughter languages English, Spanish, German and French, respectively) so that relationships between languages and groups of languages can readily be detected and reconstructed (*17*, *18*). On the other hand, features of linguistic structure, such as patterns in grammar or phonology, tend to be more volatile over time (*19–21*). The distribution of these features, as well as the introduction of new concepts and words, is often parsimoniously explained by borrowing between specific pairs of populations (*22*) or between several populations within a wider geo-historical area (*23*, *24*, *3*, *25–29*).

Borrowing is typically initiated by speakers who have already acquired the basics of their native language; younger children tend to be overly resilient and conservative learners, and they mostly lack the social power to initiate change (*30–34*). As a result, borrowing is expected to be more likely in features that remain relatively easy to learn after early childhood, for example, new lexical concepts are expected to remain easier to learn than new grammatical categories (*35–38*). These expectations largely match observations about differences in borrowability from case studies (*3*, *5*), but it is unknown to what extent they hold globally and to what extent borrowing instead depends more on specific social histories, such as imbalances of power or demography when populations are in contact.

While frequent, borrowing is not the only outcome of contact. Anthropologists have long noted that contact can also lead to the opposite, viz. *schismogenesis*, a process of signaling divergence between populations in contact (*39*). Divergence in language features has often been noted in local case studies (*40–44*). In specific cases, like within Austronesian, contact has been identified as an important driver of diversification in vocabulary, but less so in grammar (*19*) and only partially so in phonology (*45*). At a global scale, divergence has been systematically quantified only for dialect contact (*46*), finding features of grammatical form to diverge most, and for patterns in language phylogenies inferred from core vocabulary data (*47*), finding bursts of change after language splits.

While language contact enjoys substantial attention, its global study has suffered from severe biases (*48*). Most importantly, research has focused on post-hoc inferences, where the current distribution of some features is most plausibly explained by borrowing during past contact. The evidence for such contact is sometimes independently supported by geo-historical evidence, but in most parts of the world this evidence is largely circumstantial. As a result, the extent to which contact shapes linguistic evolution, the degree to which features differ in their probability to be borrowed, and the net balance between borrowing and divergence under contact remain poorly understood.

Here, we introduce genetic admixture as an alternative proxy of population contact to study its effects on language. We quantify the effects of genetic contact on language structure and ask whether genetic contact and shared geo-cultural history (to the extent known) yield similar linguistic outcomes. We furthermore compare these effects across specific linguistic features and assess differences in borrowability as expected from case studies and research on learnability over the lifespan.

## Sampling contact between populations

The motivation to capture contact through genetics relies on the effects of human population admixture. The intense demographic contact leading to an admixed population profile provides ample opportunity for contact affecting not only genes, but a wide range of cultural traits, including language. Indeed, genes and languages are sometimes transferred together (*49*), and together with languages, so are linguistic features. For instance, the borrowing of click sounds from Khoisan into Bantu languages in Zambia was coupled with demographic exchange between distant groups after large-scale migrations (*50*, *51*).

To probe for genetic evidence of contact, we searched for pair-wise contact between genetic ancestries that were sufficiently divergent to be distinguishable. In particular, we identified populations with admixture from one ancestry component that is associated with a linguistically unrelated group (Fig. 1A, Materials and Methods). This procedure ensures consistency in the criteria for genetic contact, and it keeps our sample relatively free of confounds from shared linguistic inheritance. However, our procedure is limited by the non-uniform coverage in our sample across the world (Fig. 1B) and by having enough samples to represent each ancestry (*52*). A further limitation is that we cannot cover the impact of cases of pairwise admixture in genetic ancestries underrepresented in our sample, of gradients of admixed ancestries (*53*), and of admixture scenarios with more than two ancestries. Finally, the restriction to unrelated language pairs might underestimate both borrowing and divergence effects because these are particularly expected within language families, where structural similarity boosts borrowing (*3*, *54–56*, *16*) and shared history boosts divergence (*40*, *46*).

**Fig. 1.**
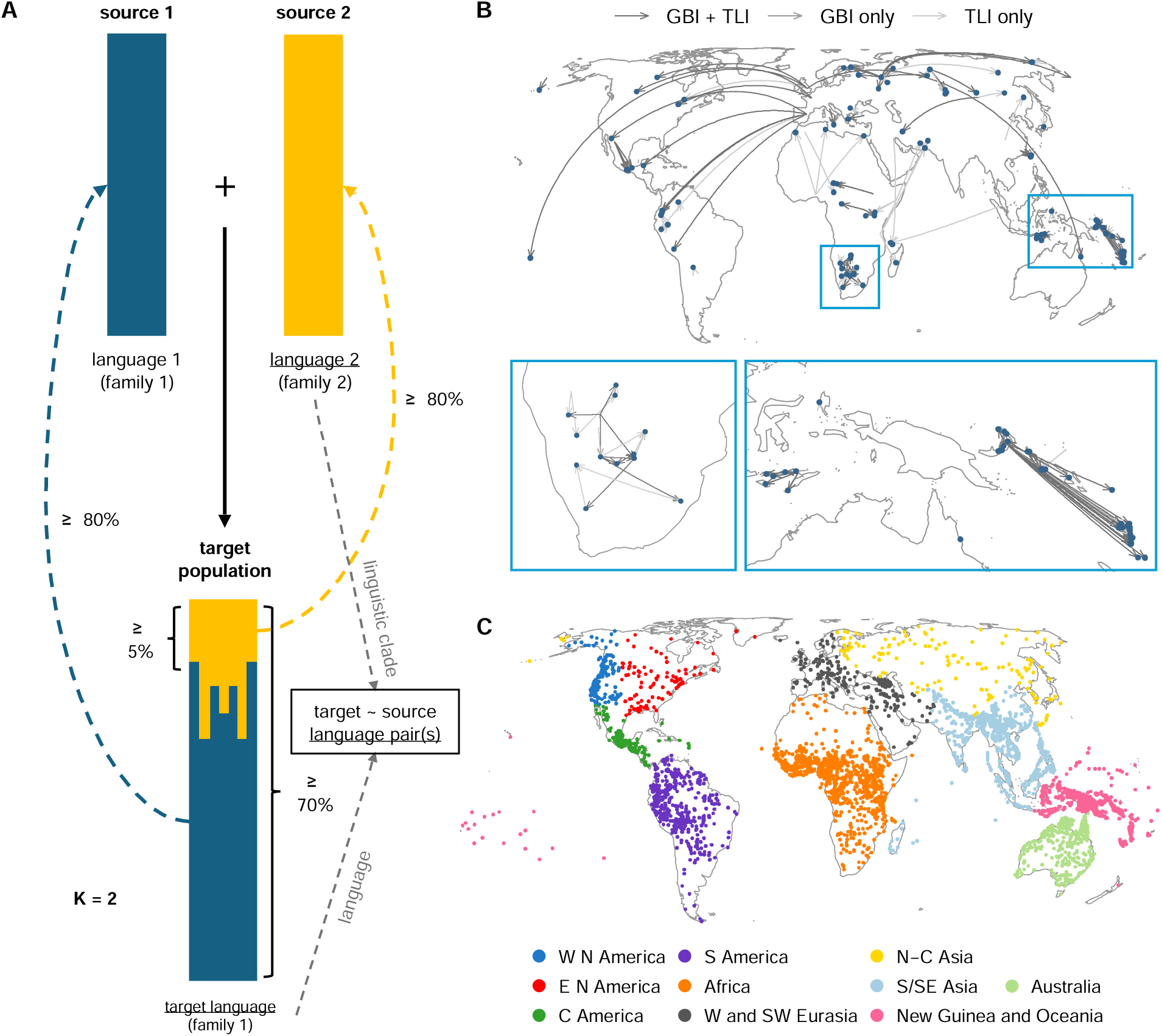
Sampling contact between populations. (**A**) Sampling admixture in populations (solid arrow) if their two largest ancestry components amount to at least 70% of genetic ancestry, of which the minor source contributes at least 5%, and if the admixture is evident through at least 5 different levels of globally assumed components (*K*, tested from 12 to 30, see also fig. S1A for results at *K* = 12). The two components were associated (dashed arrows) with source populations exhibiting the ancestry to at least 80%, using *F*_ST_ distances to the target as a guide. Target populations were only considered if their two main ancestries were assigned to populations (source 1, source 2) speaking languages (language 1, language 2) from different language families (family 1, family 2), whereby one of these families (family 1 in this illustration) should be the family of the target language spoken by the target population. Targets were associated (grey arrows) with their currently spoken language (‘language 1’ in the figure) and the source with the phylogenetic clade that best characterizes the language varieties (‘language 2’) at the time of contact. To control for shared inheritance we only sampled pairs where target and source are from different families (family 1 and 2). (**B**) 126 target-source language pairs. Blue: target languages, yellow: source clades (centered on one language for visualization purposes only); TLI and GBI: different linguistic datasets (see main text). (**C**) Languages from which features were drawn colored by geo-historical area from AUTOTYP (fig. S1B for an alternative).

To identify populations with admixture, we ran ADMIXTURE (*57*) on genomic SNP chip data from an expanded version of GeLaTo (*49*) (4768 individuals in 558 populations associated with 373 languages; table S1), a database that matches genetic populations to their languages, and supplemented the results with cases of admixture reported in the literature (fig. S1A, Materials and Methods).

The resulting list of genetically admixed populations covers different geographical and historical scales, including local cases of demic contact, larger Neolithic displacements of farmers and pastoralist groups (*58*), and intercontinental contacts from the past five centuries of population movements (invasions, displacements) associated with European colonialism (*59*) and slave trade (*60–62*).

From this list we derived target-source pairs (N=126, table S2) and associated them with languages. For the target, representing a present-day population, we followed the association provided by GeLaTo and the additional literature. For the source population, representing a past population, we have no access to the language(s) with which it was associated at the time of contact. In response, we resorted to the nearest clade of the languages currently spoken by present-day proxies for the past source population, assuming that this includes most (often all) of the likely features reconstructible for contact time (Materials and Methods). For example, for the Quechua-Spanish pair, we sampled the source feature states from the Romance clade to approximate the 16^th^ century dialect variation at the time when contact began; solely considering modern standard Castillian Spanish would artificially reduce this variation.

We then linked target and source languages to their features from two databases curated to remove logical and strong statistical dependencies between features (*63*): GBI (“Grambank Independent”) covering grammatical features, and TLI (“Typology Linked and Independent”) covering a variety of grammatical, lexical, and phonological features (table S3). To control for universal baseline expectations about the presence of features in the target and source languages, we further sampled 300 random pairs of unrelated languages. We extracted the data separately for every state in features with more than two states, resulting in a total of N=638 feature states (N=202 in GBI, N=481 in TLI).

This procedure focuses on pairwise contact. However, contact can also characterize entire networks of languages that jointly evolved in a given geo-historical area (*3*, *24*, *48*, *25*, *26*). Even though the relevant historical and ethnographic evidence is often (as noted) circumstantial, we allow for this alternative scenario of contact by additionally assigning all sampled languages to areas that have been established as particularly prone to contact (Fig. 1C). We source such areas from the AUTOTYP database (*64*), and – for a sensitivity analysis – from Glottolog (*65*, *66*) (fig. S1B) rather than geographical distances between current locations (*67*, *68*), because the areas are less sensitive to recent migrations (*69–71*).

We then modeled the probability of a language pair to share states as a function of genetic admixture and areal co-location in a series of Bayesian multilevel logistic regressions (Suppl. Text, figs. S12–S29, tables S4–S21). Model comparison with approximate leave-one-out cross-validation suggests that a model with both genetic and areal predictors strongly outperforms models with only one of these predictors (fig S2, all Δ_elpd_>260 + 46).

## Genetic contact and areas increase the probability of sharing feature states

The best-fitting model shows that both genetically and areally defined contact increases state sharing, suggesting a global effect of borrowing under contact (Fig. 2A). Genetic contact increases sharing probabilities by a posterior mean of 4.0% in the GBI dataset (89%-HPDI = [1.2%, 5.5%], P(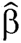>0)=0.99) and 7.3% in the TLI dataset (89%-HPDI = [2.6%, 11.7%]; P(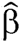>0)=0.99). Belonging to the same area increases sharing probabilities by a posterior mean of 3.4% in both GBI (89%-HPDI = [1.2%, 6.7%], P(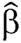>0)=0.99) and TLI (89%-HPDI = [2.2%, 4.9%], P(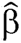>0)=1.00). All results are robust against the alternative area definition from Glottolog (fig. S3). The effects are relatively strong given that the baseline expectations are well above chance (i.e., a 50% probability of sharing a state) and leave only limited room for probability increases (with all posterior mean sharing probabilities in the baseline higher than 64.0%; 89%-HPDI = [64.0%, 68.1%] for GBI and 89%-HPDI = [72.5%, 75.8%] for TLI). Especially in the case of genetic contact, we find strong evidence for borrowing (HPD > 89%) even in cases where the universal baseline probability of sharing is very high and the effects therefore remain small.

**Fig. 2.**
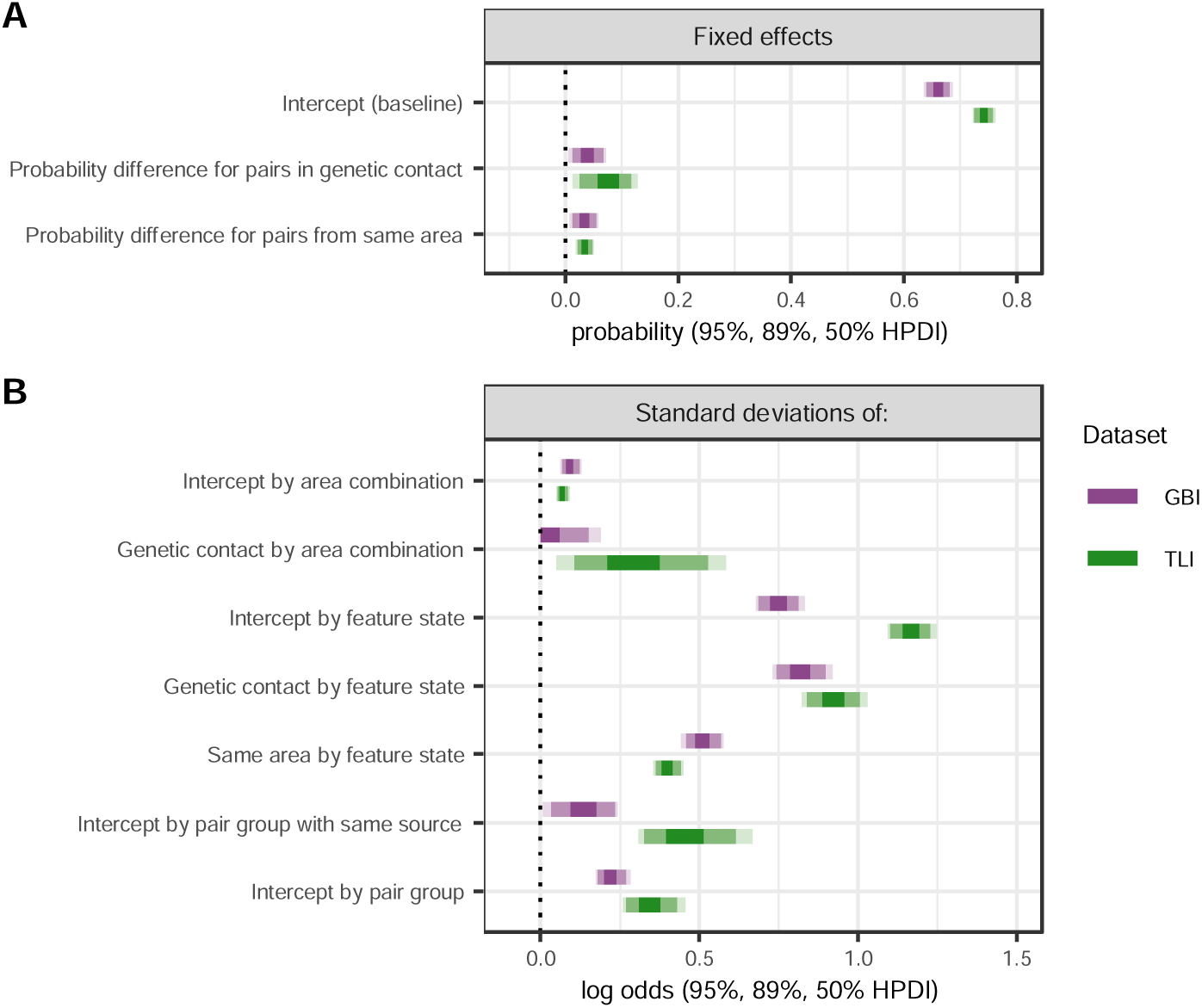
Posterior predictions of the best-fitting model. (**A**) The probability of sharing feature states decisively increases in contact compared to a random baseline (ΔP>0 in over 99% of the posterior probabilities), across two linguistic datasets (GBI focusing on grammatical and lexical features, TLI additionally including phonological features) but (**B**) with substantial standard deviations by feature state.

## Effects of contact vary across feature states and contact types

The different effect sizes of genetic contact between the GBI and TLI datasets suggest a fair amount of variation driven by the specific features coded in each. This is confirmed by the relatively large standard deviations of feature states (Fig. 2B, all posterior mean *SD* > 0.40±0.06 in log odds and two standard errors). Fig. 3 shows the estimated effects in terms of probability differences for all 683 states (for details, see tables S22–S23). In addition to states with excess sharing under contact, i.e., borrowing (positive differences; 34% and 28% of states in genetic contact for GBI (Fig. 3A) and TLI (Fig. 3C) respectively; 28% and 15% of states in areal contact for GBI (Fig. 3D) and TLI (Fig. 3F) respectively), there is a substantial number of states that are unaffected by contact (with zero included in the 89% highest posterior density interval; 48% and 64% of states in genetic contact for GBI and TLI respectively; 65% and 83% of states in areal contact for GBI and TLI, respectively). Additionally, to a lesser but still noticeable extent, there are states with decreased sharing probabilities, i.e., cases of divergence under genetic contact (18% of states in GBI, 8% in TLI, Fig. 2C). Under areally defined contact, divergence effects are much rarer (7% of states in GBI, 2% in TLI, Fig. 2B). This might reflect the fact that divergence is intrinsically difficult to discover in areas because the total proportion of shared states in an area can be high even if most geographically adjacent languages show in fact local divergence.

**Fig. 3.**
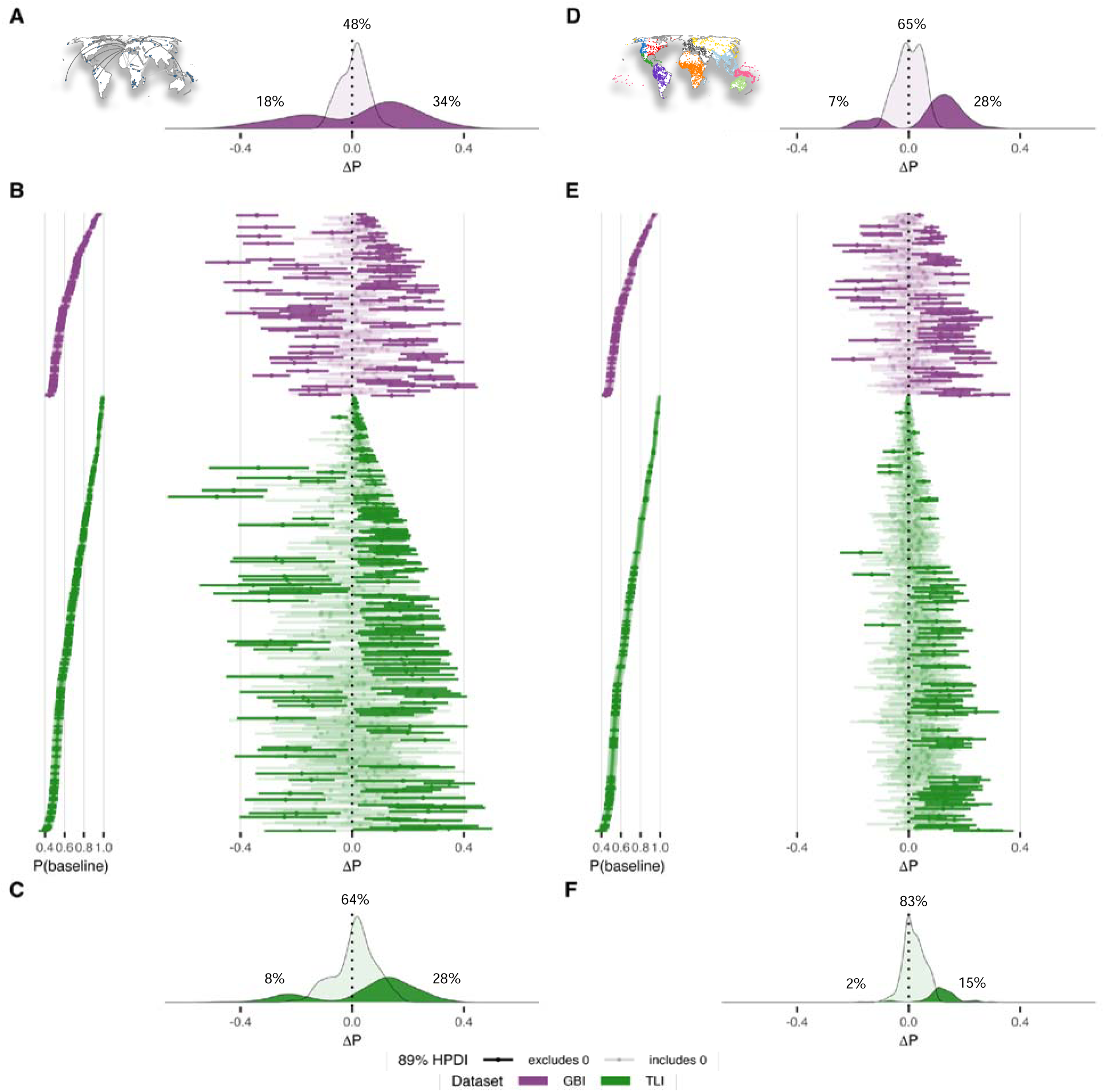
Posterior baseline sharing probability per feature state and posterior ΔP for each feature state under contact (ordered by dataset and effect size). (**A–C**) As a result of genetic admixture vs. (**D–F**) co-location in a geo-historical area. For alternatively ordered visualizations of posterior ΔP, see fig. S6A. (**A**) and (**C**) show density plots of states with 89% HPD excluding vs. including zero in GBI and TLI under genetic contact, (**D**) and (**F**) under areal co-location. (**B**) and (**E**) are interval plots per feature state, ordered by increasing posterior baseline. For details see tables S21–23. There are more features with decreases in sharing probability (ΔP<0) in genetic than areal contact (density plots of features with 89%-HPD excluding vs. including 0), suggesting specific effects of divergence in addition to borrowing. Although states with higher baseline sharing probabilities have less scope for increased rates of borrowing under contact, we detect borrowing for states across the range of baseline sharing probabilities.

The total distribution inside an area might also influence the chances of detecting borrowing under genetic contact when sampling pairs from within that area (e.g., Bantu and Khoisan languages) as opposed to between different areas (e.g., Iberian and Quechuan languages). This possibility is particularly important because the difference between the two scenarios tends to correlate with different histories: genetic contact between different areas in our sample predominantly involves cases associated with European colonialism, with more recent timing and stronger demographic imbalances than most other cases of genetic contact, as in the case of Creole languages, (*72*). However, quantifying the effect of genetic contact separately for contact between vs. within areas (fig. S4C–D) revealed the same overall trends as without the distinction (fig. S4A)

The variation among features often shows opposite trends for areal vs. genetic contact: the feature states that show most borrowing in areal contact (maximum: a feature tracing whether speaker and hearer are marked as prefixes and/or as proclitics on verbs or not) show divergence or no difference under all types of genetic contact (global vs. within vs. between areas).

Conversely, the feature state that shows most divergence in areal contact (maximum: a feature tracing the presence of an existential verb like ‘to be’) show borrowing or no difference in genetic contact (fig. S4, table S22). These observations hold beyond the single-most extreme cases of borrowing and divergence, and they are very similar in the sensitivity analysis (figs. S5– S7, table S23). They suggest that specific feature states differ in how likely they are to act as carriers for signaling social divergence vs. convergence in the form of borrowing (*46*).

## Meta-analyses highlight differences between domains of language

Feature states differ in how they react to contact, but do these differences match the borrowability hierarchies proposed on the basis of case studies and/or expectations about the ease of learning new features across the lifespan? To answer these questions, we conducted meta-analyses comparing contact effects across domain classifications given in the GBI and TLI datasets (*63*) (Table 1, table S3). While motivated by considerations of linguistic analysis, they approximate some of the distinctions for which one would expect differences in borrowability and learnability based on previous reports (*3*, *24*, *35–38*, *5*).

**Table 1.** Feature classifications for meta-analyses. Definitions and examples based on Graff et al. (*63*). Note that the GBI dataset does not contain phonological data.

| Name | Definition | Examples |
| --- | --- | --- |
| LEXICAL SEMANTICS | Structuring of lexical meaning | Colexifications, e.g. same word for ‘arm’ and ‘hand’ |
| LEXICAL CLASSES | Formal, morphosyntactically relevant information in the lexicon | Presence of verb classes, grammatical gender |
| GRAMMATICAL CATEGORIES | Presence and nature of semantic notions in grammar (not their concrete expression) | Presence of past tense, evidentiality |
| LINEAR ORDER | Ordering of sentence elements | Position of the object relative to the verb |
| OTHER GRAMMAR | All other formal aspects of grammar | Suffix vs. prefix expression of past tense marking on verbs, type of relative clause formation |
| PHONOLOGY | Patterns and rules in the sound system | (Any of the following) |
| PROSODY | Rhythm, stress and intonation | Stress placement |
| VOCALIC | Features of the vowel system | Vowel nasalization |
| CONSONANTAL | Features of the consonant system | Presence of fricatives |

Our results are only partially consistent with expectations. Most consistent with expectations is our finding that LINEAR ORDER shows similar or higher borrowing probabilities (ΔP>0) compared to other aspects of grammar (Fig. 4A–D; see fig. S8 for details and fig. S10 for the sensitivity analysis). For instance, in GBI, LINEAR ORDER shows a 3% higher median posterior probability of sharing states under genetic contact (Fig. 4A) while there is no evidence for a difference under areal contact (grey-shaded cell, Fig. 4D). This is largely in line with findings from case studies (*3*, *24*), the underuse of linear order for schismogenesis (*46*), and its relatively persistent learnability over age (*73*). However, the effect is not as robust across conditions as one might expect, and it is only supported with high posterior probabilities in the GBI dataset (Fig. 4A–H, fig. S10). Another finding that seems to confirm previous reports (*68*, *74*) is higher or similar borrowing probabilities for CONSONANTAL over VOCALIC features, but strong evidence is again limited to only some conditions (Fig. 4E–H, figs. S9 and S11).

**Fig. 4.**
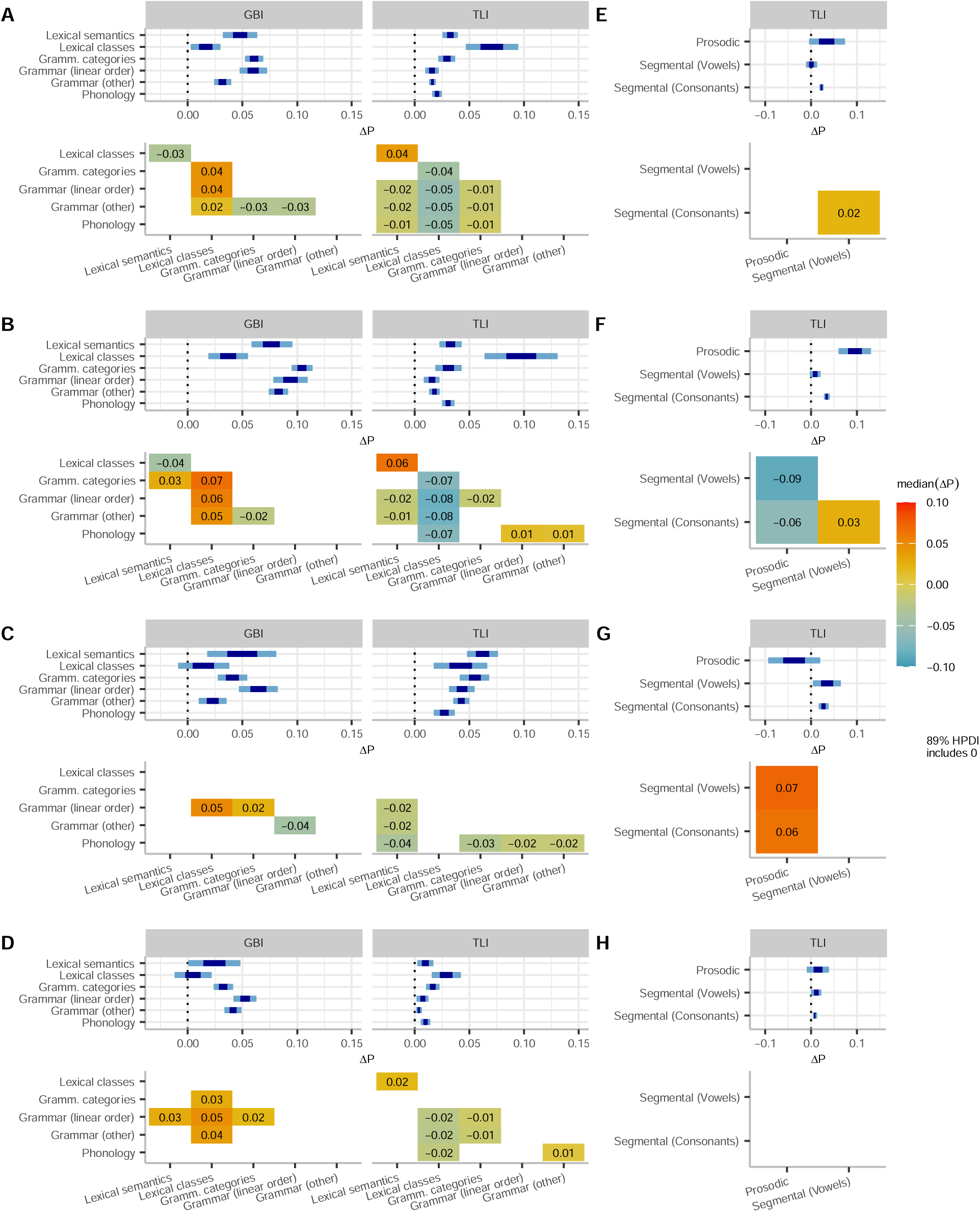
Meta-analyses of contact effects under different types of contact scenarios. (**A–D**) Across domains of language of the TLI and GBI dataset; (**E–H**) across phonological features (TLI dataset only, the GBI dataset does not include phonological data). (**A,E**) Genetic contact, all pairs; (**B,F**) genetic contact: different area pairs only; (**C,G**) genetic contact: same area pairs only; (**D,H**) co-location in the same area. Intervals show the 50% and 89% credible intervals of the combined effects per domain. Heatmaps show the median posterior difference between domains, with gray shading for differences whose 89% highest posterior density interval includes zero. For example, lexical class features in the GBI dataset are 3% less likely to be shared under genetic contact than features of lexical semantics (**A**).

For other domains our findings are even less clearly in line with previous research. LEXICAL SEMANTICS shows borrowing probabilities in a similar or higher range than grammar (LINEAR ORDER and OTHER GRAMMAR) and PHONOLOGY. There is one exception to this trend in the same-area condition in the GBI dataset (Fig. 4D) where LEXICAL SEMANTICS shows lower borrowing probability than LINEAR ORDER. These observations seem to partially confirm the notion that new lexical concepts remain easier to learn after childhood than other aspects of language, perhaps similar to linear order. At the same time, we note that GRAMMATICAL CATEGORIES show borrowing probabilities in a similar range (grey-shaded cells, Fig. 4A, C–D) as LEXICAL SEMANTICS or even higher (Fig. 4B) and, like LEXICAL SEMANTICS, they outrank LINEAR ORDER, OTHER GRAMMAR and PHONOLOGY, with median posterior differences of 1–3%, again with an exception in the same-area condition of the GBI dataset and also in the within-area genetic condition of the GBI dataset (Fig. 4C–D). Furthermore, LEXICAL CLASSES show higher or similar borrowing probabilities than lexical semantics in the TLI but not in the GBI dataset.

These observations are unexpected given that GRAMMATICAL CATEGORIES are difficult to acquire after the critical period of early childhood (*75*, *76*, *37*, *77*, *38*, *78*, *79*), i.e., at an age at which borrowing is mostly likely to be initiated. Several explanations of this mismatch are possible. One possibility is that adult learning of categories (e.g., the existence of a future tense) as opposed to their formal marking (e.g., by prefixes vs suffixes) is easier than suspected, or that this is specifically the case for the categories coded in our data and less so for those that have been commonly studied in (predominantly) European languages. Another possibility is that our categorization of features might not pick up the distinctions that matter for learnability. For example, it is possible that the opposite effects in LEXICAL CLASSES in the GBI and TLI datasets stem from differences in parts of speech, since the GBI dataset focuses more on nominal than verbal classes (table S23). Yet another possibility is that children might be more actively involved in borrowing than expected. This might be particularly important in complex and intensive contact situations such as the genetic contact between areas (Fig. 4B), a condition where our data mostly rests on contact under European colonialism and where the probability of borrowing grammatical categories is decisively higher than borrowing lexical semantics, at least in the GBI dataset.

While we cannot presently quantify the relative contributions of these possibilities to an overall explanation of our findings, the social conditions of colonial contact might be specifically relevant for capturing effects we find in features of PROSODY (Fig. 4F–G). Prosody is a particularly important marker of social belonging and differentiation (*80*, *81*). We find higher borrowing probabilities for prosody than for other aspects of phonology in genetic contact between areas (Fig. 4F). These are conditions with strong power imbalances associated with colonialism, exerting high pressure for assimilation due to prestige hierarchies (*82*). By contrast, features of prosody show lower borrowing probabilities than other aspects of phonology in genetic contact within areas. In fact, in this contact condition, prosodic features are even less likely to be shared than in the baseline condition, indicating divergence (Fig. 4G). This might be due to the function of prosodic features as markers of divergence in socially more balanced contact situations.

Overall, our findings suggest that, in contrast to received scholarship, models of globally fixed hierarchies of borrowing mostly fail to capture the outcomes of language contact. While some of the differences we find in borrowing probability (linear order vs. other grammar, consonants vs. vowels, lexical semantics vs. other features) partially match expectations, the evidence is not robust across conditions and datasets. Moreover, beyond these domains, we find glaring differences in the linguistic features that tend to be borrowed in contact. Critical drivers of these differences are demographic, political and sociocultural imbalances (*82*, *3*, *83*, *84*, *56*, *46*). Uneven contact situations, such as those resulting from colonialism, are usually associated with overall higher linguistic convergence between populations. More balanced contact events also usually lead to borrowing, but can also foster differentiation in language domains particularly common in identity signaling, such as prosody. These results paint a complex picture for the past and future of the world’s languages. Hierarchical social contact has likely impacted distributions of linguistic features and will continue to do so in an increasingly globalized and capitalized world confronted with demographic displacements induced by global warming.

## Acknowledgments

We thank Steven Moran for valuable discussion and ideas in the early phase of the project. We are also grateful to John Mansfield for discussions regarding schismogenesis. BP is grateful to the ASLAN project within the program “Investments for the Future”, French National Research Agency (ANR), ANR-10-LABX-0081.

## Author contributions

Conceptualization: AG, DEB, BP, CB, BB. Data curation: AG, CB, VB, BP. Methodology: AG, DEB, ER, BP, CB, BB. Investigation: AG, DEB, ER, CB, BB. Visualization: AG, CB, BB. Validation: AG, ER. Funding acquisition: KS, BB. Project administration: AG, CB. Supervision: KS, CB, BB. Writing – original draft: AG. Writing – review & editing: AG, DEB, ER, VB, DB, KS, BP, CB, BB.

## Data and materials availability

The results of the best ADMIXTURE runs per K, the linguistic data and metadata on both populations and languages are available on OSF: https://osf.io/29bam. The repository also hosts all analytical outputs, tables and figures and all code necessary to produce them.

