## Supplementary Materials for "Global patterns of genetic admixture reveal effects of language contact"

Materials and Methods

Materials

The genetic analysis relies on genomic data from an expanded version of the GeLaTo dataset (*85*, *49*). The database includes 4768 unrelated individuals in 558 genetic populations (median: 8 individuals) representing 373 languages. Each genetic population is assigned to a language with a Glottocode (*66*), following curating criteria based on anthropological and ethno-linguistic information available from the original genetic publication (*49*). These populations are listed in table S1. The literature review for additional admixture cases identified 45 additional instances of contact compatible with the criteria used through our ADMIXTURE run on genomic data (*86*–*92*, *61*, *93*–*99*). The structural linguistic data used in this research draws from the full “statistical” curations of the GBI (“Grambank Independent”) and TLI (“Typology Linked and Independent) datasets (*63*). These datasets were curated to reduce logical and strong universal statistical dependencies between features and are therefore suited for models like ours that assume conditional independence of the response. For area coding we used the 10 continent-sized areas (Fig. 1C) from AUTOTYP (*64*) in the main analysis, and the 6 macroareas (fig. S1B) from Glottolog (*66*) in the sensitivity analysis.

Methods

For the genetic analysis, we ran the ADMIXTURE software (*57*) assuming *K*=2 to *K*=30 ancestries with 20 runs per each *K*, retaining the run with the highest likelihood per *K*. The smallest cross-validation errors are found at K=23 and K=22. After checking the likelihood profiles and visually inspecting the consistency of the runs, we selected those from *K*=12 to *K*=30, since the lowest *K* (*K*=12) returned recognizable blocks of ancestry (fig. S1A). Previous studies using ADMIXTURE with global datasets but fewer population samples recognized solid blocks of ancestry at K=5 (*100*), K=8 (*101*) and K=10 (*89*), but these studies did not explore higher values of K. We then identified populations as admixed if their two largest ancestry components amounted to at least 70% of genetic ancestry, of which the smaller source contributed at least 5%. Minor sources contributing less than 5% of ancestry were considered so small that they would reflect noise. We furthermore required that these thresholds hold through at least 5 different levels of *K*: if they were admixed for fewer than 5 levels of *K*, we disregarded the admixture signal for the population as insufficiently consistent.

To identify the two source populations for the ancestry components in the admixed populations, we approximated these source populations for each *K* by populations available in the GeLaTo sample. We did this by flagging all populations that incorporate the admixture component in question at ≥ 80%. For each admixed (target) population, we therefore had a set of possible source 1 populations denoting the larger admixture component and a set of possible source 2 populations denoting the smaller admixture component. Source populations were ranked by *F_ST_*-value to the target population. *F_ST_* is a measure of pairwise genetic distance between populations, directly proportional to divergence time and inversely proportional to migration rate. Hence, lower *F*_ST_ values are shared between populations who are genetically related (*102*). Our final filtering step was to list all possible combinations of source 1 and source 2, where one of the two sources was represented by a population speaking a language of the same language family as the target population and the other represented a population speaking an unrelated language. This resulted in a list of genetic triplets consisting of a target, a source 1 and a source 2 population. For each triplet, we logged the following information: the number of distinct *K* over which the target population was admixed; for how many of these *K* the source 1 population appeared as a match for the larger admixture component; for how many of these *K* the source 2 population appeared as a match for the smaller admixture component; availability of structural linguistic data for the target population as well as each of the source populations. The target populations along with their unrelated source population were considered potential candidates for further analysis are retrievable from OSF (<https://osf.io/29bam/>, output/longlists/ longlist_for_manual_curation.csv) Of all 3506 candidates considered under the 70% admixture threshold, 81% also held under a more conservative threshold of 80% and 56% even held under a threshold of 90%.

This list of candidate triplets was then filtered to yield language pairs for which we had linguistic feature data available in GBI and TLI, with a (single) target language denoting the target population and a set of languages representing one or more languages in a clade unrelated to the target language associated with the source population. For some populations, we used closely related proxy languages in the linguistic database to increase data coverage. Table S1 lists all relevant populations’ language assignments in the original GeLaTo database and our mappings to GBI and TLI.

Many of the candidate triplets were spurious in that they only appeared inconsistently across a small number of *K*. For instance, Greek populations appeared admixed at the 70%-threshold for 5 different values of *K.* However, only one of these *K,* namely *K* = 12, marked the Greek as admixed with a non-Indo-European source (in this case: Afro-Asiatic). This signal was too weak to be included for analysis. Similarly, many candidates were dubious given external knowledge of population genetics in the relevant world region. For instance, the ancestry we know to be Indo-European that acted as a source for many admixture events in South America was often also additionally labelled as a Uralic ancestry. In these cases, we did not include contact pairs involving Uralic languages given our historical knowledge of imperialism and the colonization of the Americas. Lastly, in some cases, we were not able to assign languages to the admixture sources with any reasonable degree of confidence, or there was no reasonable linguistic data available for the languages. Overall, our curation yielded a set of 81 genetic contact pairs, to which we added 45 further pairs based on a literature review. Each pair set was assigned a unique pair ID, which was nested within 39 broad pair IDs that group together pairs with the same admixture source – for example, a single broad pair ID (broad pair ID 21) encompassed all seven instances of Spanish admixture into Central and South American populations (pair IDs 21.01, 21.02, 21.03, 21.04, 21.05, 21.06 and 21.07). The final set of pairs is available as table S2.

We then intersected the pairs with linguistic features from GBI and TLI. Specifically, for each dataset we assembled all available language pairs whose speaker populations are admixed. A single pair ID could thereby be represented by multiple language pairs, if several languages and/or dialects from the specified source clade were available for a given feature. In these cases, pair IDs and broad pair IDs handled relatedness between these potential source languages. For the GBI features, this yielded 348 language pairs; for the TLI features, it yielded 818 language pairs.

For every feature matched to at least one pair ID, we extracted the data for all pair sets. We recorded each target language’s and each source language’s state for the feature and recorded whether the pair has the same state or not. We additionally recorded whether the language pair is from the same continent-sized AUTOTYP area or (for the sensitivity analysis) Glottolog macro-area. To these language pairs, we added 300 randomly drawn language pairs for every feature to act as a baseline. For every feature with more than two states, we drew pairs for every state separately. This is because what carries potential for contact effects in multi-state features are feature *states* (e.g., a specific word order such as subject-object-verb) and not the overall feature (e.g., the word order of subject, object and verb).

Subsets of the resulting language pair data then served as input for a series of 5 Bayesian multi-level logistic regressions (m1–m5), once for the main analysis and once for the sensitivity analysis. We implemented the models (as defined in the Supplementary Text) in R’s brms (*103*) interface to Stan (*104*). Because the features from these datasets are only curated for reduced statistically dependence within but not across datasets, we fitted every model on the GBI and TLI data separately.

All models allow the global intercept (fixed effect) to vary by feature state (random effect). Given our state-wise (binarized) approach to modelling the features, the global intercept is at least 0.5 on the probability scale, motivating a prior with a positive mean for it. The models including information on areal co-location additionally include a global slope for the effect of areal contact that we allow to vary again by feature state, and also varying intercepts and slopes for each area combination, accounting for varying degrees of baseline similarity and of different contact effects among pairs, depending on the areas they are from. Models including genetic information included global slopes for the effect of genetic contact, that were allowed to vary by feature state as well as by each instance of contact labelled under the same pair ID. These latter slopes were nested within their broad pair ID. Together, these varying slopes account for structuring among different contact pairs. In the models including information on both areal and genetic contact, we also included varying slopes for genetic contact by area combination of the languages involved.

We ran prior and posterior predictive checks for every fitted model (figs. S12–S29). We performed model comparison with leave-one-out cross-validation using Pareto-smoothed importance sampling from the posterior, using the loo package (*105*).

To report state sharing, we drew posterior samples from the fitted statistical models from m1, m4 and m5 and computed probability differences for all feature states under genetic contact (in m1, m4 and m5) compared to the baseline or under the same area condition (in m1) compared to the different area condition. Using the mean and standard deviation of each difference, we then performed meta-analyses (see Supplementary Text for model definitions) with varying slopes by feature state group, using a categorical predictor to subsume feature states of the same group.

All data and scripts necessary to reproduce all elements of the analysis are available on OSF (<https://osf.io/29bam/>).

Supplementary Text

Formal definition of the statistical models

*Combined model, with both genetic and information on areal co-location (m1)*

$${same\_state}_{i} \sim Bernoulli\left( \pi_{i} \right)$$

$\pi_{i}= logit^{-1}(\eta_{i}$)

$$\eta_{i}=\alpha+\alpha_{STATE[i]}+\alpha_{AA[i]}$$

$$+{(\beta}_{1}+\beta_{A,STATE[i]})\times A_{i}$$

$$+(\beta_{2}+\beta_{G,STATE\left[ i \right]}+\beta_{\mathrm{AA}\left[ i \right]}+\beta_{BROAD\_ID/PAIR\_ID\left[ i \right]})\times G_{i}$$

$$\binom{\alpha_{\mathrm{AA}}}{\beta_{\mathrm{AA}}} \sim mvN\left( \binom{0}{0},\Sigma_{\mathrm{AA}} \right)$$

$$\Sigma_{\mathrm{AA}}=\left( \begin{matrix} \sigma_{\alpha_{\mathrm{AA}}}^{2} & \sigma_{\alpha_{\mathrm{AA}}}\sigma_{\beta_{\mathrm{AA}}}\rho\\ \sigma_{\alpha_{\mathrm{AA}}}\sigma_{\beta_{\mathrm{AA}}}\rho& \sigma_{\beta_{\mathrm{AA}}}^{2} \end{matrix} \right)$$

$$\left( \begin{matrix} \alpha_{\mathrm{STATE}} \\ \beta_{A,STATE} \\ \beta_{G,STATE} \end{matrix} \right) \sim mvN\left( \left( \begin{matrix} 0 \\ 0 \\ 0 \end{matrix} \right),\Sigma_{\mathrm{STATE}} \right)$$

$$\Sigma_{\mathrm{STATE}}=\left( \begin{matrix} \sigma_{\alpha_{\mathrm{STATE}}}^{2} & \sigma_{\alpha_{\mathrm{STATE}}}\sigma_{\beta_{A,STATE}}\rho& \sigma_{\alpha_{\mathrm{STATE}}}\sigma_{\beta_{G,STATE}}\rho\\ \sigma_{\alpha_{\mathrm{STATE}}}\sigma_{\beta_{A,STATE}}\rho& \sigma_{\beta_{A,STATE}}^{2} & \sigma_{\beta_{A,STATE}}\sigma_{\beta_{G,STATE}}\rho\\ \sigma_{\alpha_{\mathrm{STATE}}}\sigma_{\beta_{G,STATE}}\rho& \sigma_{\beta_{A,STATE}}\sigma_{\beta_{G,STATE}}\rho& \sigma_{\beta_{G,STATE}}^{2} \end{matrix} \right)$$

$\alpha$ indicates the intercept, β_1_ indicates the fixed effect of a language pair being from the same area (1) or not (0) (A). β_2_ indicates the fixed effect of a language pair representing a genetic contact pair (1) or a baseline pair (0) (G).

The terms $\alpha_{STATE[i]}$ and $\alpha_{AA[i]}$ indicate that the intercept can vary by state and area combination. $\beta_{A,STATE\left[ i \right]}$ and $\beta_{G,STATE\left[ i \right]}$ indicate that the both the area and the genetic contact slope can vary by state. $\beta_{AA[i]}$ and $\beta_{BROAD\_ID/PAIR\_ID\left[ i \right]}$ indicate that the genetic contact slope can further vary by area combination and by each contact instance (pair_id), nested within broad_id (grouping contact instances with the same source).

We set the following priors on model parameters:

$$\alpha\sim N\left( 0.75, 0.5 \right)$$

$$\beta_{1,}\beta_{2,}\sim N\left( 0, 1.5 \right)$$

$$\alpha_{\mathrm{STATE}\left[ i \right]},\beta_{A,STATE\left[ i \right]},\beta_{G,STATE\left[ i \right]}\sim N\left( 0, \sigma_{STATE[i]} \right)$$

$$\alpha_{\mathrm{AA}\left[ i \right]}{,\beta}_{\mathrm{AA}\left[ i \right]} \sim N\left( 0, \sigma_{AA[i]} \right)$$

$$\beta_{BROAD\_ID/PAIR\_ID\left[ i \right]} \sim N\left( 0, \sigma_{BROAD\_ID/PAIR\_ID\left[ i \right]} \right)$$

$$\sigma_{STATE[i]}, \sigma_{AA[i],}\sigma_{BROAD\_ID/PAIR\_ID\left[ i \right]} \sim N\left( 0, 2 \right)$$

$$\mathbf{R}_{\mathbf{AA}},\mathbf{R}_{\mathbf{STATE}} \sim LKJ\left( 2 \right)$$

The intercept is assumed to follow a normal prior with mean 0.75 and standard deviation 0.5. Fixed effects are each assumed to follow a normal prior with mean 0 and standard deviation 1.5. Each varying effect individually follows a normal prior with mean 0 and standard deviation 2. Jointly, the varying effects by area combination follow multivariate normal priors with mean 0 and variance-covariance $\Sigma_{\mathrm{AA}}$. The effects by feature state follow multivariate normal priors with mean 0 and variance-covariance $\Sigma_{\mathrm{STATE}}$. The co-variance matrices are decomposed into a prior standard deviation vector and a correlation matrix **R**. These **R** matrices are individually drawn from an LKJ(2) prior.

This model (m1) was fitted using feature states from GBI and TLI separately.

*Model with areal information only (m2)*

$${same\_state}_{i} \sim Bernoulli\left( \pi_{i} \right)$$

$\pi_{i}= logit^{-1}(\eta_{i}$)

$$\eta_{i}=\alpha+\alpha_{\mathrm{STATE}\left[ i \right]}+\alpha_{\mathrm{AA}\left[ i \right]}+{(\beta}_{1}+\beta_{\mathrm{STATE}\left[ i \right]})\times A_{i}$$

$$\binom{\alpha_{\mathrm{STATE}}}{\beta_{\mathrm{STATE}}} \sim mvN\left( \binom{0}{0},\Sigma_{\mathrm{STATE}} \right)$$

$$\Sigma_{\mathrm{STATE}}=\left( \begin{matrix} \sigma_{\alpha_{\mathrm{STATE}}}^{2} & \sigma_{\alpha_{\mathrm{STATE}}}\sigma_{\beta_{\mathrm{STATE}}}\rho\\ \sigma_{\alpha_{\mathrm{STATE}}}\sigma_{\beta_{\mathrm{STATE}}}\rho& \sigma_{\beta_{\mathrm{STATE}}}^{2} \end{matrix} \right)$$

The priors drawn are the same as in model m1.

$\alpha$ indicates the intercept, β_1_ indicates the fixed effect of a language pair being from the same area (1) or not (0) (A). The terms $\alpha_{STATE[i]}$ and $\alpha_{AA[i]}$ indicate that the intercept can vary by feature state and area combination. $\beta_{\mathrm{STATE}\left[ i \right]}$ indicates that the area slope can vary by feature state.

This model (m2) was fitted using feature states from GBI and TLI separately.

*Model with genetic information only (m3, m4, m5)*

$${same\_state}_{i} \sim Bernoulli\left( \pi_{i} \right)$$

$\pi_{i}= logit^{-1}(\eta_{i}$)

$$\eta_{i}=\alpha+\alpha_{\mathrm{STATE}\left[ i \right]}+{(\beta}_{1}+\beta_{\mathrm{STATE}\left[ i \right]}+\beta_{BROAD\_ID/PAIR\_ID\left[ i \right]})\times G_{i}$$

$$\binom{\alpha_{\mathrm{STATE}}}{\beta_{\mathrm{STATE}}} \sim mvN\left( \binom{0}{0},\Sigma_{\mathrm{STATE}} \right)$$

$$\Sigma_{\mathrm{STATE}}=\left( \begin{matrix} \sigma_{\alpha_{\mathrm{STATE}}}^{2} & \sigma_{\alpha_{\mathrm{STATE}}}\sigma_{\beta_{\mathrm{STATE}}}\rho\\ \sigma_{\alpha_{\mathrm{STATE}}}\sigma_{\beta_{\mathrm{STATE}}}\rho& \sigma_{\beta_{\mathrm{STATE}}}^{2} \end{matrix} \right)$$

The priors drawn are the same as in model m1.

$\alpha$ indicates the intercept, β_1_ indicates the fixed effect of a language pair representing a genetic contact pair (1) or a baseline pair (0) (G). The term $\alpha_{STATE[i]}$ indicates that the intercept can vary by feature state. $\beta_{\mathrm{STATE}\left[ i \right]}$ indicates that the genetic contact slope can vary by feature state. The random slope $\beta_{BROAD\_ID/PAIR\_ID\left[ i \right]}$ allows for contact effects to vary for each contact instance (pair_id), nested within broad_id (grouping contact instances with the same source).

This model was fitted once on available pairs (m3), once on only pairs of languages from the same area (m4) and once on only pairs from different areas (m5). In each case, separate models were fitted for GBI and TLI feature states.

*Meta-analysis model*

$$y_{i} \sim N(\mu_{i},\sigma_{i})$$

$$\mu_{i}=\beta\times g_{i}$$

$$\beta\sim N(0, 1.5)$$

The response $y_{i}$ refers to the effect of contact for each feature state. $\beta$ indicates the coefficient for each level of *g*, where *g* is the categorical variable classifying states into domain groups (see Main text). We fitted separate models for two separate classifications g1 and g2 for each of four types of contact (genetic contact, all pairs; genetic contact, different area pairs only; genetic contact, same area pairs only; same area).

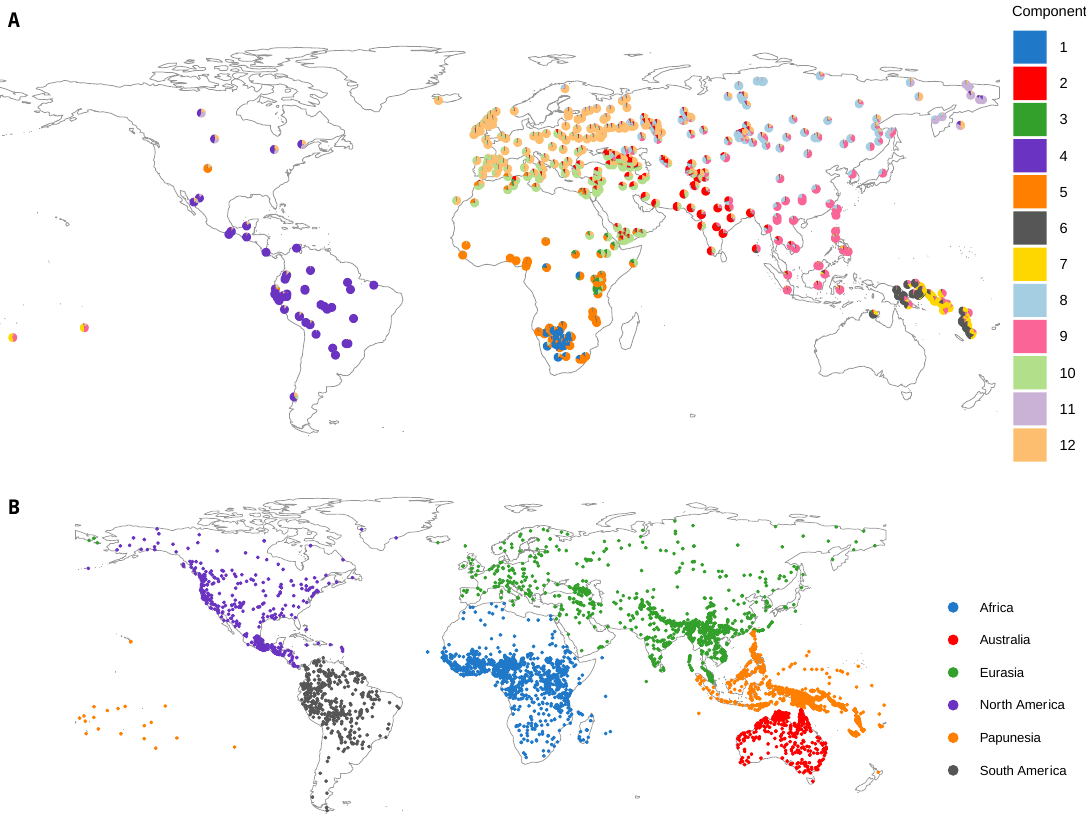

fig. S1.

(**A**). Admixture pie chart for GeLaTo populations, assuming *K =* 12 ancestry components. (**B**). Language assignments to the six “macroareas” from Glottolog.

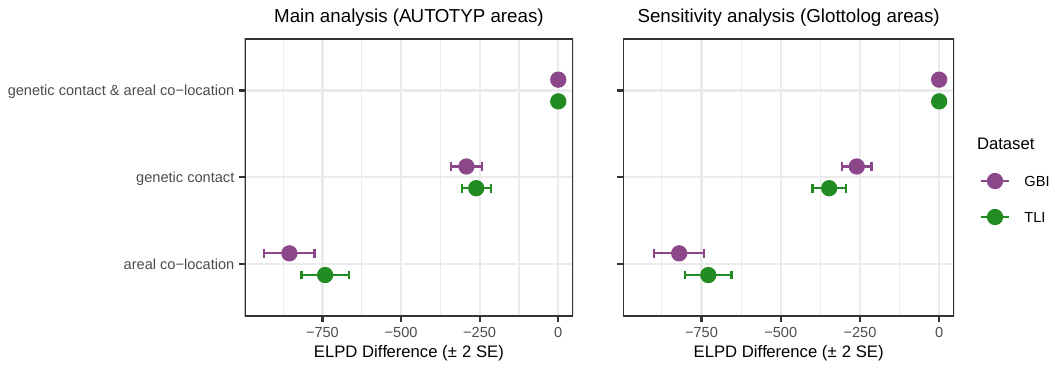

fig. S2.

Model comparison using ELPD differences between the models using as fixed effects: both genetic contact and areal co-location (m1), only areal co-location (m2), only genetic contact (m3).

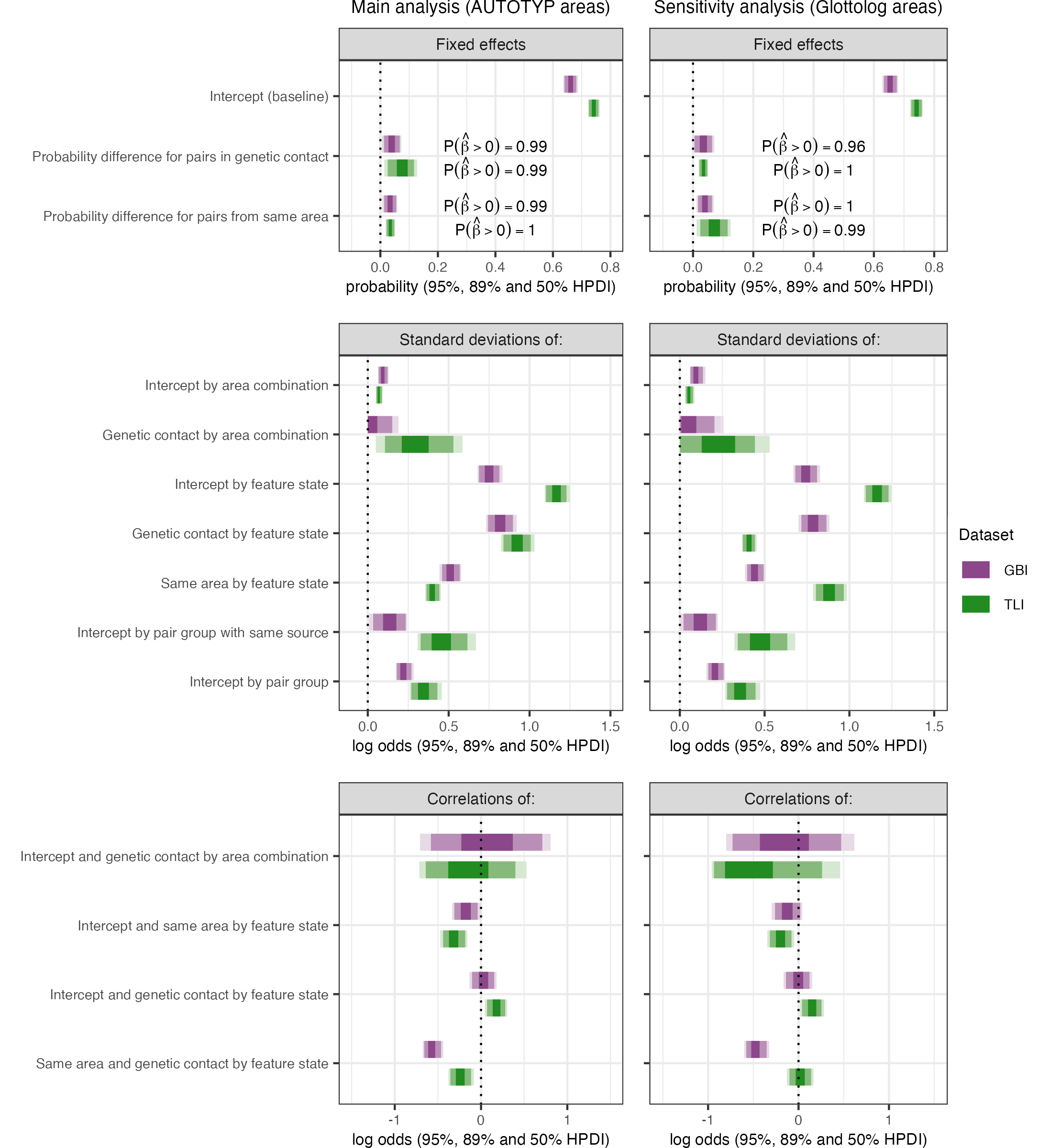

fig. S3.

95%-, 89%- and 50%-HPDIs for model estimates of the combined models (m1) from fig. S2.

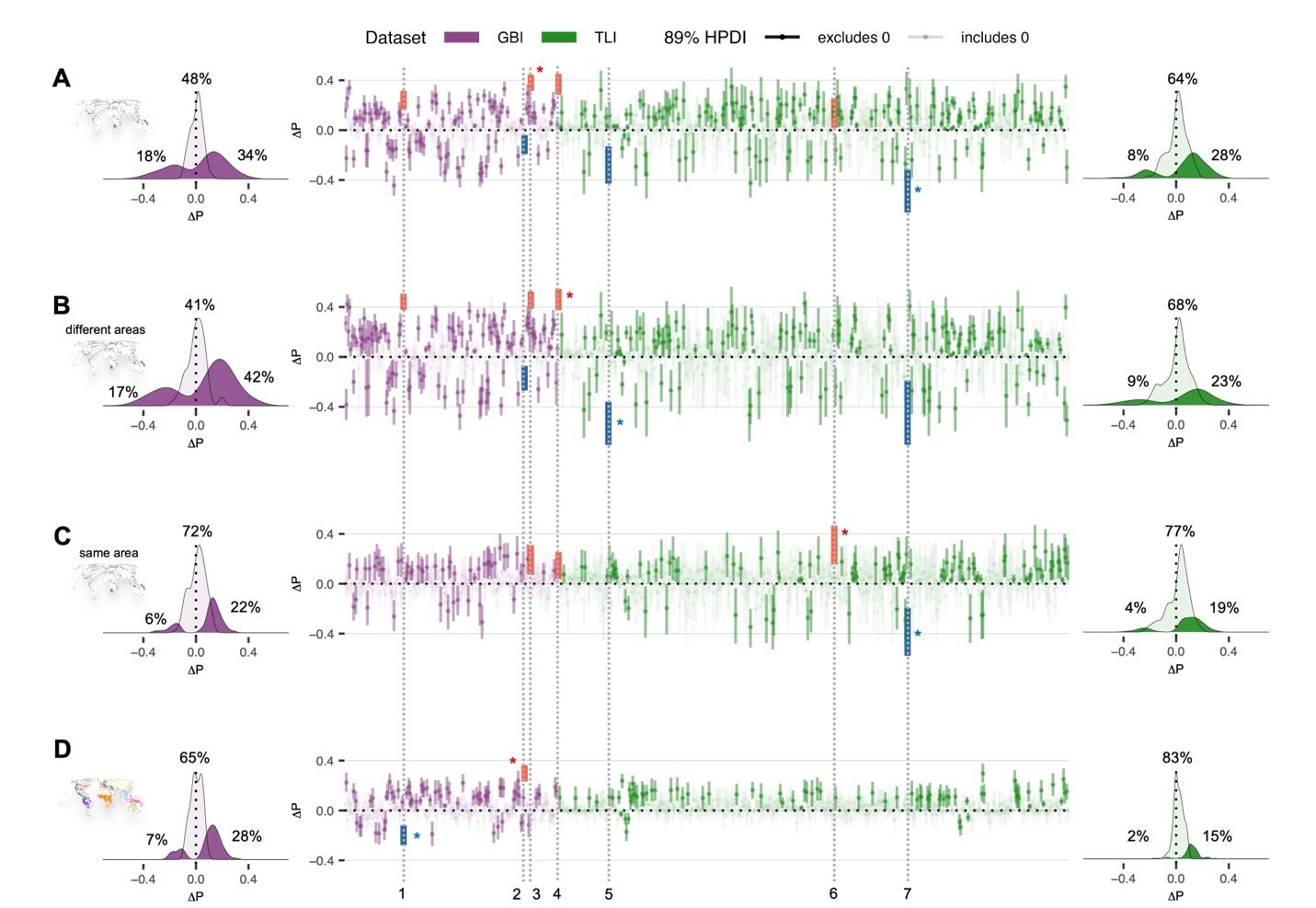
fig. S4.

Effects of each contact type on the probability of language pairs sharing structural states in each dataset, using the AUTOTYP areas: (**A**) genetic contact (all pairs); (**B**) genetic contact (different area pairs only); (**C**) genetic contact (same area pairs only). (**D**) areal contact. Effects including zero in the 89% HPDI are plotted with transparent shading. Density plots show the mean contact effect on sharing across all features states, with percentage values indicating the proportion of states less likely, equally likely and more likely to be shared under each contact type compared to the relevant baseline. Interval plots show the effect intervals per state. The x-axis shows all 683 states, ordered alphabetically. The y-axis shows the difference in probability of sharing between contact vs. baseline pairs. Stars indicate the state with the highest positive (red) and highest negative (blue) effect in each panel. These states are then highlighted across the four contact types (panels A-D) and labelled (1-7). Grey lines allow the comparison of the effect of each highlighted state across contact conditions: States are coloured if they have a significant (excluding zero) diverging effect (in blue) or borrowing effect (in red). 1: GB126, existential verb. 2: GB623drmc, verbal prefixes and/or proclitics for speech act participants. 3: GB704drm, diminutive and/or augmentative marking on noun. 4: GB995F, noun-adjective and noun-demonstrative order. 5: TLI0560, concept of ‘eye’ colexified in the word for ‘tear’. 6: TLI0959: presence of distributive numerals. 7: TLI1017, word for distributive numerals.

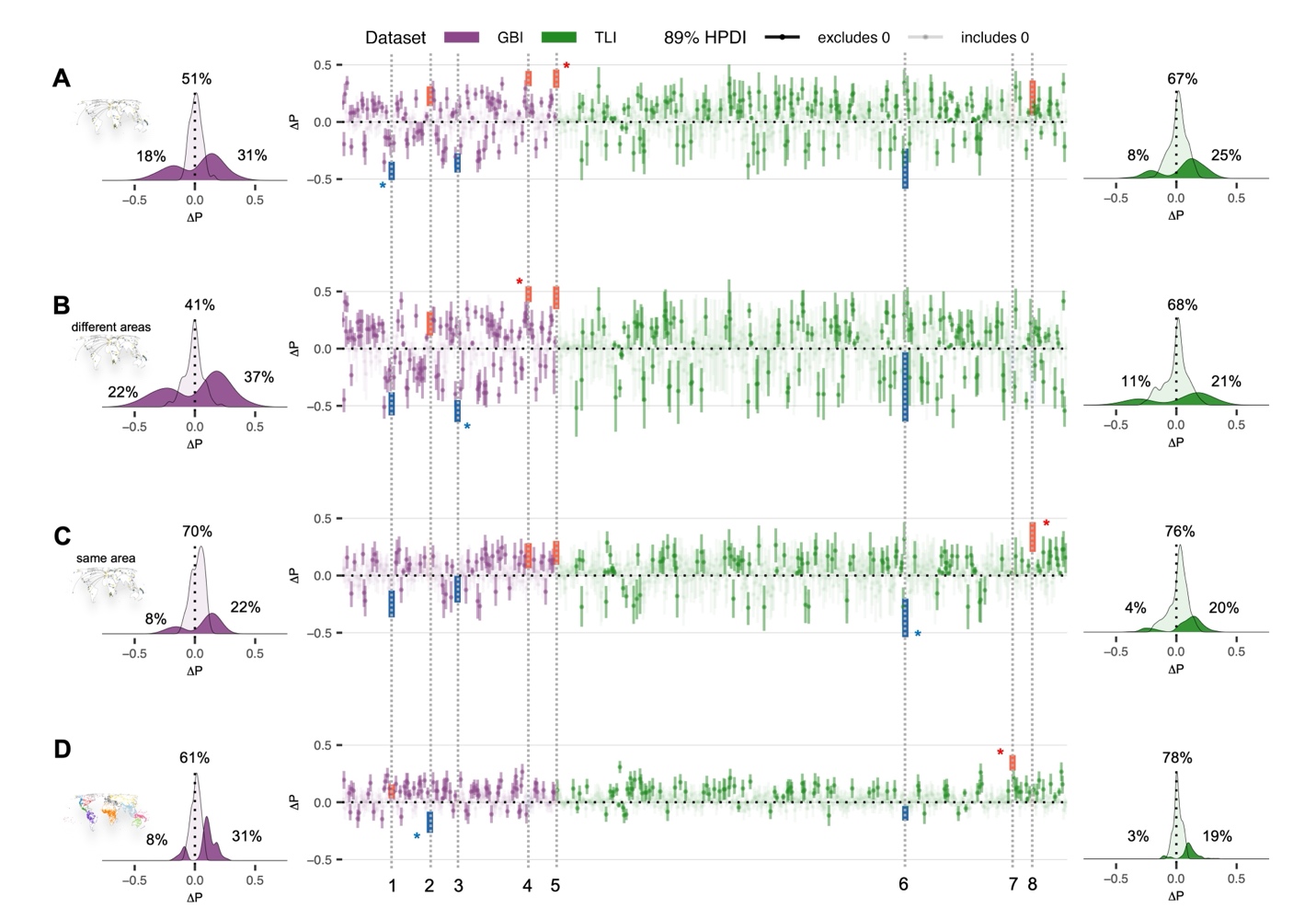
fig. S5.

Effects of each contact type on the probability of language pairs sharing structural states in each dataset, using Glottolog areas. Same plotting conventions as in fig. S4: GB110cC, verb suppletion for tense or aspect (if morphological tense/aspect present). 2: GB196C, male-female distinction in 2p pronoun (if gender in 3p pronoun). 3: GB302c, phonologically free passive marker (if passive present). 4: GB704drm, diminutive and/or augmentative marking on noun. 5: GB995F, word order noun-adjective and noun-demonstrative. 6: TLI1017, word for distributive numerals. 7: TLI1084, indefinite pronouns related to generic nouns. 8: TLI1094, valency marking on fully inflected verb.

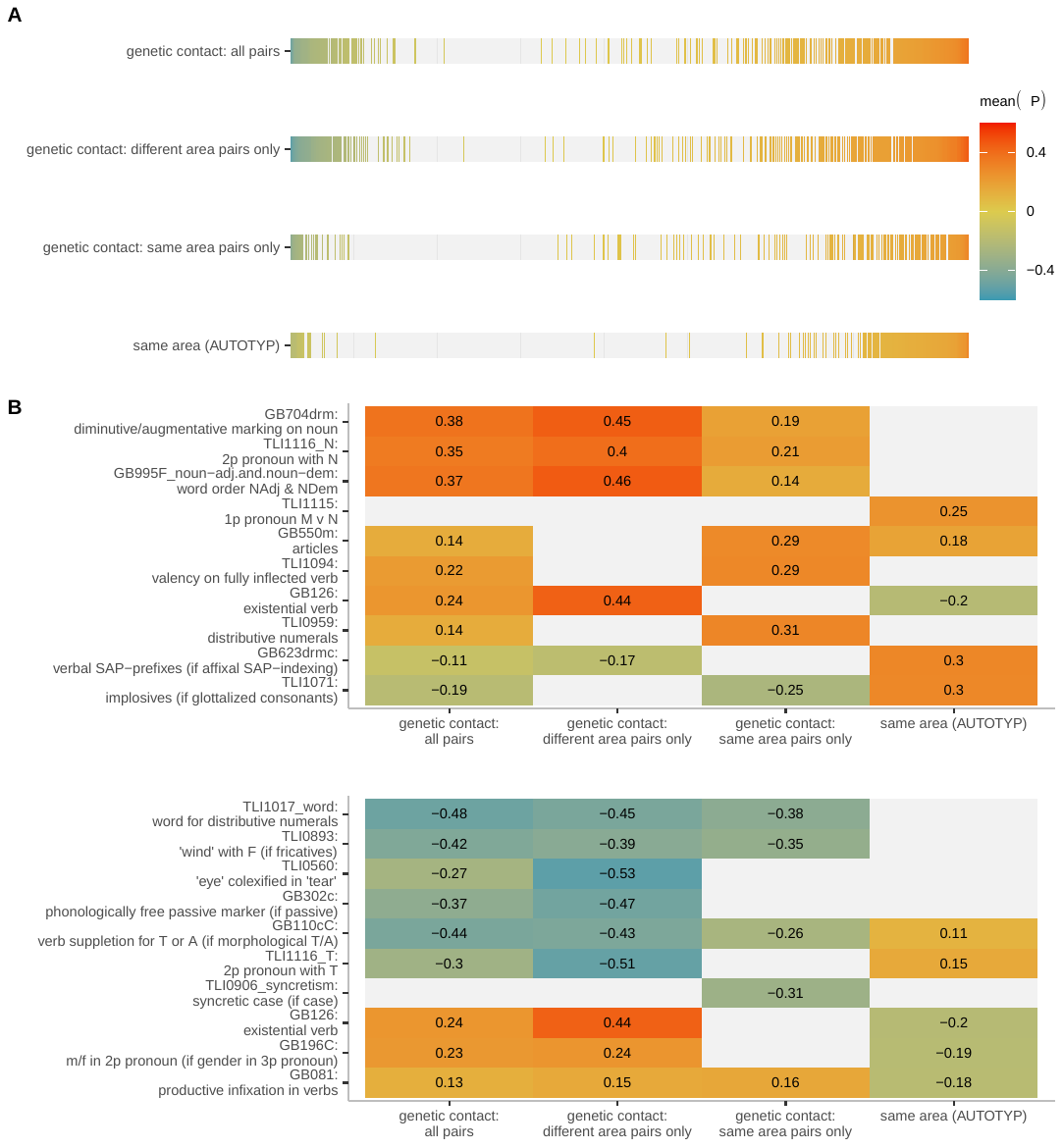
fig. S6.

Alternative visualization of state sharing effect scales under each contact condition using AUTOTYP areas (genetic contact with all pairs; with different area pairs only; with same area pairs only; areal contact). Color indicates state sharing differences: shades of orange and red indicate convergence; shades of green and blue indicate divergence under contact. 89%-HPDIs including 0 are colored grey: for these, we cannot confidently declare a contact effect in any direction. (**A**) Individual heatmap scales of state sharing effects under each condition, each sorted in ascending magnitude of mean posterior effect. (**B**) Heatmaps of the three feature states with the strongest positive and negative (upper v. lower panel) effect on state sharing under each contact condition.

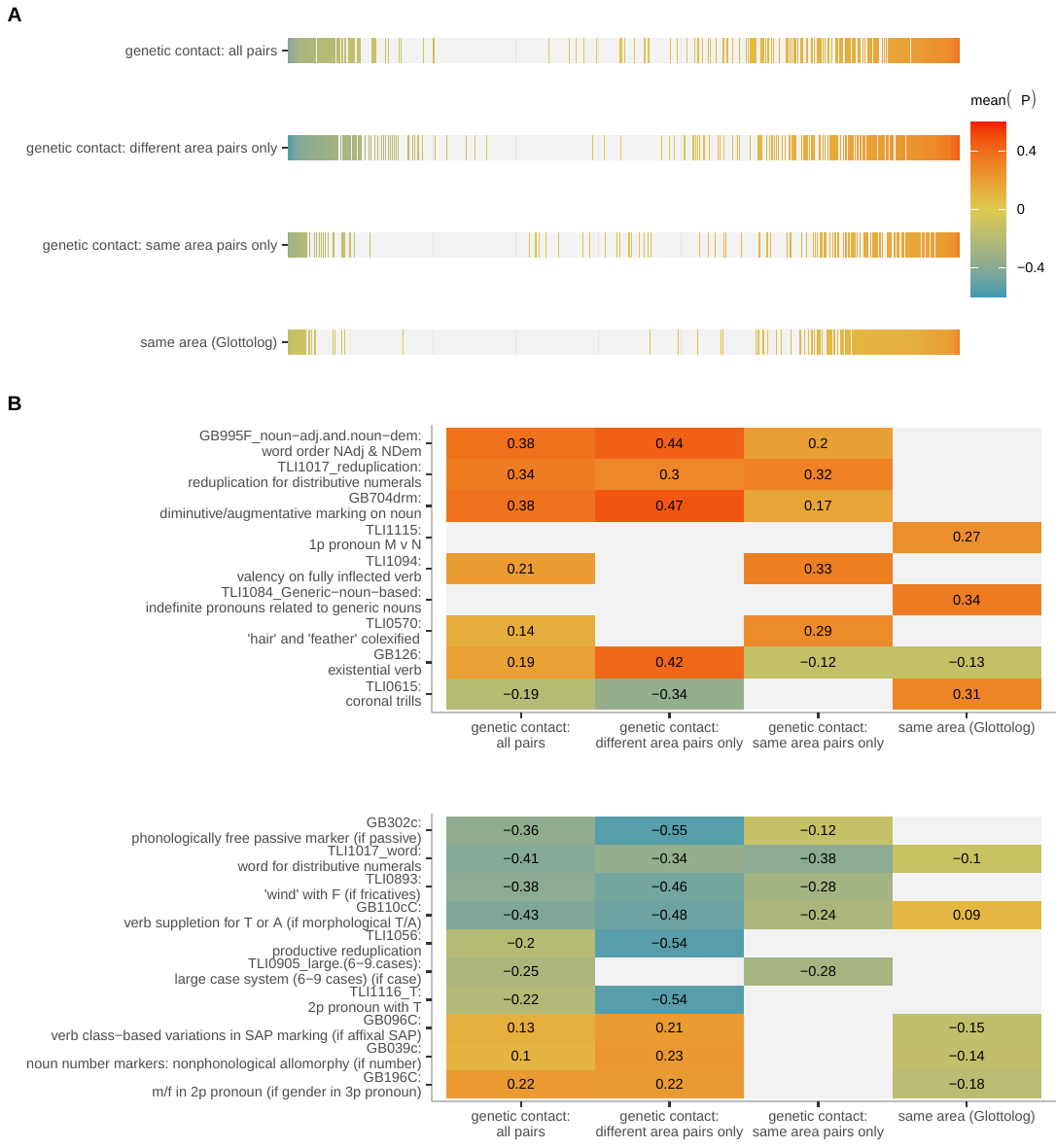
fig. S7.

Alternative visualization of patterns of state sharing effect scales under each contact condition in the sensitivity analysis using the Glottolog areas. Same plotting conventions as in fig. S6.

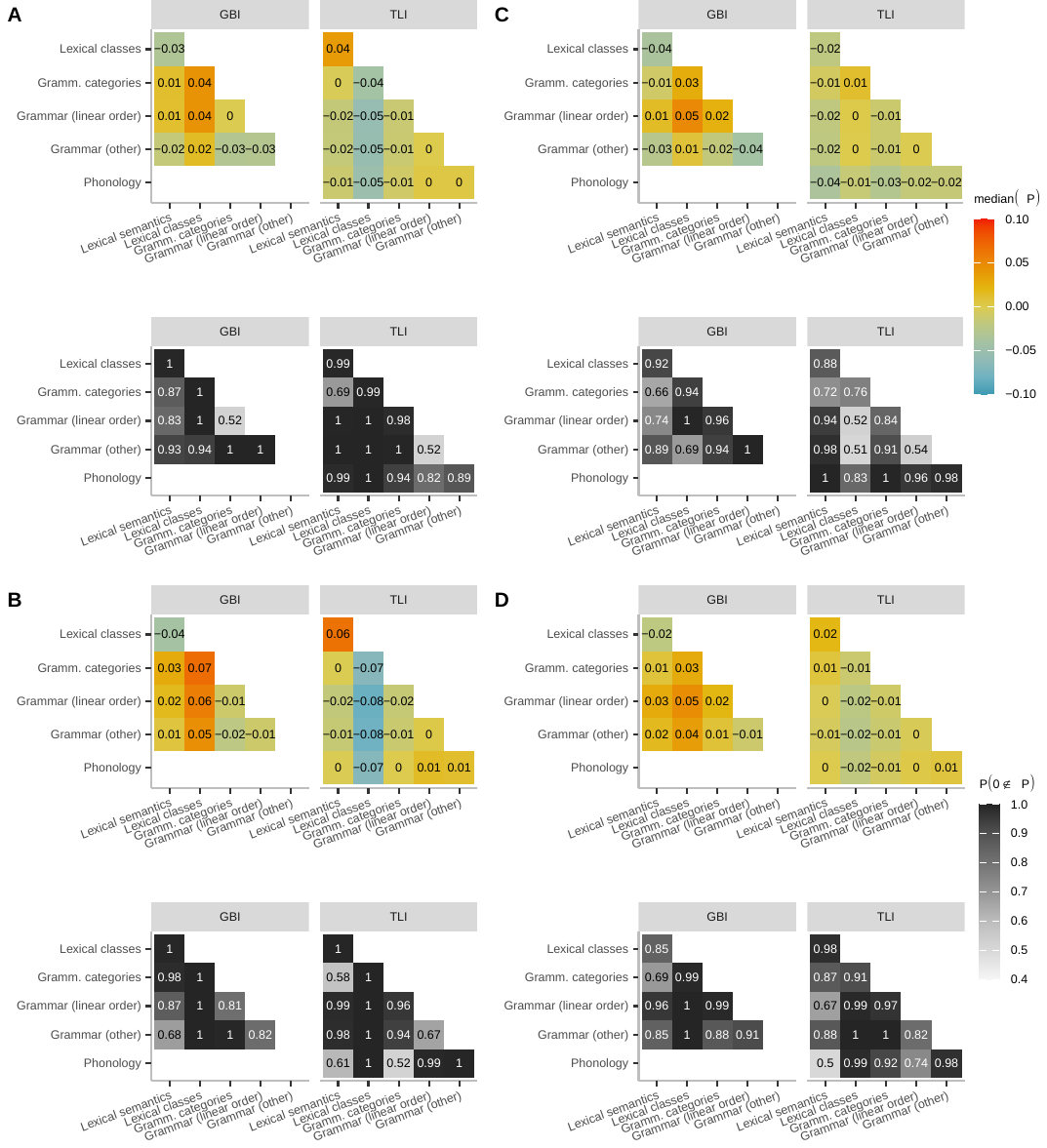
fig. S8.

Meta-analysis of contact effects on features across domains of language under different types of contact, as an extension of Fig 4. The heatmap (upper plot) of each panel shows the posterior medians of borrowing probability differences between pairs of domains, irrespectively of the posterior’s total distribution. The lower plot of each panel with a grey scale records the posterior probability of a non-zero difference. Contact types in subfigures are: (**A**) genetic contact, all pairs; (**B**) genetic contact: different area pairs only; (**C**) genetic contact: same area pairs only; (**D**) belonging to the same area (AUTOTYP). The GBI dataset does not include phonological data.

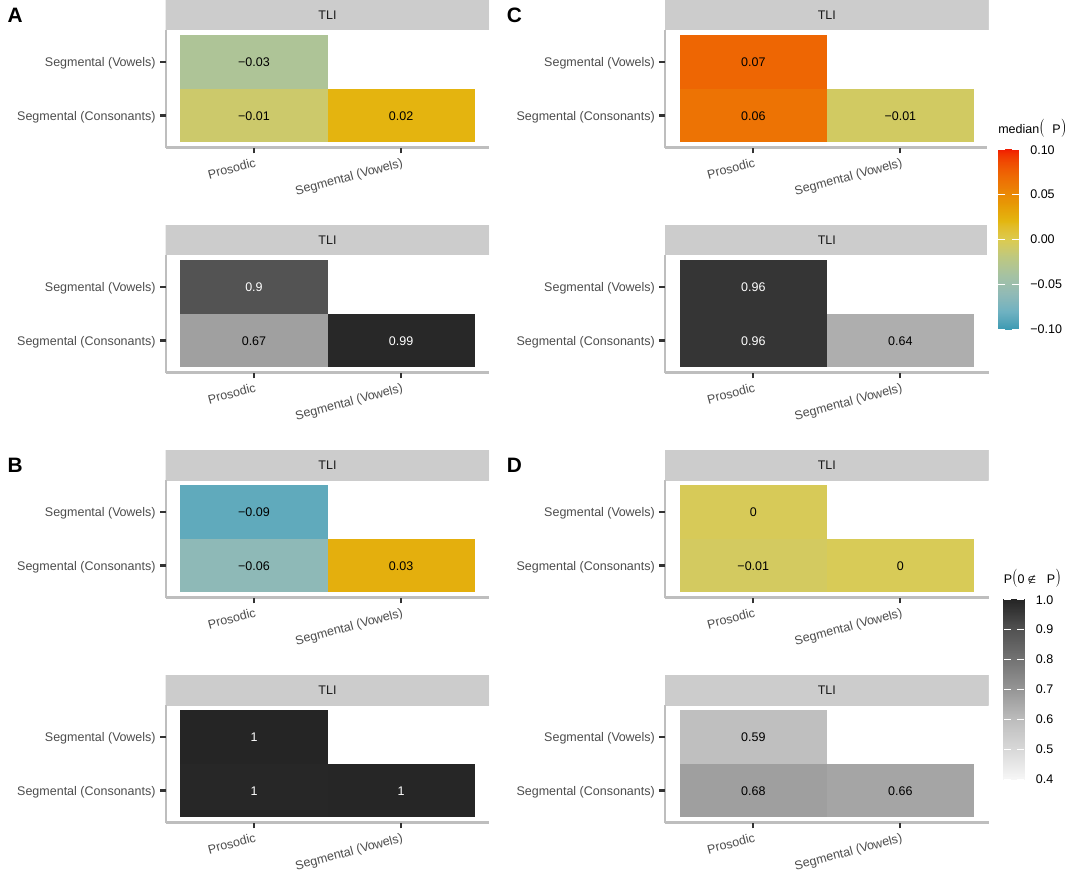
fig. S9.

Meta-analysis of contact effects in phonology under different types of contact, extension of Fig 4. Same plotting conventions as fig. S8.

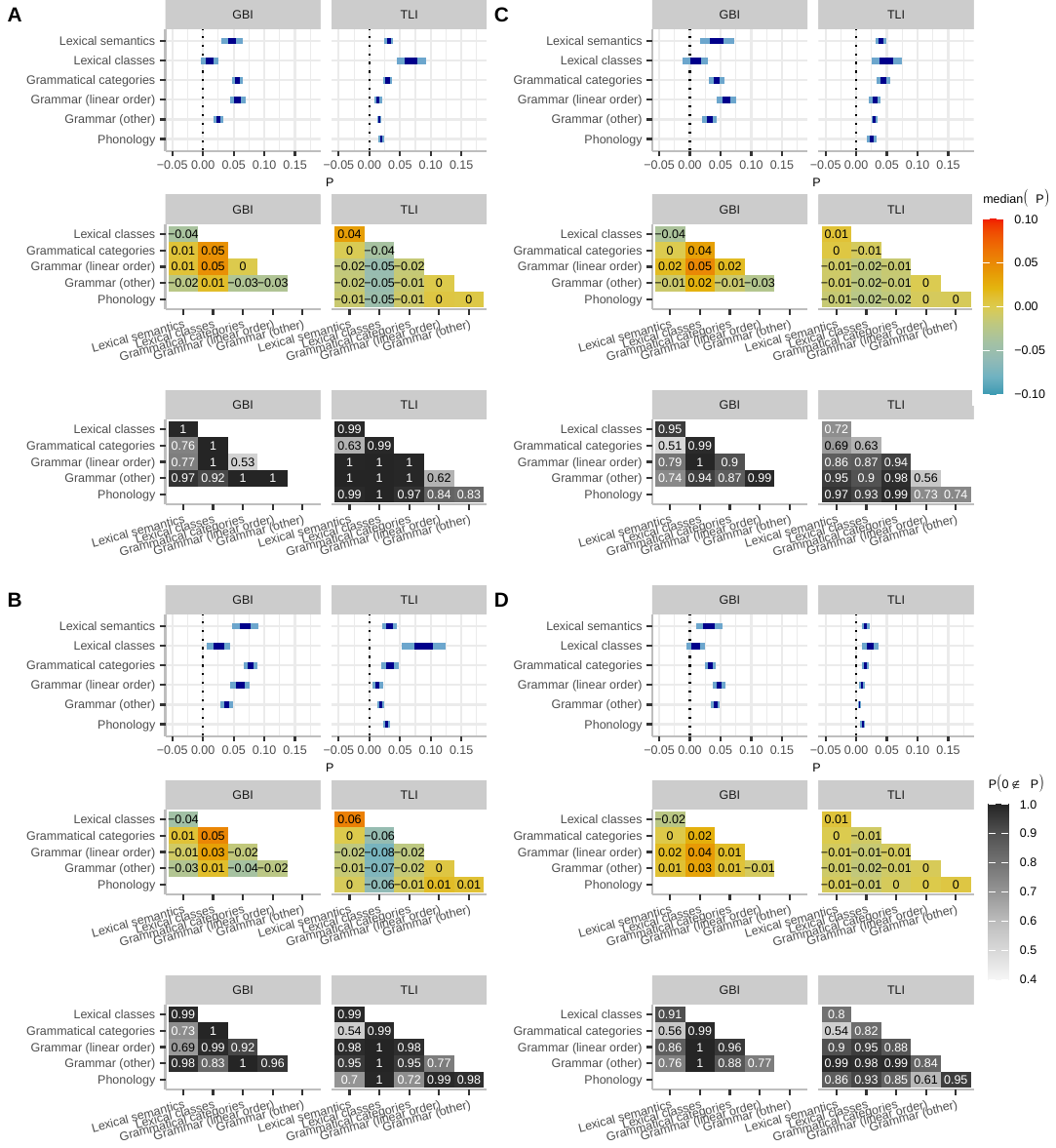
fig. S10.

Meta-analysis of contact effects on features across domains under different types of contact, using the six Glottolog areas (sensitivity analysis). Same plotting conventions as fig. S8.

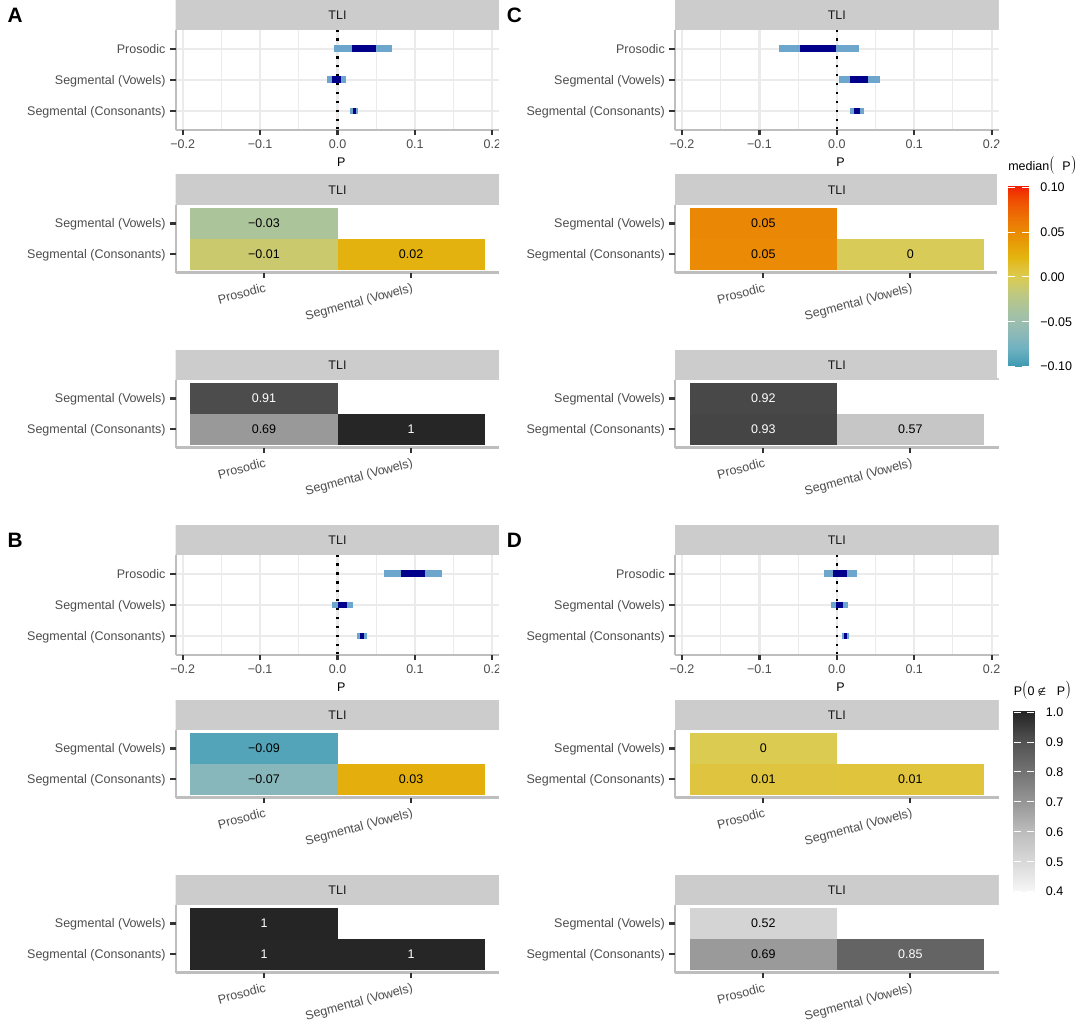

fig. S11.

Meta-analysis of contact effects in phonology, using the six Glottolog areas (sensitivity analysis). Same plotting conventions as fig. S8.

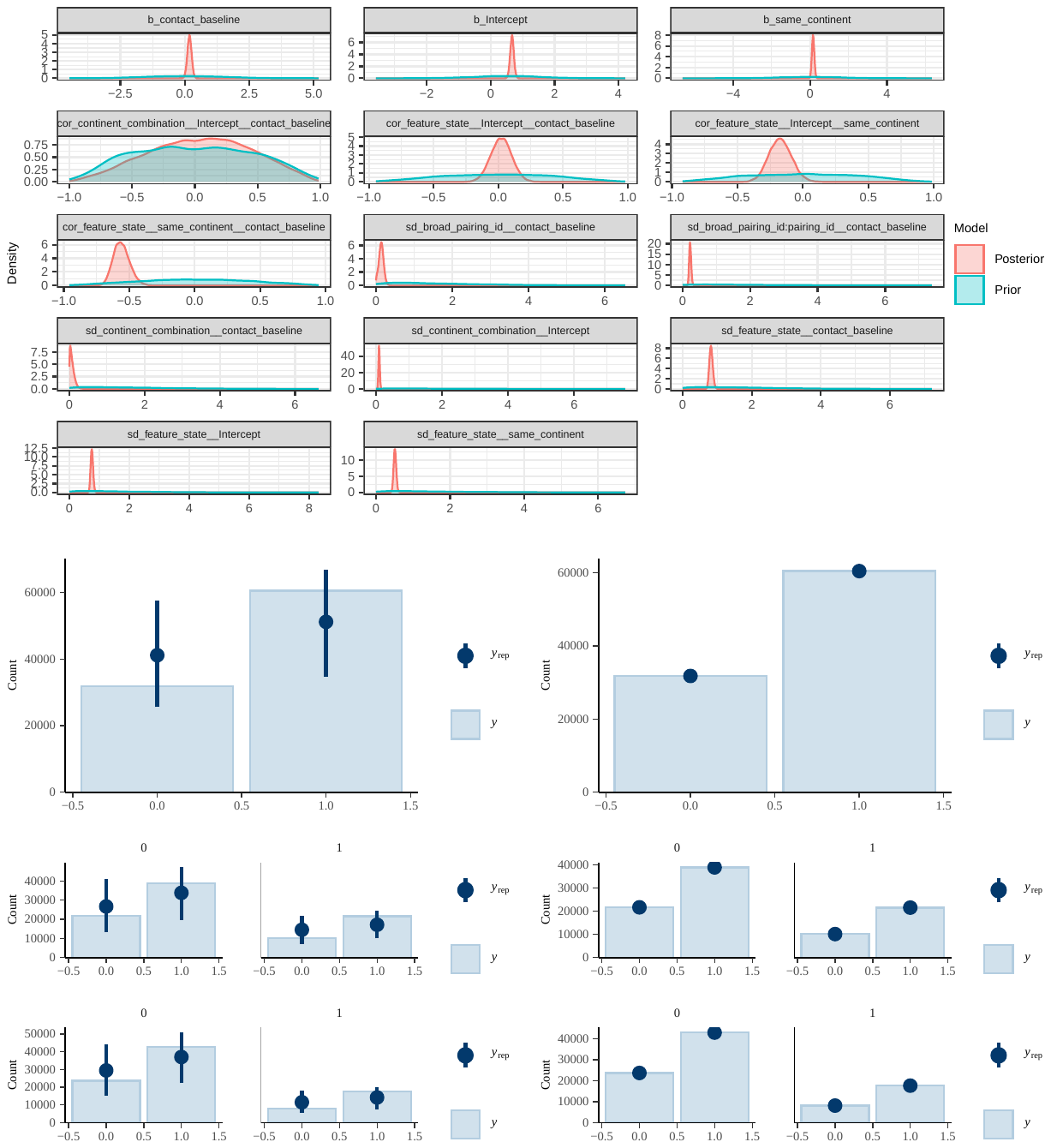

fig. S12.

Prior and posterior distributions for regression coefficients and multilevel hyperparameters (top) as well as prior predictive checks (bottom left) and posterior predictive checks (bottom right) for regression coefficients: combined model (genetic contact and information on areal co-location), including all pairs, GBI data, main analysis (AUTOTYP areas).

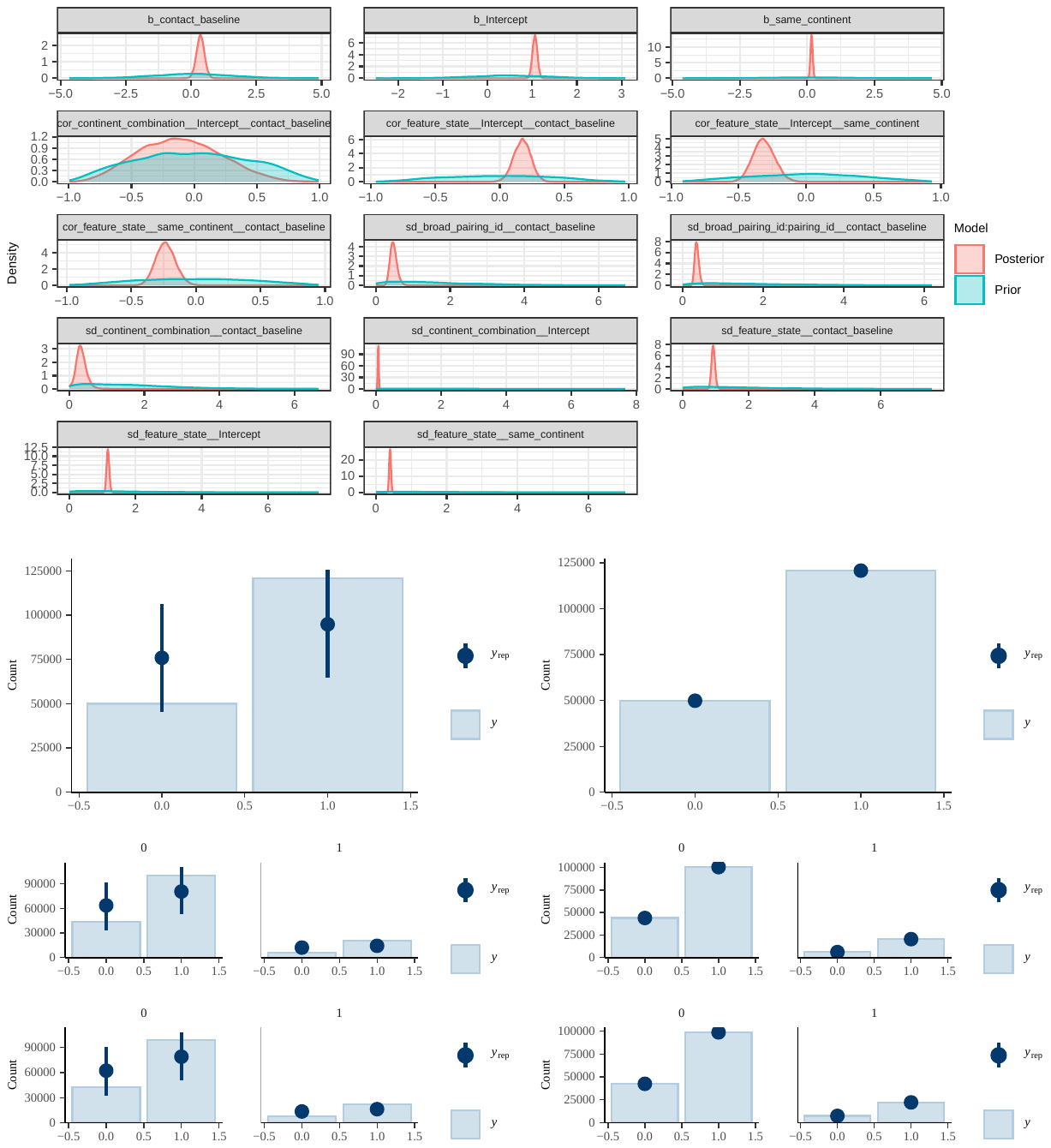

fig. S13.

Prior and posterior distributions for regression coefficients and multilevel hyperparameters (top) as well as prior predictive checks (bottom left) and posterior predictive checks (bottom right) for regression coefficients: combined model (genetic contact and information on areal co-location), including all pairs, TLI data, main analysis (AUTOTYP areas).

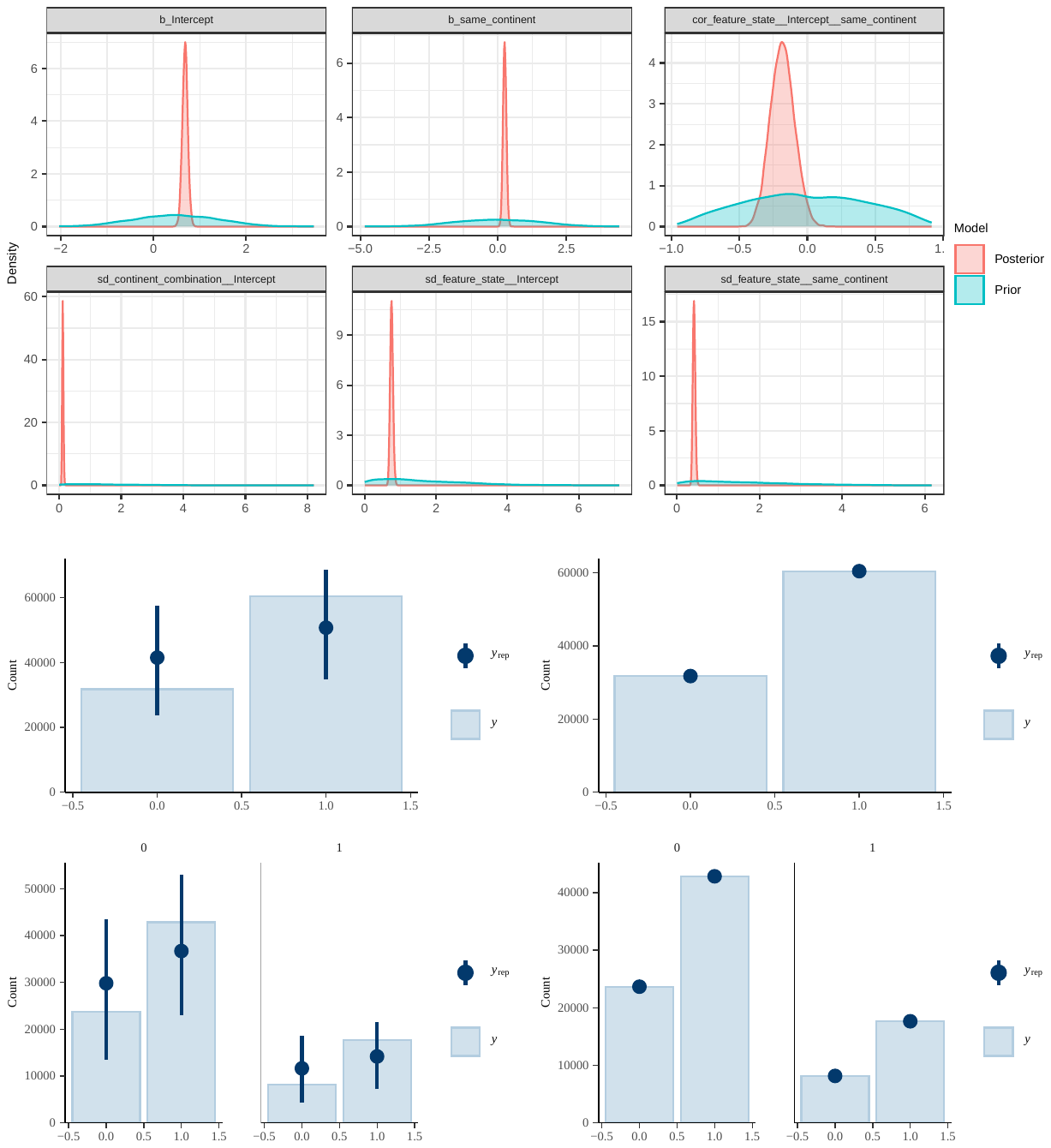

fig. S14.

Prior and posterior distributions for regression coefficients and multilevel hyperparameters (top) as well as prior predictive checks (bottom left) and posterior predictive checks (bottom right) for regression coefficients: areal model, including all pairs, GBI data, main analysis (AUTOTYP areas).

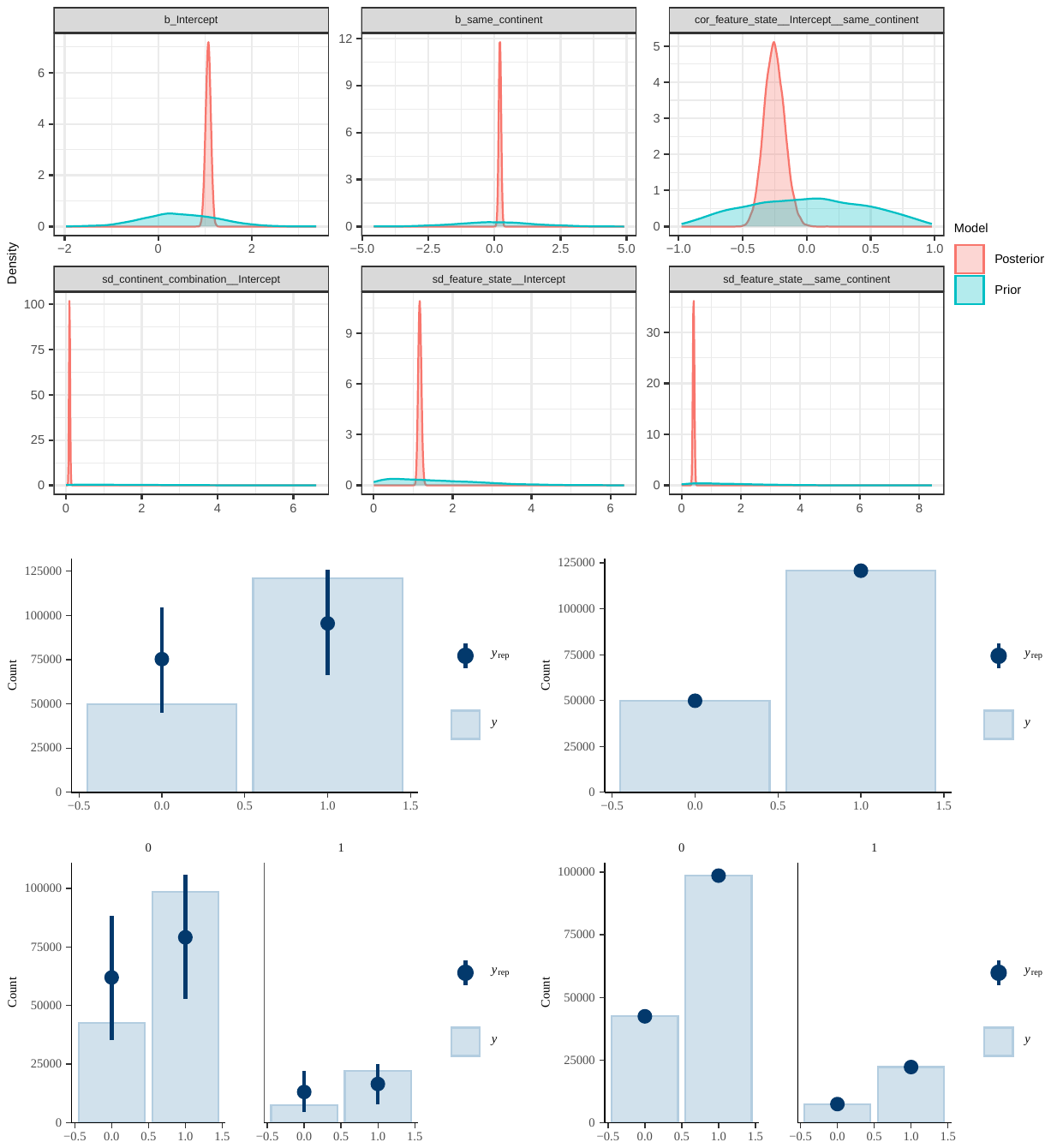

fig. S15.

Prior and posterior distributions for regression coefficients and multilevel hyperparameters (top) as well as prior predictive checks (bottom left) and posterior predictive checks (bottom right) for regression coefficients: areal model, including all pairs, TLI data, main analysis (AUTOTYP areas).

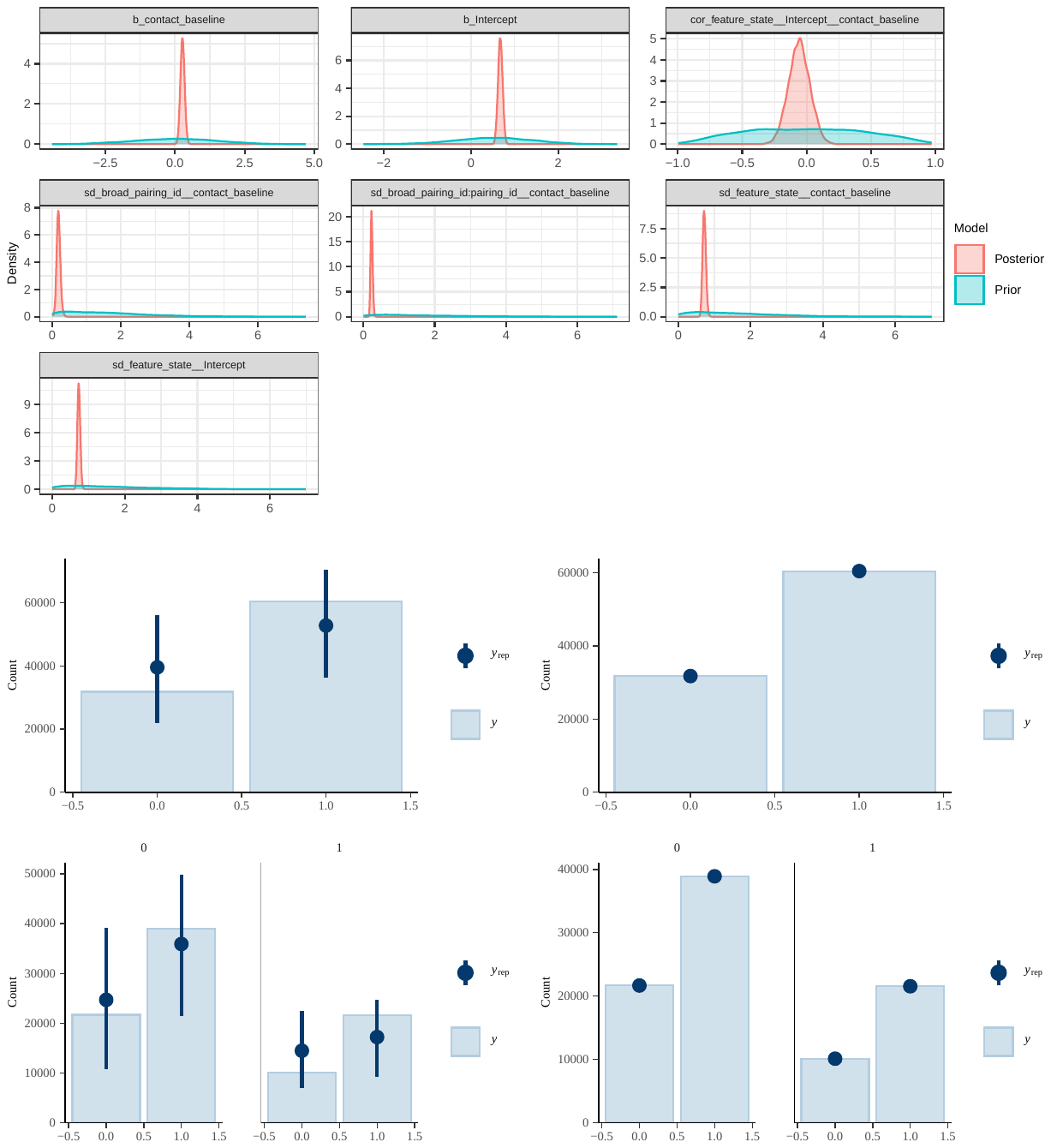
fig. S16.

Prior and posterior distributions for regression coefficients and multilevel hyperparameters (top) as well as prior predictive checks (bottom left) and posterior predictive checks (bottom right) for regression coefficients: genetic model, including all pairs, GBI data.

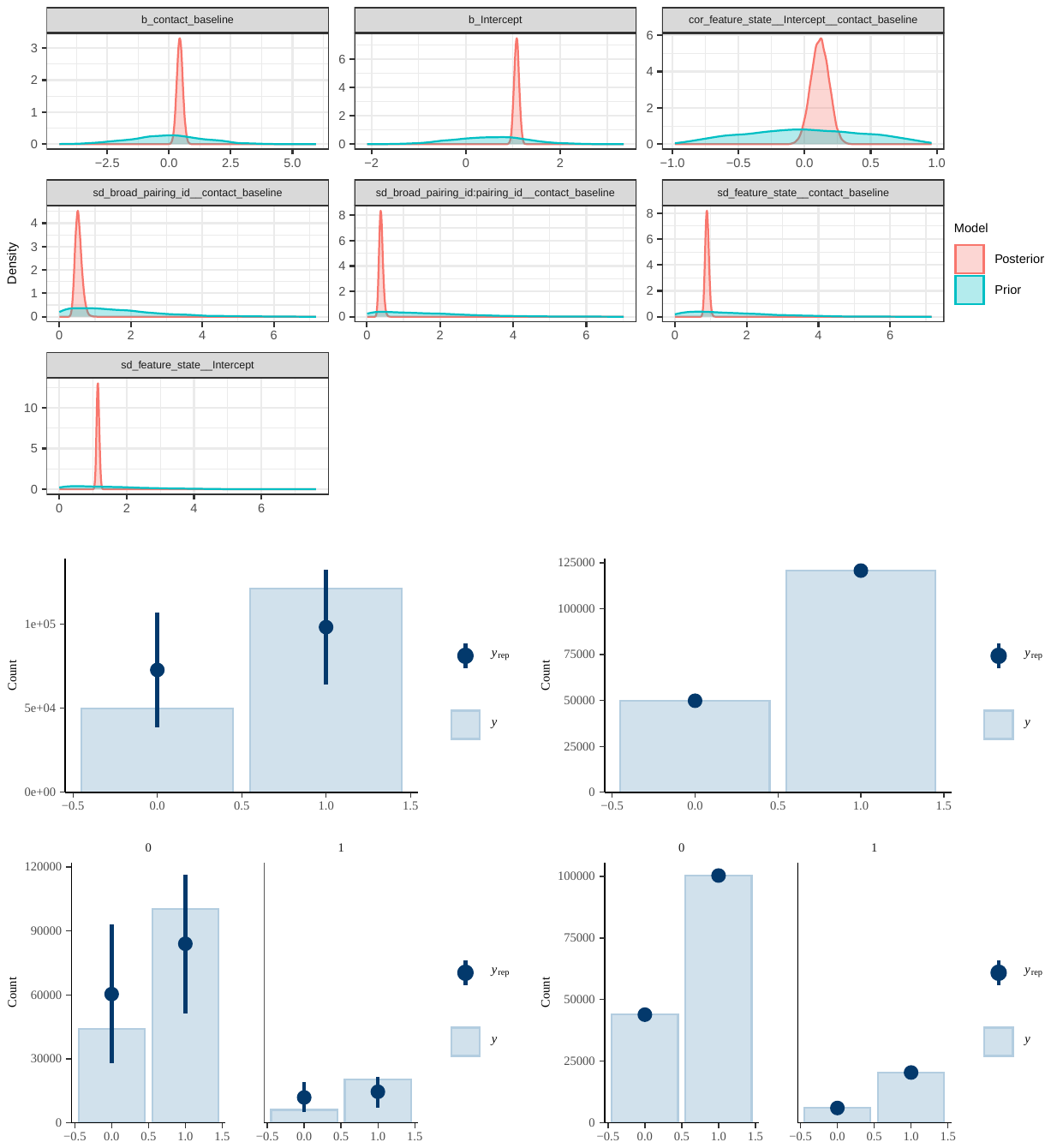
fig. S17.

Prior and posterior distributions for regression coefficients and multilevel hyperparameters (top) as well as prior predictive checks (bottom left) and posterior predictive checks (bottom right) for regression coefficients: genetic model, including all pairs, TLI data.

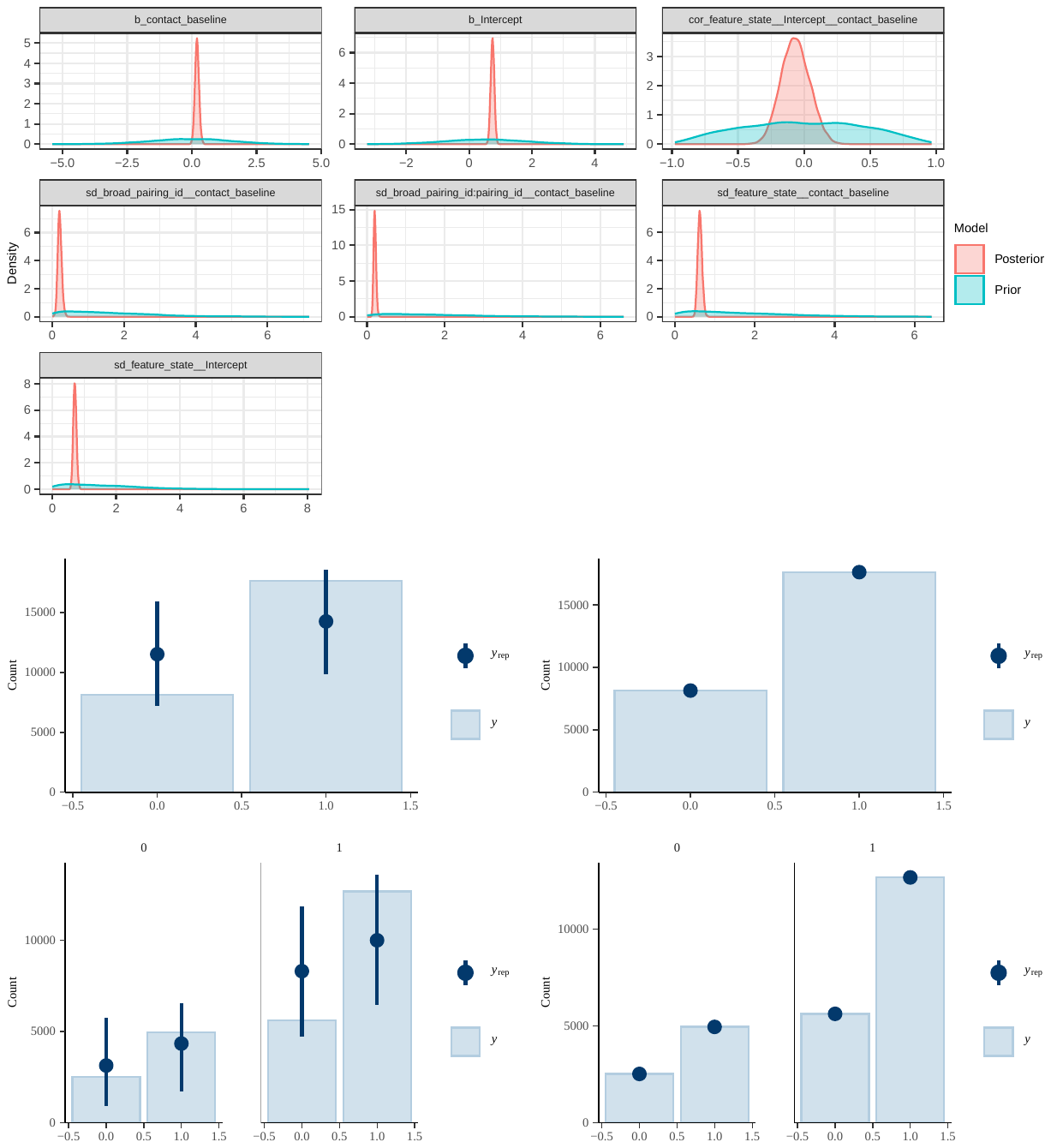
fig. S18.

Prior and posterior distributions for regression coefficients and multilevel hyperparameters (top) as well as prior predictive checks (bottom left) and posterior predictive checks (bottom right) for regression coefficients: genetic model, including only pairs from the same area, GBI data, main analysis (AUTOTYP areas).

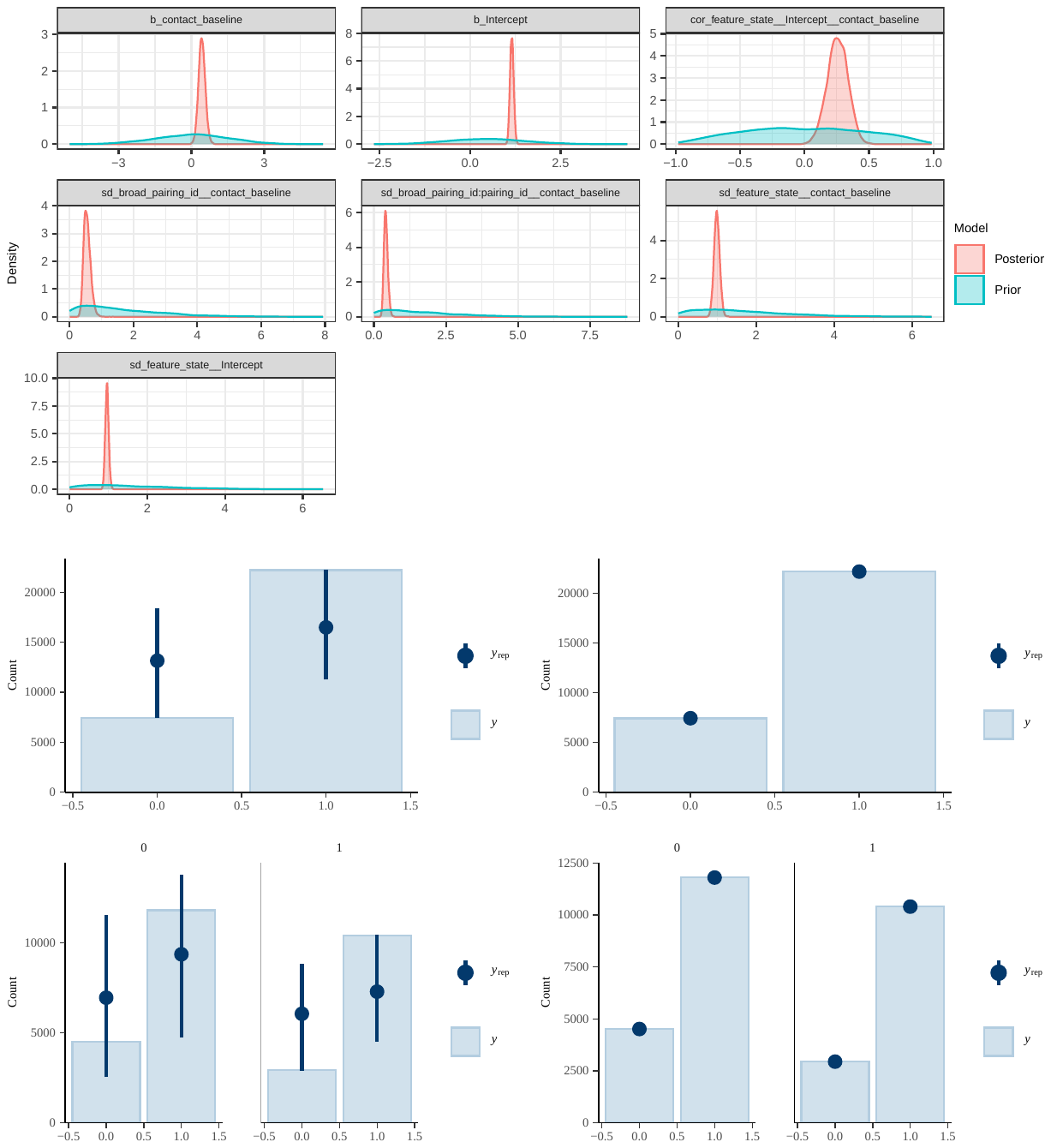
fig. S19.

Prior and posterior distributions for regression coefficients and multilevel hyperparameters (top) as well as prior predictive checks (bottom left) and posterior predictive checks (bottom right) for regression coefficients: genetic model, including only pairs from the same area, TLI data, main analysis (AUTOTYP areas).

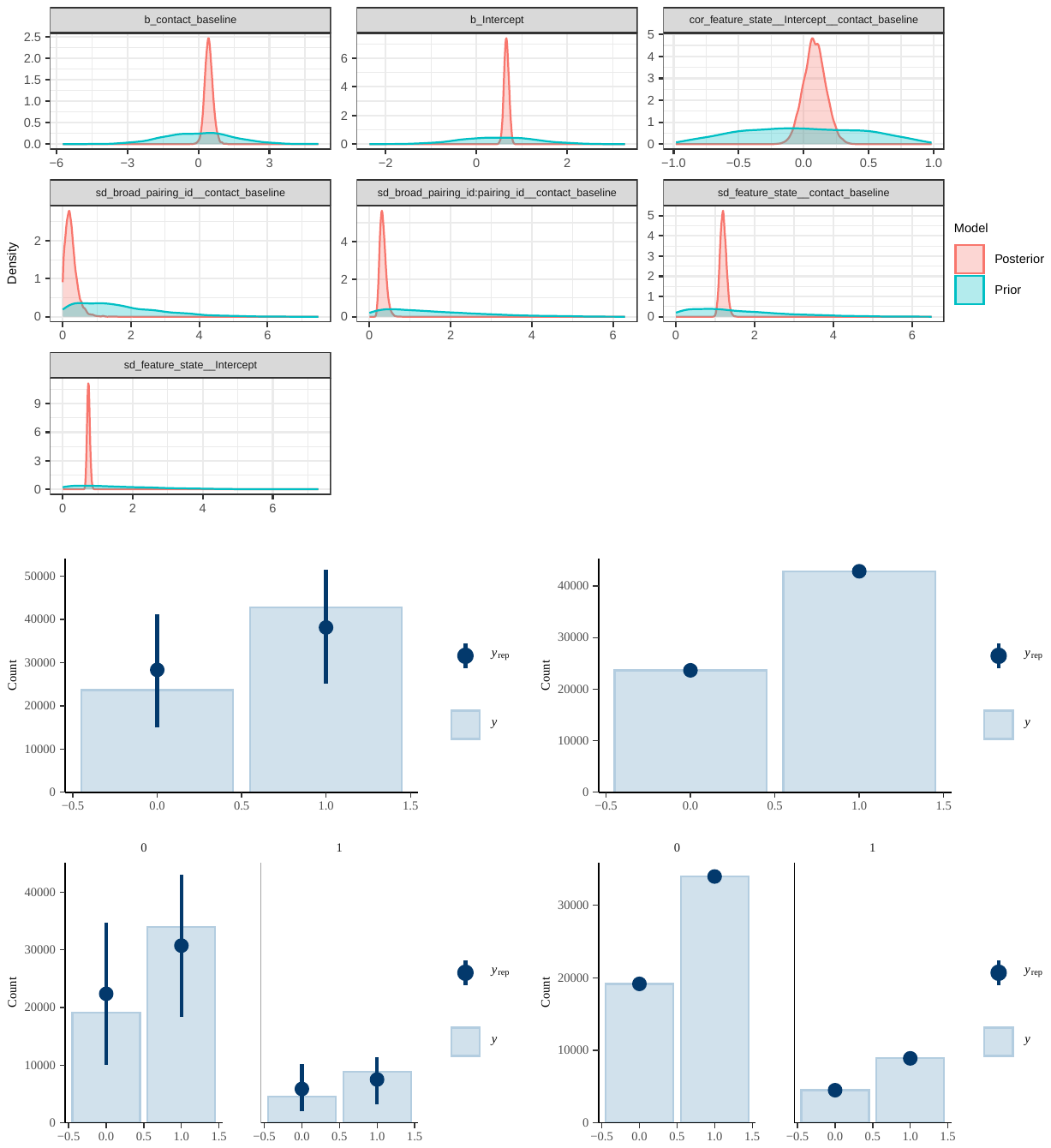
fig. S20.

Prior and posterior distributions for regression coefficients and multilevel hyperparameters (top) as well as prior predictive checks (bottom left) and posterior predictive checks (bottom right) for regression coefficients: genetic model, including only pairs from different areas, GBI data, main analysis (AUTOTYP areas).

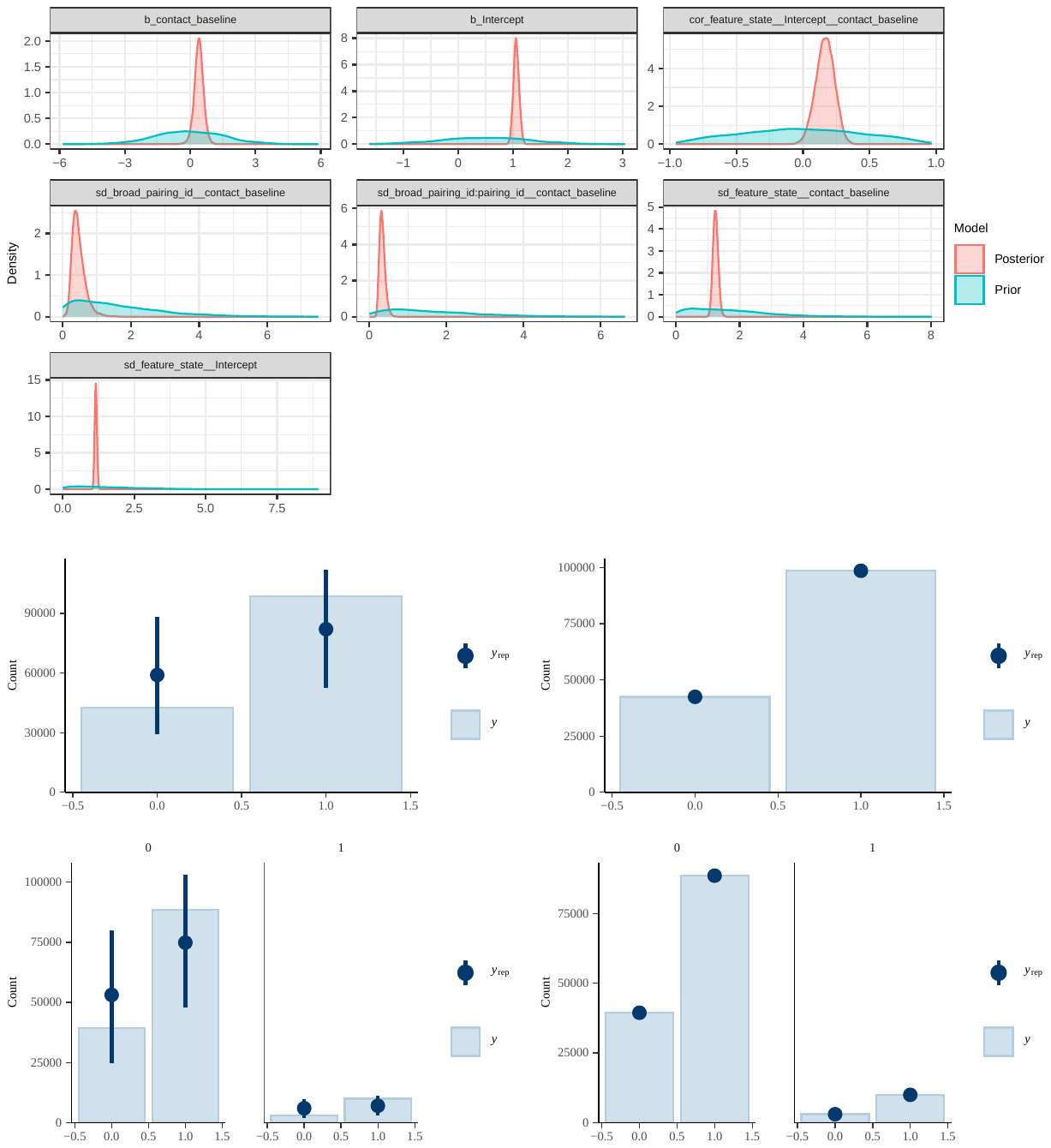
fig. S21.

Prior and posterior distributions for regression coefficients and multilevel hyperparameters (top) as well as prior predictive checks (bottom left) and posterior predictive checks (bottom right) for regression coefficients: genetic model, including only pairs from different areas, TLI data, main analysis (AUTOTYP areas).

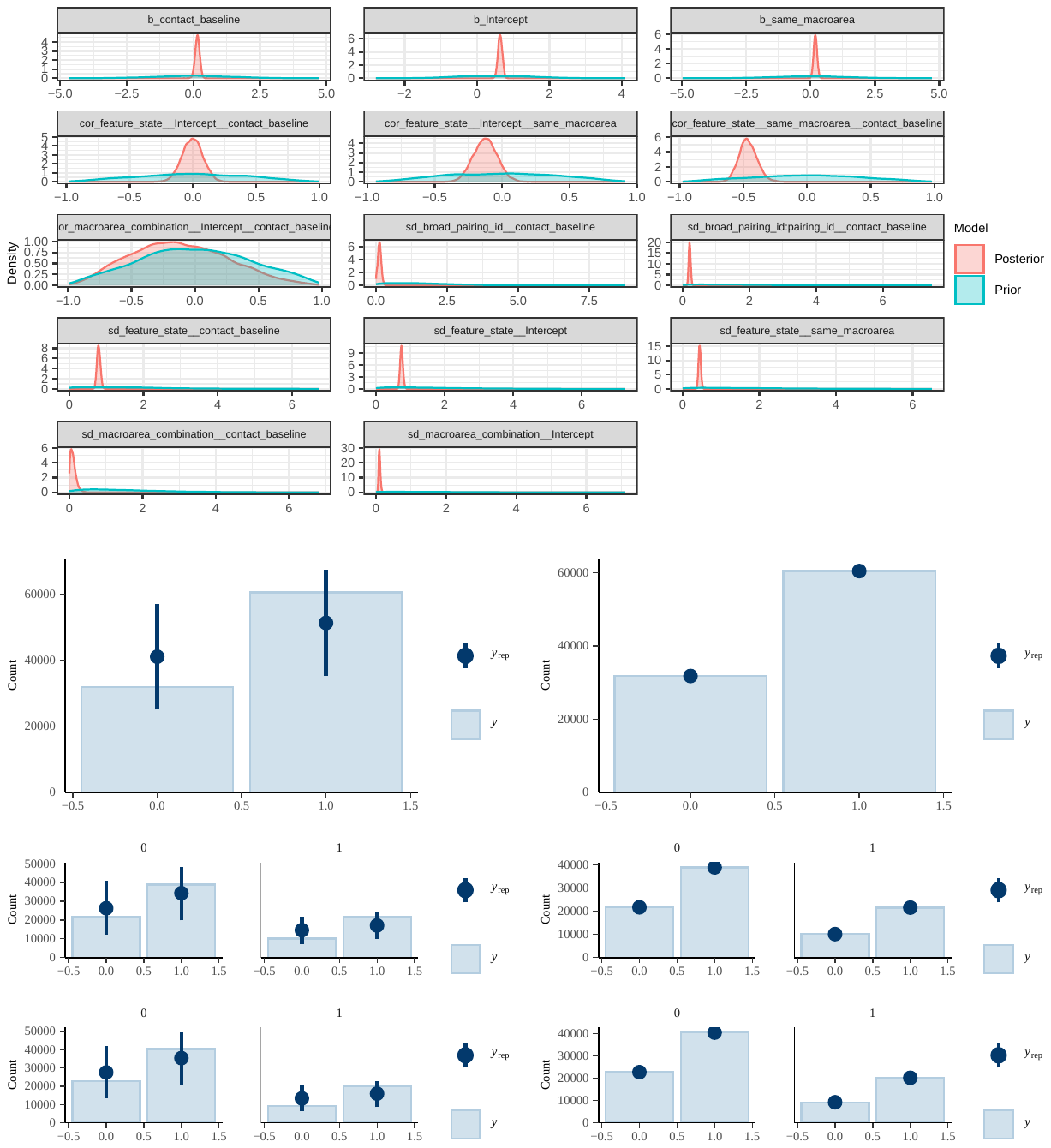
fig. S22.

Prior and posterior distributions for regression coefficients and multilevel hyperparameters (top) as well as prior predictive checks (bottom left) and posterior predictive checks (bottom right) for regression coefficients: combined model (genetic contact and information on areal co-location), including all pairs, GBI data, sensitivity analysis (Glottolog areas).

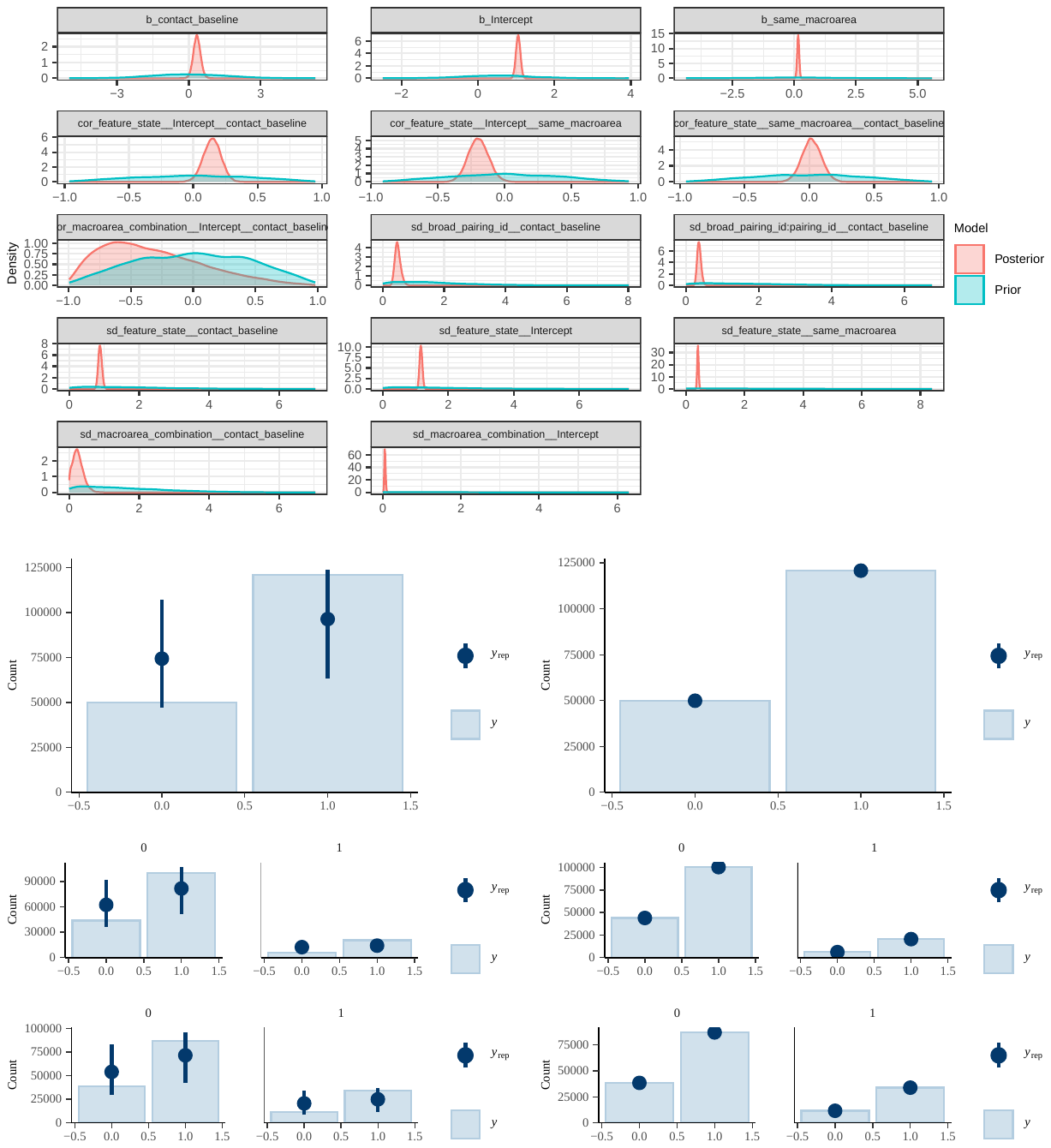
fig. S23.

Prior and posterior distributions for regression coefficients and multilevel hyperparameters (top) as well as prior predictive checks (bottom left) and posterior predictive checks (bottom right) for regression coefficients: combined model (genetic contact and information on areal co-location), including all pairs, TLI data, sensitivity analysis (Glottolog areas).

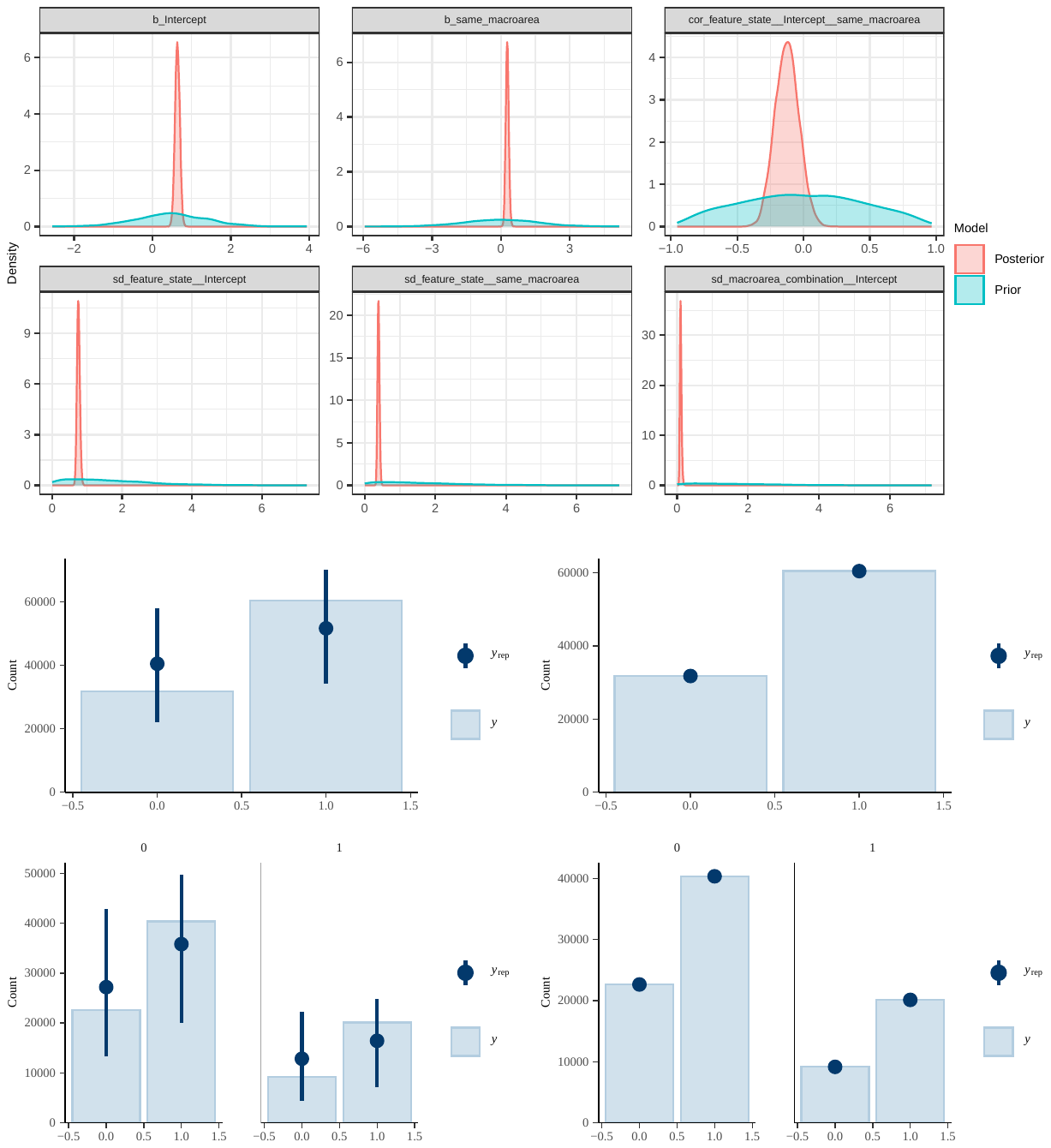
fig. S24.

Prior and posterior distributions for regression coefficients and multilevel hyperparameters (top) as well as prior predictive checks (bottom left) and posterior predictive checks (bottom right) for regression coefficients: areal model, including all pairs, GBI data, sensitivity analysis (Glottolog areas).

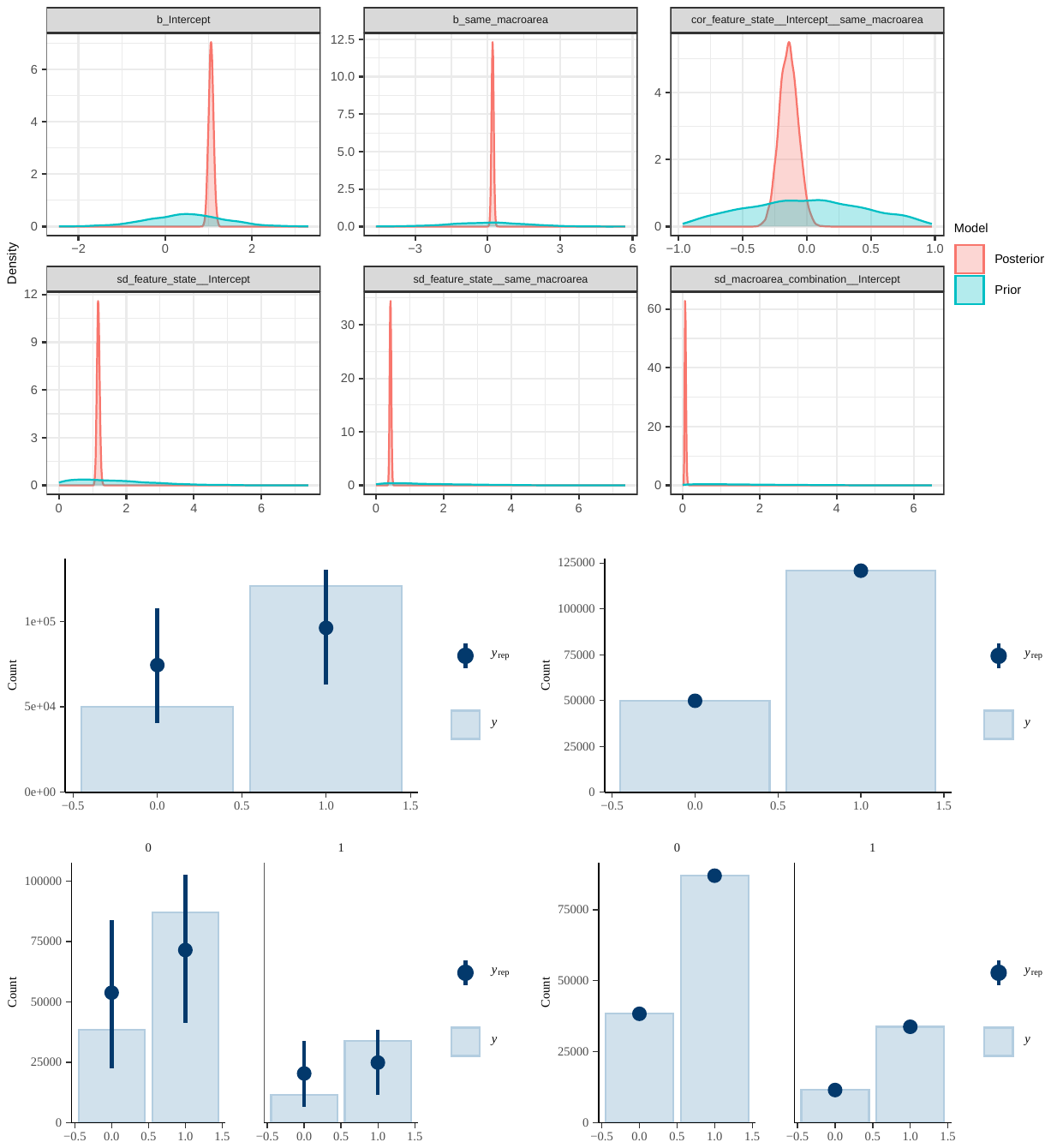
fig. S25.

Prior and posterior distributions for regression coefficients and multilevel hyperparameters (top) as well as prior predictive checks (bottom left) and posterior predictive checks (bottom right) for regression coefficients: areal model, including all pairs, TLI data, sensitivity analysis (Glottolog areas).

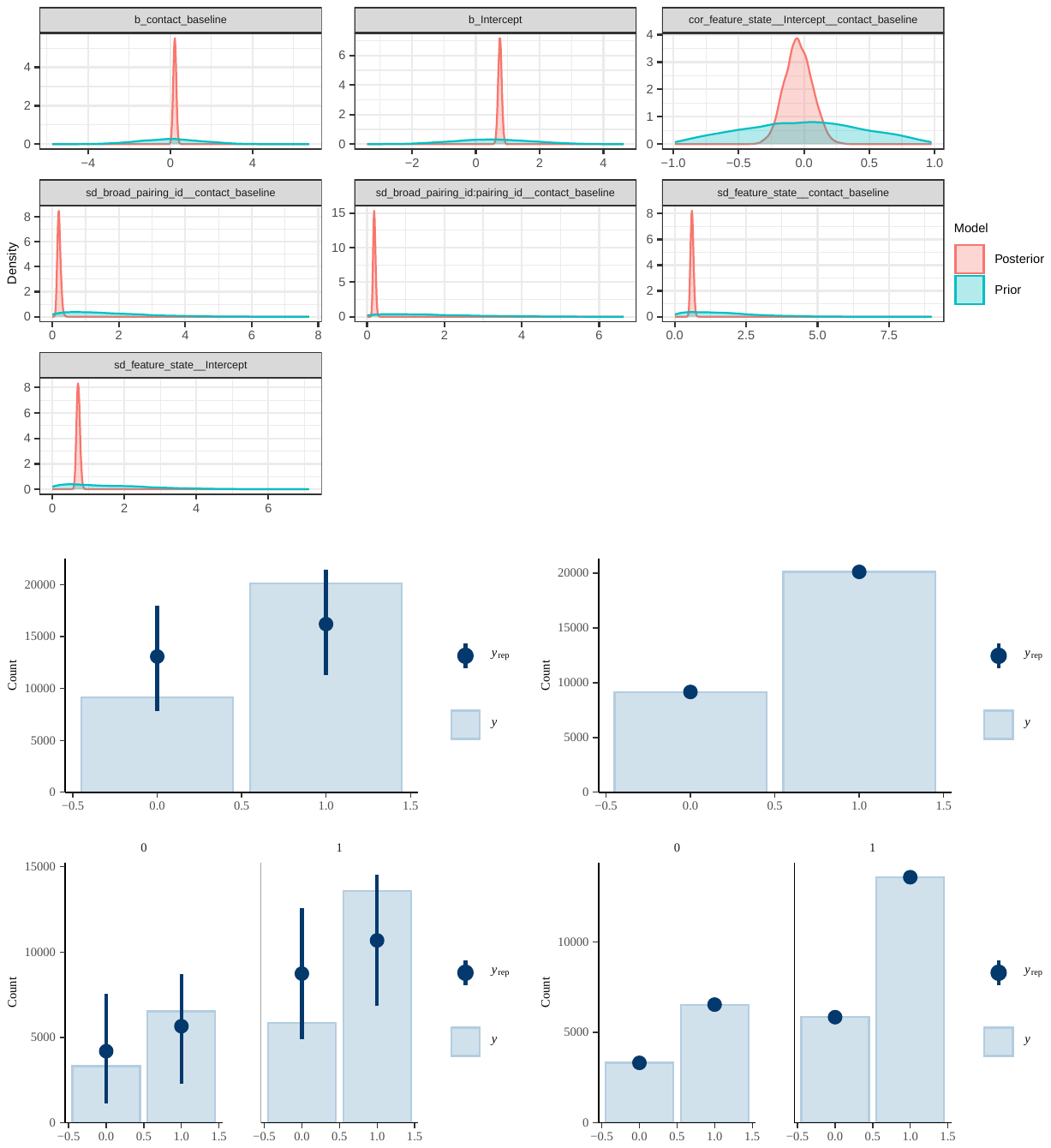
fig. S26.

Prior and posterior distributions for regression coefficients and multilevel hyperparameters (top) as well as prior predictive checks (bottom left) and posterior predictive checks (bottom right) for regression coefficients: genetic model, including only pairs from the same area, GBI data, sensitivity analysis (Glottolog areas).

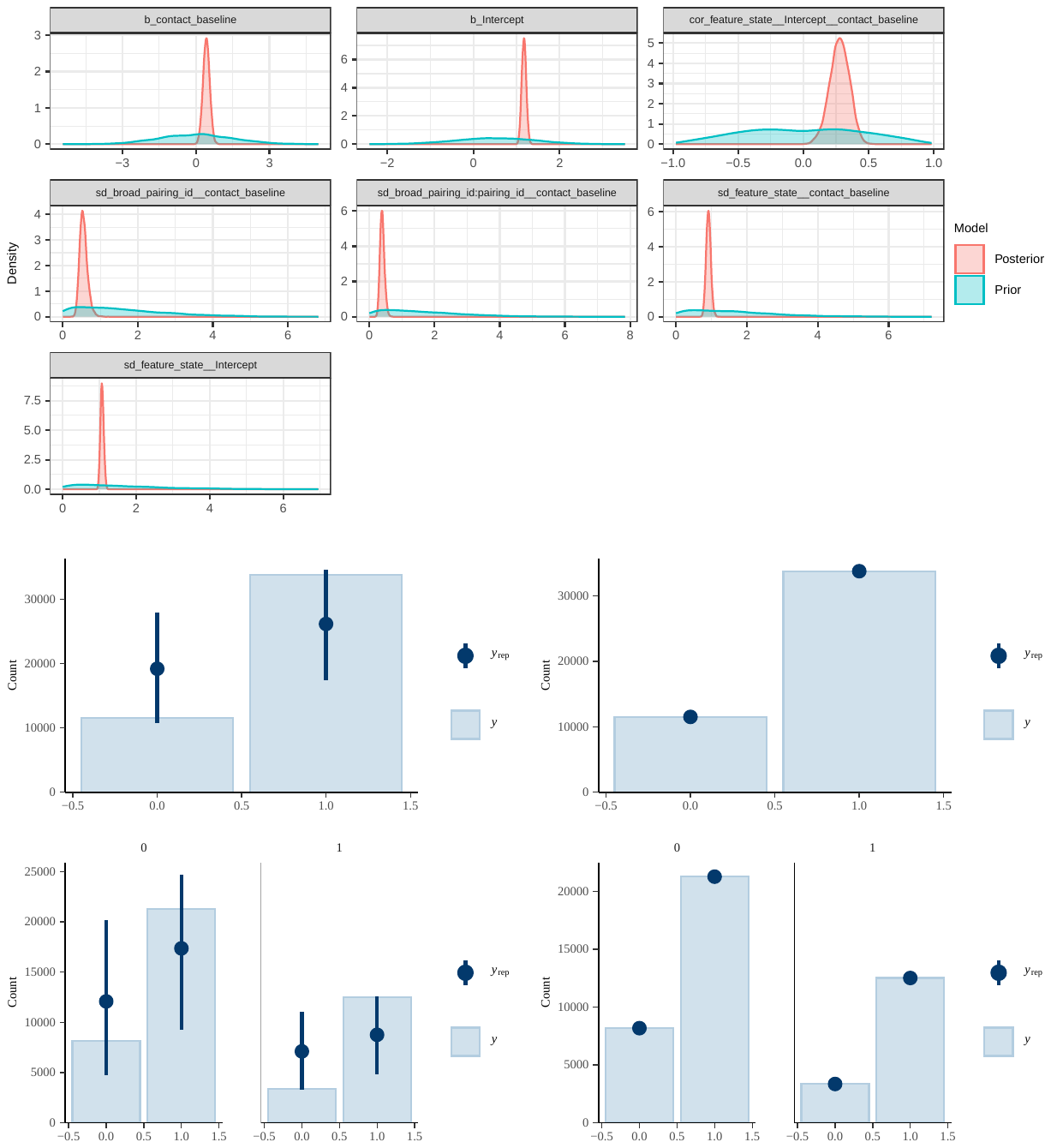
fig. S27.

Prior and posterior distributions for regression coefficients and multilevel hyperparameters (top) as well as prior predictive checks (bottom left) and posterior predictive checks (bottom right) for regression coefficients: genetic model, including only pairs from the same area, TLI data, sensitivity analysis (Glottolog areas).

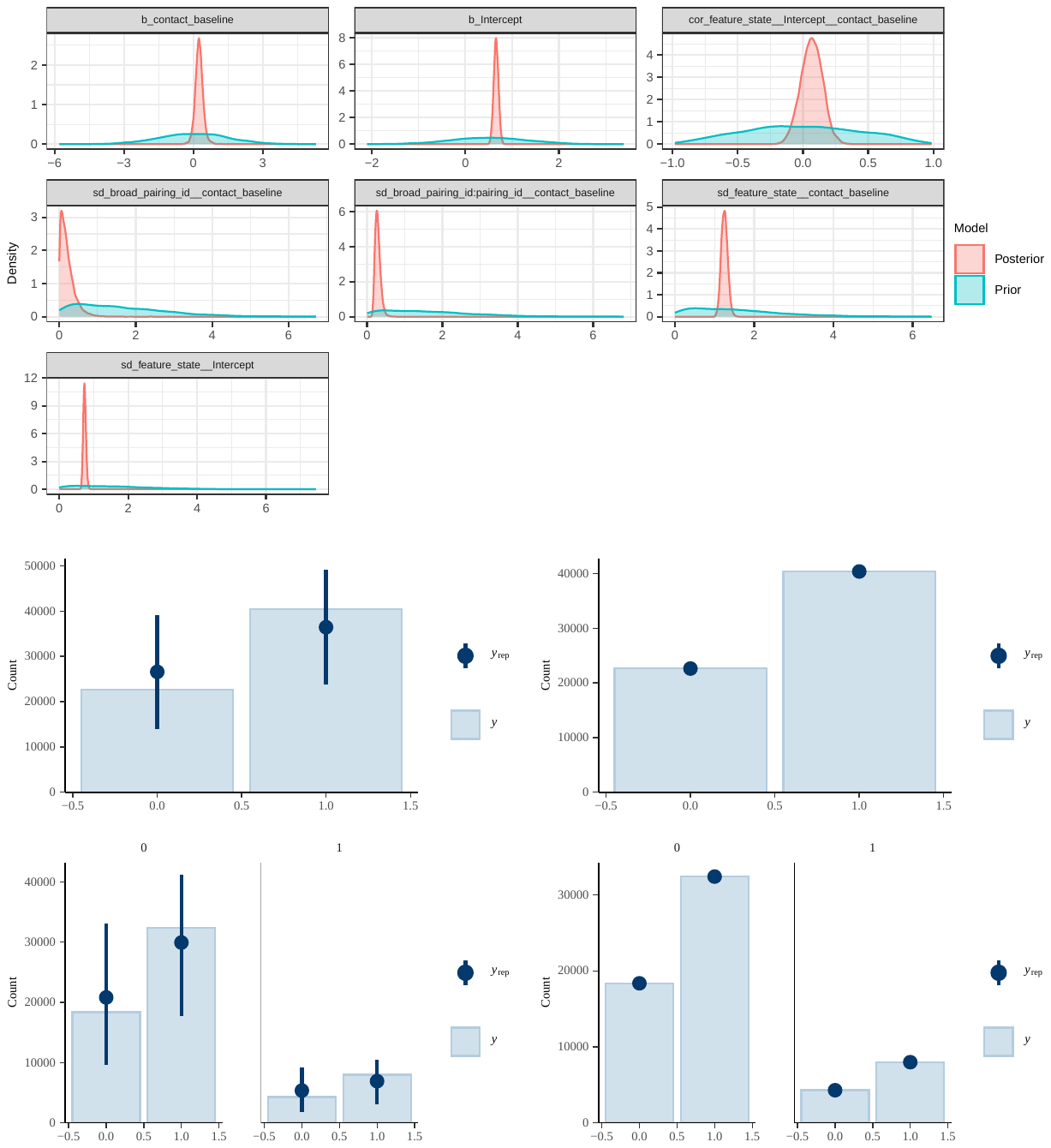
fig. S28.

Prior and posterior distributions for regression coefficients and multilevel hyperparameters (top) as well as prior predictive checks (bottom left) and posterior predictive checks (bottom right) for regression coefficients: genetic model, including only pairs from different areas, GBI data, sensitivity analysis (Glottolog areas).

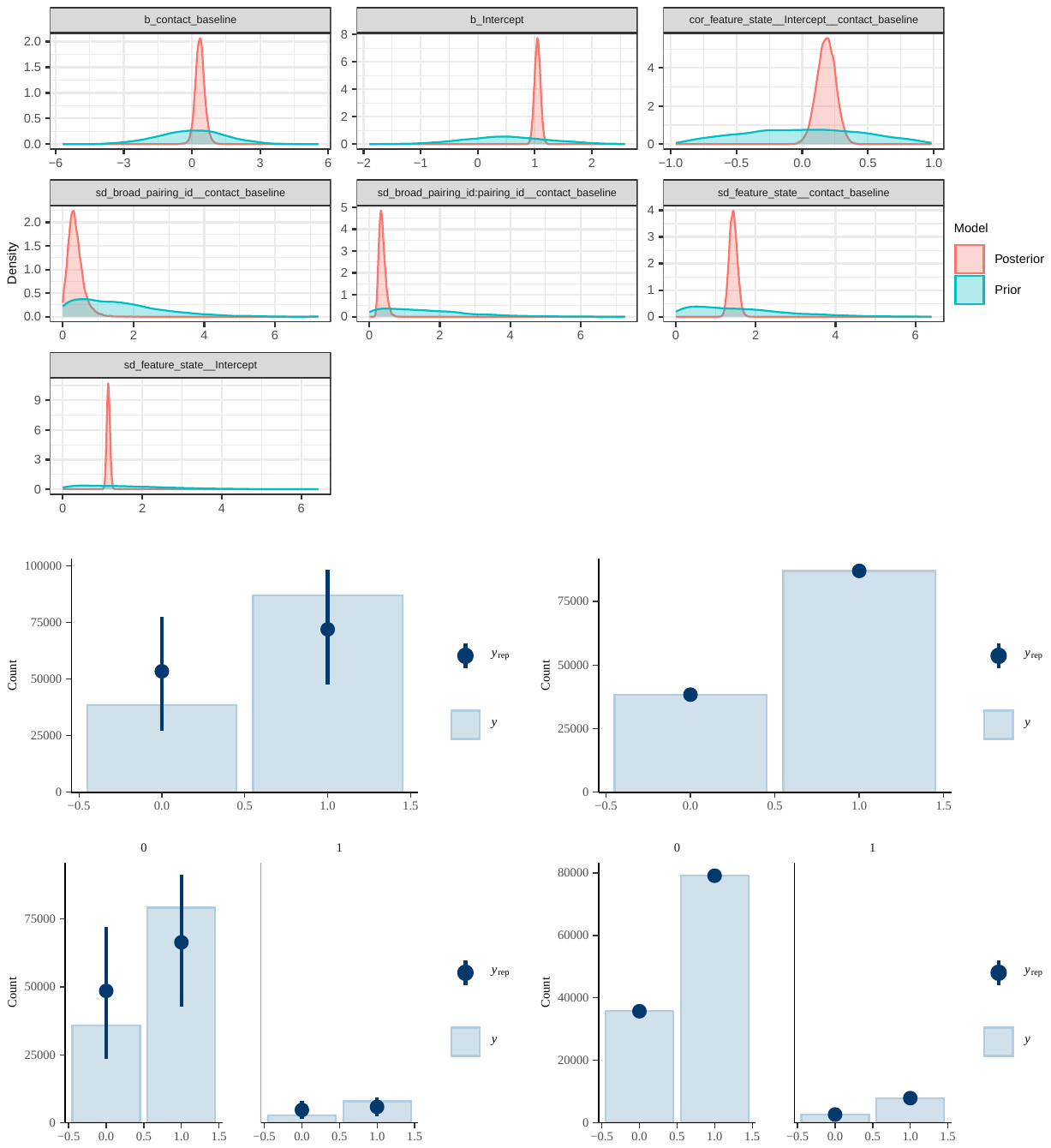
fig. S29.

Prior and posterior distributions for regression coefficients and multilevel hyperparameters (top) as well as prior predictive checks (bottom left) and posterior predictive checks (bottom right) for regression coefficients: genetic model, including only pairs from different areas, TLI data, sensitivity analysis (Glottolog areas).

table S1.

GeLaTo populations used for ADMIXTURE analysis, including number of individuals and language assignments to both datasets used for linguistic analysis.

(external .csv file, tableS1.csv)

table S2.

List of admixed genetic population pairs, including the reference (GeLaTo or a specified publication) and corresponding linguistic pair sets. Each of the 126 pairs has its unique pair id, which is nested within one of 39 broad pair ids, grouping together contact instances with the same source. The table lists language assignments to targets for GBI and TLI individually, and clade assignments to sources (valid for both GBI and TLI). A single language within the clade is picked as a source for GBI and TLI individually for plotting purposes only (Fig. 1C).

(external .csv file, tableS2.csv)

table S3.

Features from GBI-statistical and TLI-statistical and their definitions. The third and fourth columns provide the assignments of the feature groups defined in the databases and explained in the main text.

(external .csv file, tableS3.csv)

table S4.

Summary of fixed and random effects (on log-odds scale) for m1 on the GBI data, using AUTOTYP areas.

m1 (GBI), Fixed Effects Summary

| Coefficient | Estimate | Est.Error | l-89% CI | u-89% CI | Rhat | Bulk_ESS | Tail_ESS |
| --- | --- | --- | --- | --- | --- | --- | --- |
| $\alpha$ | 0.67 | 0.06 | 0.58 | 0.76 | 1.01 | 856 | 1,470 |
| $\beta_{1}(A)$ | 0.16 | 0.06 | 0.06 | 0.26 | 1.00 | 5,822 | 8,762 |
| $\beta_{2}(G)$ | 0.18 | 0.08 | 0.05 | 0.31 | 1.00 | 4,665 | 7,038 |

### Random Effects, Group: broad_pairing_id

| Hyperparameter | Estimate | Est.Error | l-89% CI | u-89% CI | Rhat | Bulk_ESS | Tail_ESS |
| --- | --- | --- | --- | --- | --- | --- | --- |
| $\sigma_{BROAD\_ID}$ | 0.14 | 0.06 | 0.03 | 0.24 | 1 | 2,180 | 2,526 |

Random Effects, Group: broad_pairing_id:pairing_id

| Hyperparameter | Estimate | Est.Error | l-89% CI | u-89% CI | Rhat | Bulk_ESS | Tail_ESS |
| --- | --- | --- | --- | --- | --- | --- | --- |
| $\sigma_{BROAD\_ID/PAIR\_ID}$ | 0.22 | 0.03 | 0.18 | 0.27 | 1 | 3,884 | 8,052 |

Random Effects, Group: continent_combination

| Hyperparameter | Estimate | Est.Error | l-89% CI | u-89% CI | Rhat | Bulk_ESS | Tail_ESS |
| --- | --- | --- | --- | --- | --- | --- | --- |
| ${\sigma_{\alpha}}_{\mathrm{AA}}$ | 0.10 | 0.02 | 0.07 | 0.12 | 1 | 5,981 | 9,814 |
| ${\sigma_{\beta}}_{\mathrm{AA}}$ | 0.07 | 0.06 | 0.01 | 0.19 | 1 | 3,107 | 5,425 |
| ${cor(\alpha}_{\mathrm{AA}}, \beta_{\mathrm{AA}})$ | 0.05 | 0.40 | -0.62 | 0.68 | 1 | 14,651 | 10,753 |

Random Effects, Group: feature_state

| Hyperparameter | Estimate | Est.Error | l-89% CI | u-89% CI | Rhat | Bulk_ESS | Tail_ESS |
| --- | --- | --- | --- | --- | --- | --- | --- |
| ${\sigma_{\alpha}}_{\mathrm{STATE}}$ | 0.75 | 0.04 | 0.69 | 0.82 | 1 | 1,828 | 3,821 |
| ${\sigma_{\beta}}_{A, STATE}$ | 0.51 | 0.03 | 0.46 | 0.57 | 1 | 5,848 | 8,880 |
| ${\sigma_{\beta}}_{G, STATE}$ | 0.82 | 0.05 | 0.75 | 0.90 | 1 | 2,994 | 4,950 |
| ${cor(\alpha}_{\mathrm{STATE}}, \beta_{A,STATE})$ | -0.17 | 0.09 | -0.31 | -0.04 | 1 | 5,694 | 8,748 |
| ${cor(\alpha}_{\mathrm{STATE}}, \beta_{G,STATE})$ | 0.02 | 0.08 | -0.11 | 0.15 | 1 | 3,623 | 6,561 |
| ${cor(\beta}_{A,STATE}, \beta_{G,STATE})$ | -0.56 | 0.06 | -0.65 | -0.46 | 1 | 2,105 | 3,828 |

table S5.

Summary of fixed and random effects (on log-odds scale) for m1 on the TLI data, using AUTOTYP areas.

m1 (TLI), Fixed Effects Summary

| Coefficient | Estimate | Est.Error | l-89% CI | u-89% CI | Rhat | Bulk_ESS | Tail_ESS |
| --- | --- | --- | --- | --- | --- | --- | --- |
| $\alpha$ | 1.06 | 0.05 | 0.97 | 1.15 | 1 | 657 | 1,426 |
| $\beta_{1}(A)$ | 0.16 | 0.04 | 0.10 | 0.23 | 1 | 11,616 | 14,468 |
| $\beta_{2}(G)$ | 0.36 | 0.15 | 0.11 | 0.61 | 1 | 9,689 | 10,611 |

Random Effects, Group: broad_pairing_id

| Hyperparameter | Estimate | Est.Error | l-89% CI | u-89% CI | Rhat | Bulk_ESS | Tail_ESS |
| --- | --- | --- | --- | --- | --- | --- | --- |
| $\sigma_{BROAD\_ID}$ | 0.47 | 0.09 | 0.34 | 0.63 | 1 | 8,188 | 12,714 |

Random Effects, Group: broad_pairing_id:pairing_id

| Hyperparameter | Estimate | Est.Error | l-89% CI | u-89% CI | Rhat | Bulk_ESS | Tail_ESS |
| --- | --- | --- | --- | --- | --- | --- | --- |
| $\sigma_{BROAD\_ID/PAIR\_ID}$ | 0.35 | 0.05 | 0.28 | 0.44 | 1 | 6,963 | 12,433 |

Random Effects, Group: continent_combination

| Hyperparameter | Estimate | Est.Error | l-89% CI | u-89% CI | Rhat | Bulk_ESS | Tail_ESS |
| --- | --- | --- | --- | --- | --- | --- | --- |
| $\alpha_{\mathrm{AA}}$ | 0.07 | 0.01 | 0.05 | 0.09 | 1 | 8,652 | 12,848 |
| $\beta_{\mathrm{AA}}$ | 0.32 | 0.14 | 0.12 | 0.55 | 1 | 4,868 | 5,201 |
| ${cor(\alpha}_{\mathrm{AA}}, \beta_{\mathrm{AA}})$ | -0.11 | 0.33 | -0.62 | 0.43 | 1 | 12,381 | 13,232 |

Random Effects, Group: feature_state

| Hyperparameter | Estimate | Est.Error | l-89% CI | u-89% CI | Rhat | Bulk_ESS | Tail_ESS |
| --- | --- | --- | --- | --- | --- | --- | --- |
| ${\sigma_{\alpha}}_{\mathrm{STATE}}$ | 1.17 | 0.04 | 1.11 | 1.23 | 1 | 1,875 | 3,774 |
| ${\sigma_{\beta}}_{A, STATE}$ | 0.40 | 0.03 | 0.36 | 0.44 | 1 | 9,907 | 14,876 |
| ${\sigma_{\beta}}_{G, STATE}$ | 0.93 | 0.05 | 0.85 | 1.02 | 1 | 5,986 | 10,523 |
| ${cor(\alpha}_{\mathrm{STATE}}, \beta_{A,STATE})$ | -0.31 | 0.08 | -0.44 | -0.18 | 1 | 15,783 | 15,780 |
| ${cor(\alpha}_{\mathrm{STATE}}, \beta_{G,STATE})$ | 0.17 | 0.07 | 0.07 | 0.28 | 1 | 7,971 | 11,944 |
| ${cor(\beta}_{A,STATE}, \beta_{G,STATE})$ | -0.23 | 0.08 | -0.35 | -0.11 | 1 | 2,329 | 4,675 |

table S6.

Summary of fixed and random effects (on log-odds scale) for m2 on the GBI data, using AUTOTYP areas.

m2 (GBI), Fixed Effects Summary

| Coefficient | Estimate | Est.Error | l-89% CI | u-89% CI | Rhat | Bulk_ESS | Tail_ESS |
| --- | --- | --- | --- | --- | --- | --- | --- |
| $\alpha$ | 0.68 | 0.06 | 0.59 | 0.78 | 1.01 | 641 | 1,075 |
| $\beta_{1}(A)$ | 0.25 | 0.06 | 0.16 | 0.35 | 1.00 | 4,462 | 7,397 |

Random Effects, Group: continent_combination

| Hyperparameter | Estimate | Est.Error | l-89% CI | u-89% CI | Rhat | Bulk_ESS | Tail_ESS |
| --- | --- | --- | --- | --- | --- | --- | --- |
| $\sigma_{\mathrm{AA}}$ | 0.12 | 0.02 | 0.09 | 0.15 | 1 | 4,599 | 8,101 |

Random Effects, Group: feature_state

| Hyperparameter | Estimate | Est.Error | l-89% CI | u-89% CI | Rhat | Bulk_ESS | Tail_ESS |
| --- | --- | --- | --- | --- | --- | --- | --- |
| ${\sigma_{\alpha}}_{\mathrm{STATE}}$ | 0.76 | 0.04 | 0.69 | 0.82 | 1 | 1,612 | 3,040 |
| ${\sigma_{\beta}}_{\mathrm{STATE}}$ | 0.42 | 0.03 | 0.37 | 0.47 | 1 | 5,164 | 8,098 |
| ${cor(\alpha}_{\mathrm{STATE}}, \beta_{\mathrm{STATE}})$ | -0.18 | 0.09 | -0.32 | -0.04 | 1 | 4,860 | 8,196 |

table S7.

Summary of fixed and random effects (on log-odds scale) for m2 on the TLI data, using AUTOTYP areas.

m2 (TLI), Fixed Effects Summary

| Coefficient | Estimate | Est.Error | l-89% CI | u-89% CI | Rhat | Bulk_ESS | Tail_ESS |
| --- | --- | --- | --- | --- | --- | --- | --- |
| $\alpha$ | 1.07 | 0.06 | 0.98 | 1.16 | 1 | 1,123 | 2,413 |
| $\beta_{1}(A)$ | 0.21 | 0.05 | 0.13 | 0.28 | 1 | 17,746 | 21,892 |

Random Effects, Group: continent_combination

| Hyperparameter | Estimate | Est.Error | l-89% CI | u-89% CI | Rhat | Bulk_ESS | Tail_ESS |
| --- | --- | --- | --- | --- | --- | --- | --- |
| $\sigma_{\mathrm{AA}}$ | 0.1 | 0.01 | 0.08 | 0.12 | 1 | 12,232 | 19,782 |

Random Effects, Group: feature_state

| Hyperparameter | Estimate | Est.Error | l-89% CI | u-89% CI | Rhat | Bulk_ESS | Tail_ESS |
| --- | --- | --- | --- | --- | --- | --- | --- |
| ${\sigma_{\alpha}}_{\mathrm{STATE}}$ | 1.17 | 0.04 | 1.11 | 1.24 | 1 | 2,984 | 6,116 |
| ${\sigma_{\beta}}_{\mathrm{STATE}}$ | 0.41 | 0.02 | 0.37 | 0.45 | 1 | 14,751 | 21,033 |
| ${cor(\alpha}_{\mathrm{STATE}}, \beta_{\mathrm{STATE}})$ | -0.25 | 0.08 | -0.38 | -0.12 | 1 | 24,507 | 23,804 |

table S8.

Summary of fixed and random effects (on log-odds scale) for m3 on the GBI data, using AUTOTYP areas.

m3 (GBI), Fixed Effects Summary

| Coefficient | Estimate | Est.Error | l-89% CI | u-89% CI | Rhat | Bulk_ESS | Tail_ESS |
| --- | --- | --- | --- | --- | --- | --- | --- |
| $\alpha$ | 0.67 | 0.05 | 0.59 | 0.75 | 1 | 768 | 1,517 |
| $\beta_{1}(G)$ | 0.27 | 0.07 | 0.15 | 0.39 | 1 | 2,614 | 4,969 |

Random Effects, Group: broad_pairing_id

| Hyperparameter | Estimate | Est.Error | l-89% CI | u-89% CI | Rhat | Bulk_ESS | Tail_ESS |
| --- | --- | --- | --- | --- | --- | --- | --- |
| $\sigma_{BROAD\_ID}$ | 0.19 | 0.05 | 0.1 | 0.27 | 1 | 2,530 | 2,782 |

Random Effects, Group: broad_pairing_id:pairing_id

| Hyperparameter | Estimate | Est.Error | l-89% CI | u-89% CI | Rhat | Bulk_ESS | Tail_ESS |
| --- | --- | --- | --- | --- | --- | --- | --- |
| $\sigma_{BROAD\_ID/PAIR\_ID}$ | 0.23 | 0.03 | 0.19 | 0.28 | 1 | 3,805 | 6,022 |

Random Effects, Group: feature_state

| Hyperparameter | Estimate | Est.Error | l-89% CI | u-89% CI | Rhat | Bulk_ESS | Tail_ESS |
| --- | --- | --- | --- | --- | --- | --- | --- |
| ${\sigma_{\alpha}}_{\mathrm{STATE}}$ | 0.73 | 0.04 | 0.68 | 0.80 | 1 | 1,547 | 3,405 |
| ${\sigma_{\beta}}_{\mathrm{STATE}}$ | 0.72 | 0.04 | 0.65 | 0.79 | 1 | 3,036 | 5,619 |
| ${cor(\alpha}_{\mathrm{STATE}}, \beta_{\mathrm{STATE}})$ | -0.06 | 0.08 | -0.19 | 0.08 | 1 | 2,352 | 4,154 |

table S9.

Summary of fixed and random effects (on log-odds scale) for m3 on the TLI data, using AUTOTYP areas.

m3 (TLI), Fixed Effects Summary

| Coefficient | Estimate | Est.Error | l-89% CI | u-89% CI | Rhat | Bulk_ESS | Tail_ESS |
| --- | --- | --- | --- | --- | --- | --- | --- |
| $\alpha$ | 1.08 | 0.05 | 0.99 | 1.16 | 1 | 649 | 1,637 |
| $\beta_{1}(G)$ | 0.44 | 0.12 | 0.25 | 0.63 | 1 | 8,350 | 13,726 |

Random Effects, Group: broad_pairing_id

| Hyperparameter | Estimate | Est.Error | l-89% CI | u-89% CI | Rhat | Bulk_ESS | Tail_ESS |
| --- | --- | --- | --- | --- | --- | --- | --- |
| $\sigma_{BROAD\_ID}$ | 0.54 | 0.09 | 0.4 | 0.7 | 1 | 9,850 | 15,545 |

Random Effects, Group: broad_pairing_id:pairing_id

| Hyperparameter | Estimate | Est.Error | l-89% CI | u-89% CI | Rhat | Bulk_ESS | Tail_ESS |
| --- | --- | --- | --- | --- | --- | --- | --- |
| $\sigma_{BROAD\_ID/PAIR\_ID}$ | 0.39 | 0.05 | 0.31 | 0.47 | 1 | 9,283 | 16,254 |

Random Effects, Group: feature_state

| Hyperparameter | Estimate | Est.Error | l-89% CI | u-89% CI | Rhat | Bulk_ESS | Tail_ESS |
| --- | --- | --- | --- | --- | --- | --- | --- |
| ${\sigma_{\alpha}}_{\mathrm{STATE}}$ | 1.15 | 0.04 | 1.09 | 1.21 | 1 | 1,812 | 4,497 |
| ${\sigma_{\beta}}_{\mathrm{STATE}}$ | 0.90 | 0.05 | 0.82 | 0.98 | 1 | 8,537 | 15,728 |
| ${cor(\alpha}_{\mathrm{STATE}}, \beta_{\mathrm{STATE}})$ | 0.12 | 0.07 | 0.01 | 0.22 | 1 | 9,248 | 15,241 |

table S10.

Summary of fixed and random effects (on log-odds scale) for m4 on the GBI data, using AUTOTYP areas.

m4 (GBI), Fixed Effects Summary

| Coefficient | Estimate | Est.Error | l-89% CI | u-89% CI | Rhat | Bulk_ESS | Tail_ESS |
| --- | --- | --- | --- | --- | --- | --- | --- |
| $\alpha$ | 0.74 | 0.06 | 0.65 | 0.83 | 1 | 3,835 | 4,717 |
| $\beta_{1}(G)$ | 0.20 | 0.08 | 0.08 | 0.32 | 1 | 5,435 | 5,744 |

Random Effects, Group: broad_pairing_id

| Hyperparameter | Estimate | Est.Error | l-89% CI | u-89% CI | Rhat | Bulk_ESS | Tail_ESS |
| --- | --- | --- | --- | --- | --- | --- | --- |
| $\sigma_{BROAD\_ID}$ | 0.21 | 0.06 | 0.13 | 0.3 | 1 | 3,037 | 3,811 |

Random Effects, Group: broad_pairing_id:pairing_id

| Hyperparameter | Estimate | Est.Error | l-89% CI | u-89% CI | Rhat | Bulk_ESS | Tail_ESS |
| --- | --- | --- | --- | --- | --- | --- | --- |
| $\sigma_{BROAD\_ID/PAIR\_ID}$ | 0.2 | 0.03 | 0.16 | 0.25 | 1 | 3,184 | 4,680 |

Random Effects, Group: feature_state

| Hyperparameter | Estimate | Est.Error | l-89% CI | u-89% CI | Rhat | Bulk_ESS | Tail_ESS |
| --- | --- | --- | --- | --- | --- | --- | --- |
| ${\sigma_{\alpha}}_{\mathrm{STATE}}$ | 0.71 | 0.05 | 0.63 | 0.79 | 1 | 4,275 | 5,935 |
| ${\sigma_{\beta}}_{\mathrm{STATE}}$ | 0.63 | 0.05 | 0.55 | 0.72 | 1 | 3,203 | 5,464 |
| ${cor(\alpha}_{\mathrm{STATE}}, \beta_{\mathrm{STATE}})$ | -0.06 | 0.11 | -0.23 | 0.11 | 1 | 3,651 | 4,695 |

table S11.

Summary of fixed and random effects (on log-odds scale) for m4 on the TLI data, using AUTOTYP areas.

m4 (TLI), Fixed Effects Summary

| Coefficient | Estimate | Est.Error | l-89% CI | u-89% CI | Rhat | Bulk_ESS | Tail_ESS |
| --- | --- | --- | --- | --- | --- | --- | --- |
| $\alpha$ | 1.15 | 0.05 | 1.08 | 1.23 | 1 | 2,021 | 3,666 |
| $\beta_{1}(G)$ | 0.43 | 0.14 | 0.22 | 0.66 | 1 | 3,351 | 4,406 |

Random Effects, Group: broad_pairing_id

| Hyperparameter | Estimate | Est.Error | l-89% CI | u-89% CI | Rhat | Bulk_ESS | Tail_ESS |
| --- | --- | --- | --- | --- | --- | --- | --- |
| $\sigma_{BROAD\_ID}$ | 0.55 | 0.11 | 0.4 | 0.73 | 1 | 3,082 | 5,109 |

Random Effects, Group: broad_pairing_id:pairing_id

| Hyperparameter | Estimate | Est.Error | l-89% CI | u-89% CI | Rhat | Bulk_ESS | Tail_ESS |
| --- | --- | --- | --- | --- | --- | --- | --- |
| $\sigma_{BROAD\_ID/PAIR\_ID}$ | 0.42 | 0.07 | 0.32 | 0.53 | 1 | 2,711 | 4,140 |

Random Effects, Group: feature_state

| Hyperparameter | Estimate | Est.Error | l-89% CI | u-89% CI | Rhat | Bulk_ESS | Tail_ESS |
| --- | --- | --- | --- | --- | --- | --- | --- |
| ${\sigma_{\alpha}}_{\mathrm{STATE}}$ | 0.96 | 0.04 | 0.90 | 1.03 | 1 | 2,891 | 4,546 |
| ${\sigma_{\beta}}_{\mathrm{STATE}}$ | 0.99 | 0.07 | 0.88 | 1.11 | 1 | 3,309 | 5,251 |
| ${cor(\alpha}_{\mathrm{STATE}}, \beta_{\mathrm{STATE}})$ | 0.26 | 0.08 | 0.13 | 0.38 | 1 | 3,487 | 4,811 |

table S12.

Summary of fixed and random effects (on log-odds scale) for m5 on the GBI data, using AUTOTYP areas.

m5 (GBI), Fixed Effects Summary

| Coefficient | Estimate | Est.Error | l-89% CI | u-89% CI | Rhat | Bulk_ESS | Tail_ESS |
| --- | --- | --- | --- | --- | --- | --- | --- |
| $\alpha$ | 0.66 | 0.05 | 0.58 | 0.75 | 1 | 451 | 976 |
| $\beta_{1}(G)$ | 0.44 | 0.17 | 0.18 | 0.72 | 1 | 1,929 | 3,443 |

Random Effects, Group: broad_pairing_id

| Hyperparameter | Estimate | Est.Error | l-89% CI | u-89% CI | Rhat | Bulk_ESS | Tail_ESS |
| --- | --- | --- | --- | --- | --- | --- | --- |
| $\sigma_{BROAD\_ID}$ | 0.25 | 0.18 | 0.03 | 0.55 | 1 | 1,965 | 2,624 |

Random Effects, Group: broad_pairing_id:pairing_id

| Hyperparameter | Estimate | Est.Error | l-89% CI | u-89% CI | Rhat | Bulk_ESS | Tail_ESS |
| --- | --- | --- | --- | --- | --- | --- | --- |
| $\sigma_{BROAD\_ID/PAIR\_ID}$ | 0.33 | 0.08 | 0.23 | 0.47 | 1 | 2,972 | 5,787 |

Random Effects, Group: feature_state

| Hyperparameter | Estimate | Est.Error | l-89% CI | u-89% CI | Rhat | Bulk_ESS | Tail_ESS |
| --- | --- | --- | --- | --- | --- | --- | --- |
| ${\sigma_{\alpha}}_{\mathrm{STATE}}$ | 0.74 | 0.04 | 0.68 | 0.80 | 1.01 | 1,155 | 2,385 |
| ${\sigma_{\beta}}_{\mathrm{STATE}}$ | 1.20 | 0.08 | 1.09 | 1.33 | 1.00 | 1,946 | 4,125 |
| ${cor(\alpha}_{\mathrm{STATE}}, \beta_{\mathrm{STATE}})$ | 0.08 | 0.08 | -0.05 | 0.22 | 1.00 | 959 | 2,068 |

table S13.

Summary of fixed and random effects (on log-odds scale) for m5 on the TLI data.

m5 (TLI), Fixed Effects Summary

| Coefficient | Estimate | Est.Error | l-89% CI | u-89% CI | Rhat | Bulk_ESS | Tail_ESS |
| --- | --- | --- | --- | --- | --- | --- | --- |
| $\alpha$ | 1.06 | 0.05 | 0.98 | 1.14 | 1.01 | 462 | 880 |
| $\beta_{1}(G)$ | 0.40 | 0.21 | 0.06 | 0.73 | 1.00 | 5,850 | 8,863 |

Random Effects, Group: broad_pairing_id

| Hyperparameter | Estimate | Est.Error | l-89% CI | u-89% CI | Rhat | Bulk_ESS | Tail_ESS |
| --- | --- | --- | --- | --- | --- | --- | --- |
| $\sigma_{BROAD\_ID}$ | 0.47 | 0.2 | 0.23 | 0.83 | 1 | 5,560 | 8,279 |

Random Effects, Group: broad_pairing_id:pairing_id

| Hyperparameter | Estimate | Est.Error | l-89% CI | u-89% CI | Rhat | Bulk_ESS | Tail_ESS |
| --- | --- | --- | --- | --- | --- | --- | --- |
| $\sigma_{BROAD\_ID/PAIR\_ID}$ | 0.34 | 0.07 | 0.24 | 0.47 | 1 | 5,358 | 9,593 |

Random Effects, Group: feature_state

| Hyperparameter | Estimate | Est.Error | l-89% CI | u-89% CI | Rhat | Bulk_ESS | Tail_ESS |
| --- | --- | --- | --- | --- | --- | --- | --- |
| ${\sigma_{\alpha}}_{\mathrm{STATE}}$ | 1.16 | 0.04 | 1.10 | 1.22 | 1 | 1,296 | 2,612 |
| ${\sigma_{\beta}}_{\mathrm{STATE}}$ | 1.24 | 0.08 | 1.12 | 1.38 | 1 | 4,917 | 9,436 |
| ${cor(\alpha}_{\mathrm{STATE}}, \beta_{\mathrm{STATE}})$ | 0.17 | 0.07 | 0.05 | 0.27 | 1 | 4,896 | 9,415 |

table S14.

Summary of fixed and random effects (on log-odds scale) for m1 on the GBI data, using Glottolog areas (sensitivity analysis).

m1 (GBI), Fixed Effects Summary

| Coefficient | Estimate | Est.Error | l-89% CI | u-89% CI | Rhat | Bulk_ESS | Tail_ESS |
| --- | --- | --- | --- | --- | --- | --- | --- |
| $\alpha$ | 0.64 | 0.06 | 0.54 | 0.73 | 1.01 | 709 | 1,207 |
| $\beta_{1}(A)$ | 0.18 | 0.07 | 0.08 | 0.30 | 1.00 | 4,332 | 5,307 |
| $\beta_{2}(G)$ | 0.15 | 0.09 | 0.01 | 0.29 | 1.00 | 3,225 | 4,796 |

Random Effects, Group: broad_pairing_id

| Hyperparameter | Estimate | Est.Error | l-89% CI | u-89% CI | Rhat | Bulk_ESS | Tail_ESS |
| --- | --- | --- | --- | --- | --- | --- | --- |
| $\sigma_{BROAD\_ID}$ | 0.12 | 0.06 | 0.03 | 0.22 | 1 | 1,350 | 2,011 |

Random Effects, Group: broad_pairing_id:pairing_id

| Hyperparameter | Estimate | Est.Error | l-89% CI | u-89% CI | Rhat | Bulk_ESS | Tail_ESS |
| --- | --- | --- | --- | --- | --- | --- | --- |
| $\sigma_{BROAD\_ID/PAIR\_ID}$ | 0.21 | 0.03 | 0.17 | 0.26 | 1 | 2,925 | 4,643 |

Random Effects, Group: feature_state

| Hyperparameter | Estimate | Est.Error | l-89% CI | u-89% CI | Rhat | Bulk_ESS | Tail_ESS |
| --- | --- | --- | --- | --- | --- | --- | --- |
| ${\sigma_{\alpha}}_{\mathrm{STATE}}$ | 0.75 | 0.04 | 0.69 | 0.81 | 1 | 1,298 | 2,808 |
| ${\sigma_{\beta}}_{A, STATE}$ | 0.45 | 0.03 | 0.40 | 0.50 | 1 | 4,258 | 6,503 |
| ${\sigma_{\beta}}_{G, STATE}$ | 0.79 | 0.05 | 0.72 | 0.87 | 1 | 1,870 | 4,232 |
| ${cor(\alpha}_{\mathrm{STATE}}, \beta_{A,STATE})$ | -0.12 | 0.09 | -0.26 | 0.02 | 1 | 4,246 | 6,403 |
| ${cor(\alpha}_{\mathrm{STATE}}, \beta_{G,STATE})$ | 0.00 | 0.08 | -0.13 | 0.13 | 1 | 2,419 | 4,197 |
| ${cor(\beta}_{A,STATE}, \beta_{G,STATE})$ | -0.46 | 0.07 | -0.57 | -0.35 | 1 | 1,102 | 2,653 |

Random Effects, Group: macroarea_combination

| Hyperparameter | Estimate | Est.Error | l-89% CI | u-89% CI | Rhat | Bulk_ESS | Tail_ESS |
| --- | --- | --- | --- | --- | --- | --- | --- |
| $\sigma_{\mathrm{AA}}$ | 0.10 | 0.02 | 0.07 | 0.14 | 1 | 3,634 | 5,218 |
| $\beta_{\mathrm{AA}}$ | 0.11 | 0.08 | 0.01 | 0.25 | 1 | 2,120 | 4,624 |
| ${cor(\alpha}_{\mathrm{AA}}, \beta_{\mathrm{AA}})$ | -0.12 | 0.38 | -0.70 | 0.52 | 1 | 8,588 | 6,991 |

table S15.

Summary of fixed and random effects (on log-odds scale) for m1 on the TLI data, using Glottolog areas (sensitivity analysis).

m1 (TLI), Fixed Effects Summary

| Coefficient | Estimate | Est.Error | l-89% CI | u-89% CI | Rhat | Bulk_ESS | Tail_ESS |
| --- | --- | --- | --- | --- | --- | --- | --- |
| $\alpha$ | 1.05 | 0.06 | 0.96 | 1.14 | 1.01 | 439 | 955 |
| $\beta_{1}(A)$ | 0.16 | 0.04 | 0.10 | 0.23 | 1.00 | 8,074 | 8,862 |
| $\beta_{2}(G)$ | 0.34 | 0.15 | 0.10 | 0.58 | 1.00 | 7,101 | 8,356 |

Random Effects, Group: broad_pairing_id

| Hyperparameter | Estimate | Est.Error | l-89% CI | u-89% CI | Rhat | Bulk_ESS | Tail_ESS |
| --- | --- | --- | --- | --- | --- | --- | --- |
| $\sigma_{BROAD\_ID}$ | 0.49 | 0.09 | 0.36 | 0.65 | 1 | 6,496 | 9,548 |

Random Effects, Group: broad_pairing_id:pairing_id

| Hyperparameter | Estimate | Est.Error | l-89% CI | u-89% CI | Rhat | Bulk_ESS | Tail_ESS |
| --- | --- | --- | --- | --- | --- | --- | --- |
| $\sigma_{BROAD\_ID/PAIR\_ID}$ | 0.36 | 0.05 | 0.28 | 0.45 | 1 | 4,418 | 8,381 |

Random Effects, Group: feature_state

| Hyperparameter | Estimate | Est.Error | l-89% CI | u-89% CI | Rhat | Bulk_ESS | Tail_ESS |
| --- | --- | --- | --- | --- | --- | --- | --- |
| ${\sigma_{\alpha}}_{\mathrm{STATE}}$ | 1.17 | 0.04 | 1.10 | 1.24 | 1 | 1,351 | 3,274 |
| ${\sigma_{\beta}}_{A, STATE}$ | 0.41 | 0.02 | 0.37 | 0.45 | 1 | 6,920 | 9,948 |
| ${\sigma_{\beta}}_{G, STATE}$ | 0.88 | 0.05 | 0.80 | 0.97 | 1 | 5,225 | 8,612 |
| ${cor(\alpha}_{\mathrm{STATE}}, \beta_{A,STATE})$ | -0.20 | 0.08 | -0.31 | -0.08 | 1 | 10,747 | 11,432 |
| ${cor(\alpha}_{\mathrm{STATE}}, \beta_{G,STATE})$ | 0.15 | 0.07 | 0.04 | 0.25 | 1 | 6,824 | 9,629 |
| ${cor(\beta}_{A,STATE}, \beta_{G,STATE})$ | 0.02 | 0.07 | -0.10 | 0.14 | 1 | 2,134 | 4,306 |

Random Effects, Group: macroarea_combination

| Hyperparameter | Estimate | Est.Error | l-89% CI | u-89% CI | Rhat | Bulk_ESS | Tail_ESS |
| --- | --- | --- | --- | --- | --- | --- | --- |
| $\sigma_{\mathrm{AA}}$ | 0.06 | 0.01 | 0.04 | 0.08 | 1 | 5,593 | 9,021 |
| $\beta_{\mathrm{AA}}$ | 0.25 | 0.15 | 0.04 | 0.52 | 1 | 3,033 | 4,720 |
| ${cor(\alpha}_{\mathrm{AA}}, \beta_{\mathrm{AA}})$ | -0.32 | 0.40 | -0.86 | 0.42 | 1 | 10,686 | 9,958 |

table S16.

Summary of fixed and random effects (on log-odds scale) for m2 on the GBI data, using Glottolog areas (sensitivity analysis).

m2 (GBI), Fixed Effects Summary

| Coefficient | Estimate | Est.Error | l-89% CI | u-89% CI | Rhat | Bulk_ESS | Tail_ESS |
| --- | --- | --- | --- | --- | --- | --- | --- |
| $\alpha$ | 0.64 | 0.06 | 0.54 | 0.73 | 1.01 | 773 | 1,914 |
| $\beta_{1}(A)$ | 0.28 | 0.07 | 0.18 | 0.38 | 1.00 | 5,363 | 7,845 |

Random Effects, Group: feature_state

| Hyperparameter | Estimate | Est.Error | l-89% CI | u-89% CI | Rhat | Bulk_ESS | Tail_ESS |
| --- | --- | --- | --- | --- | --- | --- | --- |
| ${\sigma_{\alpha}}_{\mathrm{STATE}}$ | 0.75 | 0.04 | 0.69 | 0.81 | 1 | 1,669 | 3,376 |
| ${\sigma_{\beta}}_{\mathrm{STATE}}$ | 0.39 | 0.03 | 0.35 | 0.44 | 1 | 5,687 | 9,035 |
| ${cor(\alpha}_{\mathrm{STATE}}, \beta_{\mathrm{STATE}})$ | -0.12 | 0.09 | -0.27 | 0.02 | 1 | 6,791 | 9,776 |

Random Effects, Group: macroarea_combination

| Hyperparameter | Estimate | Est.Error | l-89% CI | u-89% CI | Rhat | Bulk_ESS | Tail_ESS |
| --- | --- | --- | --- | --- | --- | --- | --- |
| $\sigma_{\mathrm{AA}}$ | 0.1 | 0.02 | 0.07 | 0.14 | 1 | 5,267 | 8,570 |

table S17.

Summary of fixed and random effects (on log-odds scale) for m2 on the TLI data, using Glottolog areas (sensitivity analysis).

m2 (TLI), Fixed Effects Summary

| Coefficient | Estimate | Est.Error | l-89% CI | u-89% CI | Rhat | Bulk_ESS | Tail_ESS |
| --- | --- | --- | --- | --- | --- | --- | --- |
| $\alpha$ | 1.06 | 0.06 | 0.97 | 1.15 | 1.01 | 807 | 1,980 |
| $\beta_{1}(A)$ | 0.21 | 0.05 | 0.13 | 0.28 | 1.00 | 10,752 | 12,841 |

Random Effects, Group: feature_state

| Hyperparameter | Estimate | Est.Error | l-89% CI | u-89% CI | Rhat | Bulk_ESS | Tail_ESS |
| --- | --- | --- | --- | --- | --- | --- | --- |
| ${\sigma_{\alpha}}_{\mathrm{STATE}}$ | 1.17 | 0.04 | 1.10 | 1.23 | 1 | 1,878 | 3,968 |
| ${\sigma_{\beta}}_{\mathrm{STATE}}$ | 0.43 | 0.02 | 0.40 | 0.47 | 1 | 9,313 | 14,525 |
| ${cor(\alpha}_{\mathrm{STATE}}, \beta_{\mathrm{STATE}})$ | -0.14 | 0.07 | -0.26 | -0.02 | 1 | 14,142 | 15,217 |

Random Effects, Group: macroarea_combination

| Hyperparameter | Estimate | Est.Error | l-89% CI | u-89% CI | Rhat | Bulk_ESS | Tail_ESS |
| --- | --- | --- | --- | --- | --- | --- | --- |
| $\sigma_{\mathrm{AA}}$ | 0.07 | 0.02 | 0.05 | 0.1 | 1 | 7,606 | 12,410 |

table S18.

Summary of fixed and random effects (on log-odds scale) for m4 on the GBI data, using Glottolog areas (sensitivity analysis).

m4 (GBI), Fixed Effects Summary

| Coefficient | Estimate | Est.Error | l-89% CI | u-89% CI | Rhat | Bulk_ESS | Tail_ESS |
| --- | --- | --- | --- | --- | --- | --- | --- |
| $\alpha$ | 0.75 | 0.06 | 0.67 | 0.84 | 1 | 2,758 | 3,842 |
| $\beta_{1}(G)$ | 0.22 | 0.07 | 0.10 | 0.33 | 1 | 5,464 | 4,817 |

Random Effects, Group: broad_pairing_id

| Hyperparameter | Estimate | Est.Error | l-89% CI | u-89% CI | Rhat | Bulk_ESS | Tail_ESS |
| --- | --- | --- | --- | --- | --- | --- | --- |
| $\sigma_{BROAD\_ID}$ | 0.2 | 0.05 | 0.13 | 0.29 | 1 | 3,438 | 4,284 |

Random Effects, Group: broad_pairing_id:pairing_id

| Hyperparameter | Estimate | Est.Error | l-89% CI | u-89% CI | Rhat | Bulk_ESS | Tail_ESS |
| --- | --- | --- | --- | --- | --- | --- | --- |
| $\sigma_{BROAD\_ID/PAIR\_ID}$ | 0.19 | 0.03 | 0.14 | 0.24 | 1 | 3,824 | 5,520 |

Random Effects, Group: feature_state

| Hyperparameter | Estimate | Est.Error | l-89% CI | u-89% CI | Rhat | Bulk_ESS | Tail_ESS |
| --- | --- | --- | --- | --- | --- | --- | --- |
| ${\sigma_{\alpha}}_{\mathrm{STATE}}$ | 0.72 | 0.05 | 0.65 | 0.80 | 1 | 3,860 | 5,624 |
| ${\sigma_{\beta}}_{\mathrm{STATE}}$ | 0.61 | 0.05 | 0.53 | 0.69 | 1 | 3,949 | 5,416 |
| ${cor(\alpha}_{\mathrm{STATE}}, \beta_{\mathrm{STATE}})$ | -0.04 | 0.10 | -0.21 | 0.13 | 1 | 3,686 | 5,243 |

table S19.

Summary of fixed and random effects (on log-odds scale) for m4 on the TLI data, using Glottolog areas (sensitivity analysis).

m4 (TLI), Fixed Effects Summary

| Coefficient | Estimate | Est.Error | l-89% CI | u-89% CI | Rhat | Bulk_ESS | Tail_ESS |
| --- | --- | --- | --- | --- | --- | --- | --- |
| $\alpha$ | 1.17 | 0.05 | 1.09 | 1.26 | 1.01 | 666 | 1,595 |
| $\beta_{1}(G)$ | 0.43 | 0.14 | 0.21 | 0.64 | 1.00 | 2,672 | 3,921 |

Random Effects, Group: broad_pairing_id

| Hyperparameter | Estimate | Est.Error | l-89% CI | u-89% CI | Rhat | Bulk_ESS | Tail_ESS |
| --- | --- | --- | --- | --- | --- | --- | --- |
| $\sigma_{BROAD\_ID}$ | 0.56 | 0.1 | 0.41 | 0.74 | 1 | 2,924 | 4,294 |

Random Effects, Group: broad_pairing_id:pairing_id

| Hyperparameter | Estimate | Est.Error | l-89% CI | u-89% CI | Rhat | Bulk_ESS | Tail_ESS |
| --- | --- | --- | --- | --- | --- | --- | --- |
| $\sigma_{BROAD\_ID/PAIR\_ID}$ | 0.4 | 0.07 | 0.3 | 0.51 | 1 | 2,585 | 4,607 |

Random Effects, Group: feature_state

| Hyperparameter | Estimate | Est.Error | l-89% CI | u-89% CI | Rhat | Bulk_ESS | Tail_ESS |
| --- | --- | --- | --- | --- | --- | --- | --- |
| ${\sigma_{\alpha}}_{\mathrm{STATE}}$ | 1.07 | 0.04 | 1.00 | 1.15 | 1 | 1,499 | 3,117 |
| ${\sigma_{\beta}}_{\mathrm{STATE}}$ | 0.92 | 0.06 | 0.82 | 1.02 | 1 | 3,175 | 4,924 |
| ${cor(\alpha}_{\mathrm{STATE}}, \beta_{\mathrm{STATE}})$ | 0.28 | 0.08 | 0.16 | 0.39 | 1 | 3,440 | 4,881 |

table S20.

Summary of fixed and random effects (on log-odds scale) for m5 on the GBI data, using Glottolog areas (sensitivity analysis).

m5 (GBI), Fixed Effects Summary

| Coefficient | Estimate | Est.Error | l-89% CI | u-89% CI | Rhat | Bulk_ESS | Tail_ESS |
| --- | --- | --- | --- | --- | --- | --- | --- |
| $\alpha$ | 0.66 | 0.05 | 0.58 | 0.74 | 1.01 | 523 | 1,320 |
| $\beta_{1}(G)$ | 0.25 | 0.18 | -0.01 | 0.52 | 1.00 | 2,712 | 3,861 |

Random Effects, Group: broad_pairing_id

| Hyperparameter | Estimate | Est.Error | l-89% CI | u-89% CI | Rhat | Bulk_ESS | Tail_ESS |
| --- | --- | --- | --- | --- | --- | --- | --- |
| $\sigma_{BROAD\_ID}$ | 0.21 | 0.2 | 0.02 | 0.56 | 1 | 2,369 | 4,552 |

Random Effects, Group: broad_pairing_id:pairing_id

| Hyperparameter | Estimate | Est.Error | l-89% CI | u-89% CI | Rhat | Bulk_ESS | Tail_ESS |
| --- | --- | --- | --- | --- | --- | --- | --- |
| $\sigma_{BROAD\_ID/PAIR\_ID}$ | 0.28 | 0.07 | 0.19 | 0.41 | 1 | 4,586 | 7,336 |

Random Effects, Group: feature_state

| Hyperparameter | Estimate | Est.Error | l-89% CI | u-89% CI | Rhat | Bulk_ESS | Tail_ESS |
| --- | --- | --- | --- | --- | --- | --- | --- |
| ${\sigma_{\alpha}}_{\mathrm{STATE}}$ | 0.73 | 0.04 | 0.67 | 0.80 | 1.01 | 1,288 | 3,057 |
| ${\sigma_{\beta}}_{\mathrm{STATE}}$ | 1.25 | 0.08 | 1.13 | 1.38 | 1.00 | 2,400 | 4,731 |
| ${cor(\alpha}_{\mathrm{STATE}}, \beta_{\mathrm{STATE}})$ | 0.07 | 0.08 | -0.07 | 0.20 | 1.00 | 1,467 | 2,584 |

table S21.

Summary of fixed and random effects (on log-odds scale) for m5 on the TLI data, using Glottolog areas (sensitivity analysis).

m5 (TLI), Fixed Effects Summary

| Coefficient | Estimate | Est.Error | l-89% CI | u-89% CI | Rhat | Bulk_ESS | Tail_ESS |
| --- | --- | --- | --- | --- | --- | --- | --- |
| $\alpha$ | 1.05 | 0.05 | 0.96 | 1.13 | 1.01 | 574 | 1,152 |
| $\beta_{1}(G)$ | 0.36 | 0.21 | 0.04 | 0.69 | 1.00 | 7,950 | 10,043 |

Random Effects, Group: broad_pairing_id

| Hyperparameter | Estimate | Est.Error | l-89% CI | u-89% CI | Rhat | Bulk_ESS | Tail_ESS |
| --- | --- | --- | --- | --- | --- | --- | --- |
| $\sigma_{BROAD\_ID}$ | 0.37 | 0.22 | 0.08 | 0.75 | 1 | 4,153 | 4,547 |

Random Effects, Group: broad_pairing_id:pairing_id

| Hyperparameter | Estimate | Est.Error | l-89% CI | u-89% CI | Rhat | Bulk_ESS | Tail_ESS |
| --- | --- | --- | --- | --- | --- | --- | --- |
| $\sigma_{BROAD\_ID/PAIR\_ID}$ | 0.36 | 0.09 | 0.24 | 0.52 | 1 | 5,511 | 10,558 |

Random Effects, Group: feature_state

| Hyperparameter | Estimate | Est.Error | l-89% CI | u-89% CI | Rhat | Bulk_ESS | Tail_ESS |
| --- | --- | --- | --- | --- | --- | --- | --- |
| ${\sigma_{\alpha}}_{\mathrm{STATE}}$ | 1.15 | 0.04 | 1.08 | 1.21 | 1 | 1,474 | 3,655 |
| ${\sigma_{\beta}}_{\mathrm{STATE}}$ | 1.44 | 0.10 | 1.29 | 1.61 | 1 | 6,033 | 11,653 |
| ${cor(\alpha}_{\mathrm{STATE}}, \beta_{\mathrm{STATE}})$ | 0.18 | 0.07 | 0.07 | 0.29 | 1 | 5,560 | 9,349 |

table S22.

Contact effects on state sharing for the main analysis, using AUTOTYP areas. The table lists, for every feature state and every model (indicating the type of contact: genetic contact: all pairs; genetic contact: different area pairs only; genetic contact: same area pairs only; same area (AUTOTYP)), the mean baseline, and the mean difference in state sharing probabilities under contact as compared to the corresponding baseline. As an alternative transformation, it also lists the mean relative risk of these probabilities. Lower and upper 89%-HPDI values are also provided for each baseline, difference and relative risk, as well as a binary variable indicating whether the 89%-HPDI includes zero (for differences) or one (for relative risk).

(external .csv file, tableS22.csv)

table S23.

Contact effects on state sharing for the sensitivity analysis, using Glottolog areas. The table lists, for every feature state and every model (indicating the type of contact: genetic contact: all pairs; genetic contact: different area pairs only; genetic contact: same area pairs only; same area (Glottolog)), the mean baseline, and the mean difference in state sharing probabilities under contact as compared to the corresponding baseline. As an alternative transformation, it also lists the mean relative risk of these probabilities. Lower and upper 89%-HPDI values are also provided for each baseline, difference and relative risk, as well as a binary variable indicating whether the 89%-HPDI includes zero (for differences) or one (for relative risk).

(external .csv file, tableS23.csv)
